# Client-server interfaces enable efficient agent-driven variant calling

**DOI:** 10.64898/2026.06.25.734665

**Authors:** Xian Yu, Zhenxian Zheng, Lei Chen, Zilan Qin, Xinyi Guo, Minggao He, Ruibang Luo

## Abstract

**Background:** Large language model (LLM) agents increasingly automate bioinformatics analyses, but most existing bioinformatics tools were built for standalone use by human experts. An agent driving such a tool must reason about its installation, configuration, and execution from documentation for human, spending many turns, tokens, and tool calls per result. How a method is exposed to an agent can therefore matter as much as the method itself. By designing agentic interfaces for these tools, agent can reduce such overhead and improve the reliability of agent-driven analyses.

**Findings:** To test this design, we re-architected Clair3, a widely used deep-learning-based long-read variant caller, into a client-server system, Clair3-Connect. The client performs all genomics related processing and holds the identifiable data. The server runs only neural-network inference, and the client sends only feature tensors to the server, while sample identifiers and genomic context remain on the client. The client exposes schema-defined agent-facing tools that an agent invokes through single structured calls. On an *APOE* diplotyping task, all 60 agent runs were correct. The agentic tools used 12K tokens in 3 turns, 6.8 to 14 times fewer tokens than the shell-driven baselines (81K-163K tokens), at about a quarter the wall-clock time and far more stably (4% versus 35% token usage variation). Dropping the pileup and phasing stages to keep the client light left SNP F1 within 0.1-0.3 points of standard Clair3 by 50× coverage, while mutual TLS and AES-256-GCM encryption added 7.2% to end-to-end runtime.

**Conclusions:** Recasting an established algorithm as developer-built, agentic tools behind a secure client-server boundary makes it more efficient, reliable, and easier to deploy for an LLM agent than a third-party wrapper, which cannot recover the defaults and conventions only its developers know. Agentic interfaces should be a first-class deliverable of bioinformatics tool development.

## Background

Large language models (LLMs) increasingly operate as agents that plan and execute multi-step tasks with limited human intervention [1]. The clearest demonstrations have come from software engineering, where coding agents resolve real issues in public repositories, navigate codebases, edit across files, and verify changes against test suites [2]. Similar systems are now being explored in the natural sciences, where agents read literature, design analyses or experiments, invoke domain tools, and interpret results [3]. In genomics, agents have begun to automate workflows ranging from general biomedical analysis [4, 5] to specialized tasks such as single-cell analysis [6].

In practice, these agents coordinate their use within multi-step analytical workflows instead of replacing domain algorithms. An agent decomposes a high-level intent into executable steps and calls existing software, databases, or services to perform each step [7]. The interface through which an algorithm is exposed therefore becomes part of the computational system. It determines what the agent can call, what information it receives, how errors are represented, and how much reasoning is required to interpret the result.

This creates a practical mismatch with current bioinformatics ecosystem, where most widely used genomics tools remain standalone command-line programs designed for expert human users. Bioinformatics tools such as sequence aligners [8–10], de novo assemblers[11, 12], structural-variant callers[13–15], and small-variant callers[16, 17] typically communicate through files, logs, and conventions that are clear to experienced users but not always explicit to an agent. A shell-based agent must reason about installation, configuration, file paths, indexes, command flags, and intermediate outputs. It must also infer tool semantics from unstructured text, headers, and final result files. These extra steps consume tokens and turn, and they introduce opportunities for unreliable parsing or unsupported assumptions.

Agent-oriented interfaces can reduce this burden by making tool behavior explicit. An agentic tool interface can expose operations, input schemas, structured outputs, error states, and default behavior directly, rather than forcing an agent to reconstruct them from command-line output. The wider software ecosystem is moving toward standardized client-server tool interfaces, including protocols such as the Model Context Protocol [18]. In biology, early evidence from agent-designed database access suggests that replacing human-facing interfaces with deterministic, agent-facing ones can improve agent accuracy and reproducibility [19]. Bioinformatics algorithms, however, are still most commonly delivered as standalone tools rather than as semantically explicit services.

Moreover, genomics also imposes constraints that make this interface problem more than a software-engineering issue. Sequencing data can contain sensitive or potentially identifying information, and large neural-network-based workflows may require hardware or runtime dependencies that are not available on every client machine. Moving full sequencing files to a remote environment can be undesirable, but local execution can be expensive or difficult to standardize. An agent-facing genomics system therefore needs to address three requirements at once: it must expose the algorithm in a form that agents can use reliably, keep sample-specific context under client control, and allow compute-intensive inference to be shared when appropriate.

To examine this design in a mature genomics tool, we use Clair3 as a representative example and re-architect it into Clair3-Connect, a client-server system for agent-facing variant calling. Clair3-Connect keeps sample-specific genomic processing on the client and exposes neural-network inference as a stateless server-side service. The agent interacts with the caller through structured tool calls rather than by scripting a command-line program. We evaluate this design by comparing agentic tool access with shell-based agent access, measuring the cost and variability of agent-driven variant calling, quantifying the accuracy trade-off introduced by a lightweight client, and estimating the overhead of secure transport. Together, these experiments test whether a mature genomics algorithm can be delivered as agent-facing infrastructure that reduces the interaction overhead for agents while preserving variant-calling performance and providing secure client-server execution.

## Findings

### Overview of the system

Clair3-Connect re-architects Clair3 around a single computational boundary: sample-specific genomic processing remains on the client, whereas neural-network inference is exposed as a stateless server-side service (Figure 1). The client reads the alignments and reference, selects variant candidate sites, constructs Clair3 feature tensors for these sites, and decodes returned probabilities into variant calls. The server runs only the neural network, computing tensors to per-site probabilities without receiving the surrounding genomic or file-system context. Thus, the exchanged data are limited to feature tensors and request identifiers than do not encode sample, genomic or file system context. This keeps sample-specific context on the client and reduces exposure of sensitive or potentially identifying information when inference is performed on shared or remote hardware.

**Figure 1.**
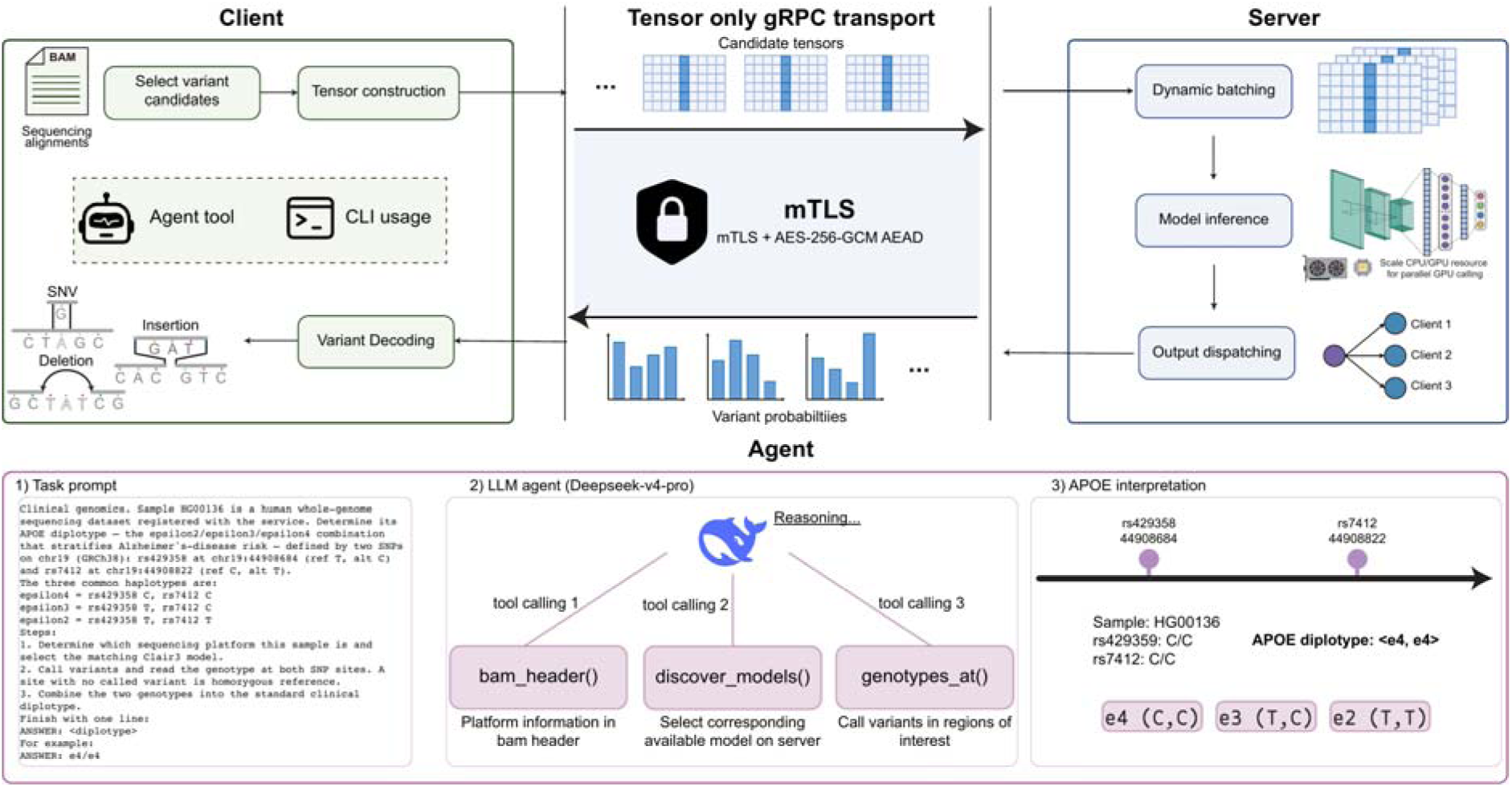
Overview of Clair3-Connect. The top panel shows the client-server architecture. The client (`clair3-connect`) reads the BAM/CRAM alignments, selects candidate variant sites, and builds the full-alignment feature tensors, exposing these operations both as a command-line program and as typed agent tools, and decodes the returned probabilities into SNP and indel calls. Only opaque candidate tensors cross the gRPC transport, secured by mutual TLS and per-payload AES-256-GCM authenticated encryption. The server (`clair3-connect-server`) performs dynamic batching and neural-network inference on CPU or GPU and dispatches the per-site probabilities back, so one server serves many clients. The bottom panel shows an LLM agent driving the typed tools on the *APOE* diplotyping task. From the task prompt, the agent (`deepseek-v4-pro`) reasons and issues three structured calls, `bam_header` to read the sequencing platform from the BAM header, `discover_models` to select the matching model, and `genotypes_at` to call variants at the two sites, then combines the rs429358 and rs7412 genotypes into the clinical diplotype (sample HG00136, ε4/ε4).

The same boundary separates GPU-intensive inference from the client-side genomics workflow. No local GPU is required for the client side, and the server can batch and accelerate inference across requests. The structured client interface also makes Clair3-Connect directly callable by applications and LLM agents. Instead of scripting a command-line program and parsing unstructured files, an agent invokes structured tool calls with defined inputs, outputs and error states. The following sections evaluate the effect of this design on agent cost, variant-calling accuracy, and secure transport overhead. Transport, runtime, and interface details are given in the Methods section.

### Driving Clair3-Connect from an LLM agent

We compared three ways of exposing Clair3 to an LLM agent while holding the model and task fixed (Methods). The RawBinary condition only gave the agent shell access to the stock Clair3 program [17]. The Skills condition added a skill document provided by Clair3’s developers to the same shell environment. The Tools condition exposed only the Clair3-Connect agentic tool interface. Each condition was run in ten independent repeats per sample. All 60 runs returned the correct *APOE* diplotype; only the cost and predictability differed (Figure 2). The prompts and per-repeat metrics are reported in Supplementary Note S1, with one representative transcript in Supplementary Note S4.

**Figure 2.**
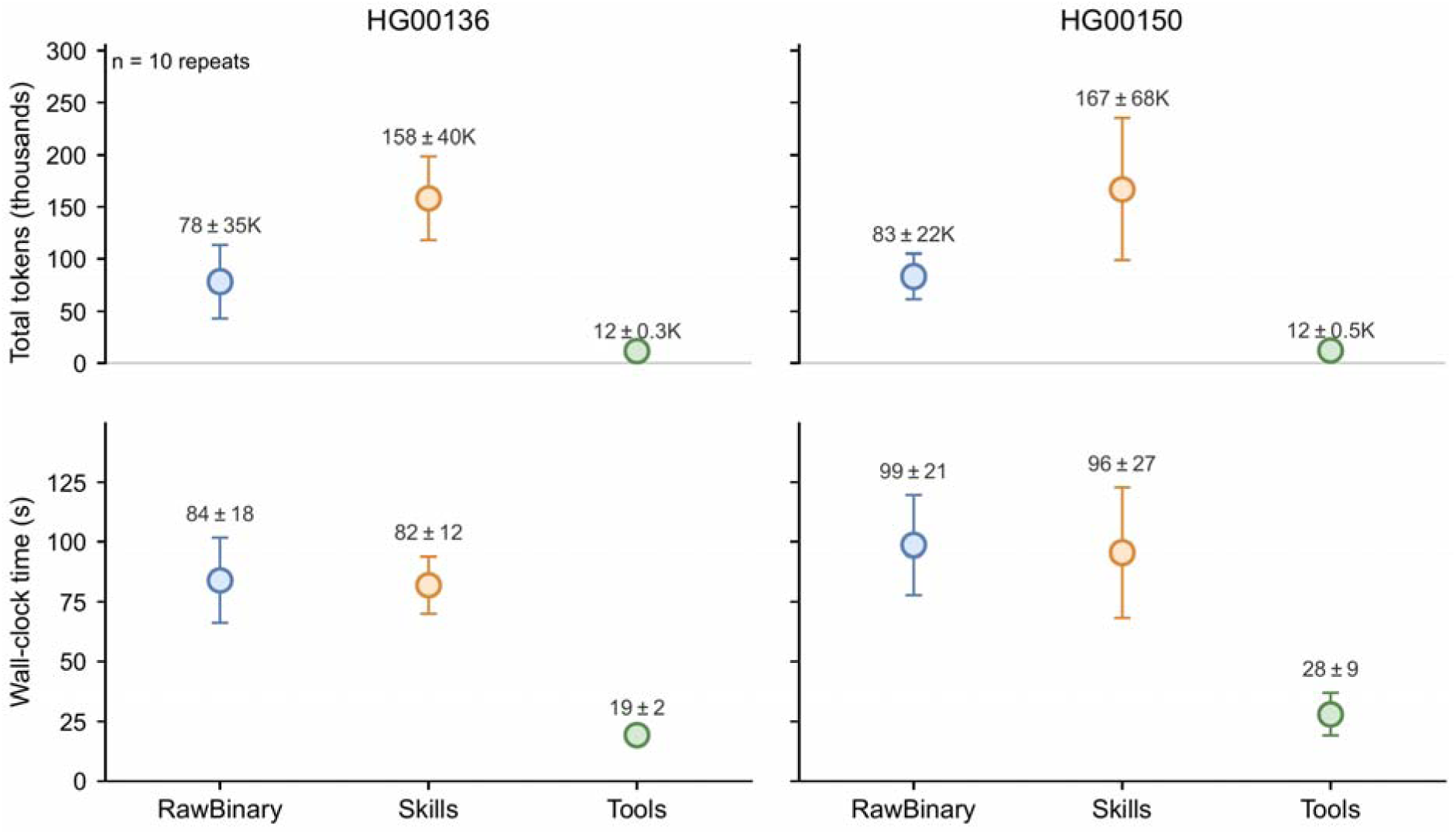
Cost of the *APOE* diplotyping task under the three deployment surfaces, over ten independent repeats per condition. Columns are the two whole-genome samples (HG00136, HG00150); the top row plots mean total tokens and the bottom row mean wall-clock time, each condition shown as a marker with a whisker of ±1 standard deviation across repeats. All conditions used the same agent model (`deepseek-v4-pro`) and returned the correct diplotype in every run. The typed Tools interface is both far affordable and consistent, its token usage varying by 4% across repeats versus about 35% for the shell conditions. Per-repeat metrics are in Supplementary Note S1 and one representative round’s full transcript in Note S4.

The way Clair3 is exposed affected the cost of driving it, with the Tools condition interface being the most efficient. The RawBinary condition used 81K tokens over 10 turns on average, and Skills used 163K tokens over 7.5 turns. The Tools condition reached the answer in 12K tokens and 3 turns, corresponding to 6.8-fold fewer tokens than the RawBinary condition and 14-fold fewer tokens than the Skills condition, with approximately one quarter of the wall-clock time. The Skills condition was most expensive because the skill document was re-injected on every turn. In both shell-based settings, the agent also had to inspect bulky unstructured outputs, including BAM headers, region dumps, command-line logs, and VCF text.

The agentic tool interface also reduced run-to-run variability. Token usage varied by only 4% across Tools repeats, compared with approximately 35% in both shell-based conditions (Figure 2). Agentic tool calls return compact, task-specific results and restrict the agent to a small action space. Shell access, by contrast, leaves the agent to decide which commands to run, which outputs to inspect, and how much intermediate evidence to re-check. Thus, the interface affects not only the cost of an agentic workflow, but also its run-to-run stability.

The HG00136 sample illustrates why agent-facing tools should encode algorithm-specific semantics. HG00136 is ε4/ε4, and the rs7412 site is homozygous for the reference allele. Under Clair3’s default behavior, such sites are absent from the VCF unless reference calls are requested with `--print_ref_calls`. None of the shell-based runs used this option, and the skill document did not mention it. The shell arms reached the correct answer only because the prompt supplied a benchmark-specific convention: for the two predefined APOE loci, an absent record in Clair3’s variant-only VCF should be treated as reference for the purpose of deriving the diplotype. This was not intended as a general rule for VCF interpretation. In general, the absence of a VCF record should not be interpreted as evidence for a homozygous-reference genotype, because missing records may also reflect low confidence, insufficient coverage, filtering, or non-callable regions.

The Tools arm did not require this convention in the prompt. The developer-built genotype query returned an explicit `0/0` genotype for positions absent from the VCF (Supplementary Note S4). This example shows a limitation of directly wrapping command line tools for agentic use: defaults, flags, and missing-record conventions are often part of the algorithm’s operational semantics, but they are not reliably recoverable from final output files alone. Exposing them through typed tools allows the agent to use the caller with less parsing, less redundant verification, and fewer opportunities for unsupported assumptions.

Together, these results show that a client-server typed-tool interface improves both the efficiency and robustness of agent-driven variant calling. They also suggest that agent-facing infrastructure should be treated as part of bioinformatics algorithm development, not as a thin wrapper added after the fact.

### System performance and implementation trade-offs

To minimize client-side computation, Clair3-Connect deviates from the standard Clair3 pipeline. Standard Clair3 first applies a pileup model to all candidate sites, phases and haplotags the reads with WhatsHap[20] or LongPhase[21], then runs the full-alignment model on the remaining sites and merges the outputs. Clair3-Connect builds only the full-alignment tensor and calls from it directly, removing the pileup and phasing stages.

The accuracy gap introduced by removing pileup and phasing was concentrated at low coverage and in noisier read data and narrowed to a small residual difference at higher coverage or with higher-accuracy reads. For SNPs on chromosome 20, the full-alignment-only configuration achieved SNP F1 scores 9.5 and 12.5 percentage points lower than standard Clair3 on Illumina and ONT data, respectively (Figure 3). The ONT accuracy gap is driven by precision, which falls to 69.5% from 90.1% as the model retains more false positives where reads are sparse. The gap then narrows with depth, falling below one F1 point by 30× and to 0.1-0.3 points by 50× on both platforms. On PacBio HiFi it stays below 0.1 points at every depth, because per-read accuracy reducing the reliance on phasing. Indels show the same pattern with larger, more persistent gaps, because indel calling is more sensitive to read-level alignment noise and benefits more from phasing and haplotagging. The residual gap is largest on ONT, where the full-alignment-only client loses about 8 F1 points at 10× and still 4 F1 points by 50× (Figure 3, bottom row). The complete benchmark for both chromosomes is in Supplementary Note S2. To conclude, at moderate-to-high depths typical of many whole-genome sequencing applications, Clair3-Connect retains most of standard Clair3’s accuracy while reducing the computation that must be performed locally.

**Figure 3.**
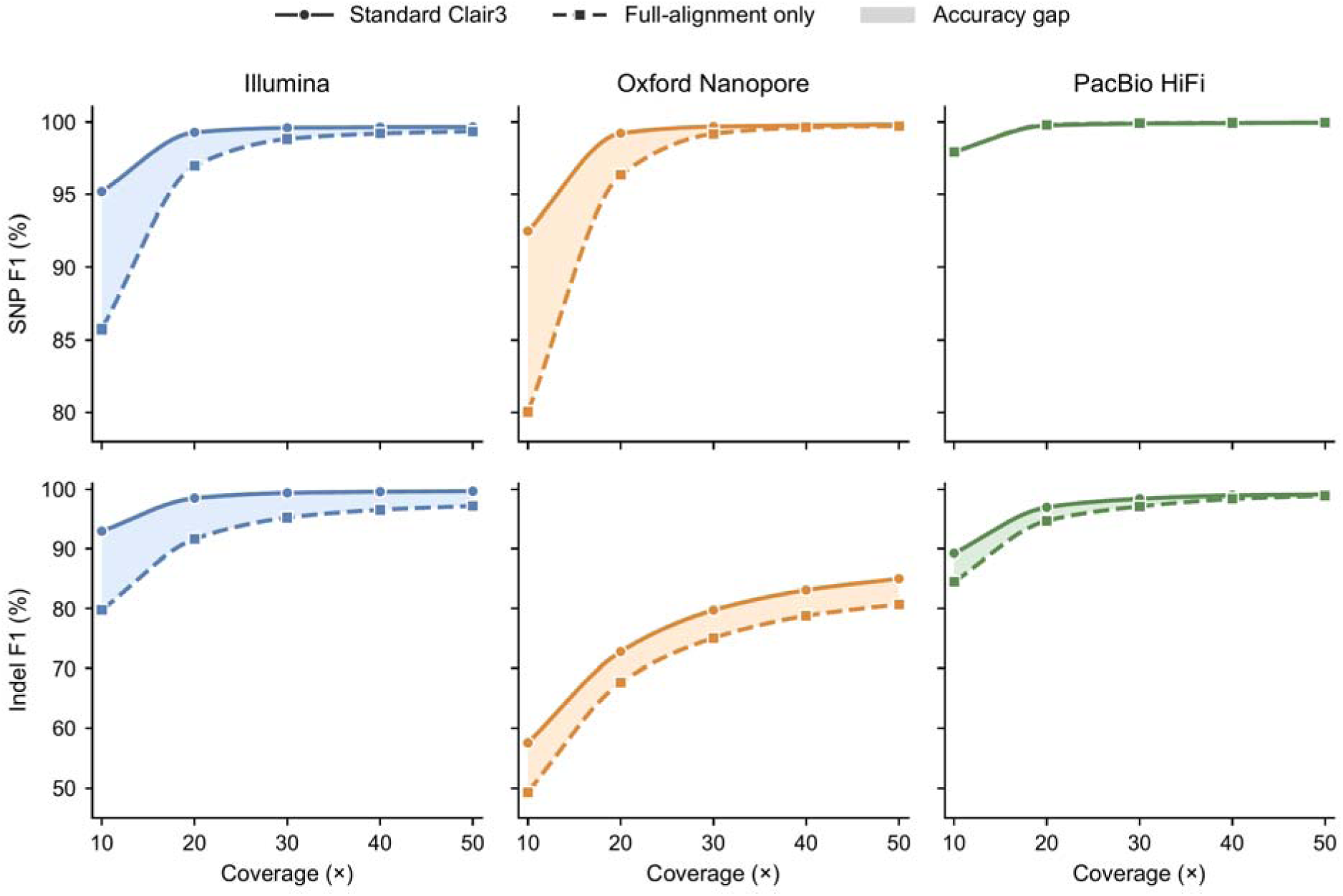
Variant-calling accuracy on chromosome 20 as a function of sequencing depth, for standard Clair3 (full pipeline, solid) versus the full-alignment-only client used by Clair3-Connect (pileup and phasing removed, dashed), on Illumina, Oxford Nanopore, and PacBio HiFi reads (one color per platform). The top row is SNPs and the bottom row indels; the shaded band between the two curves marks the accuracy gap. Removing pileup and phasing trades accuracy for far less local computation. SNP gaps are largest at low depth and on the noisier platforms and essentially vanish by 30-50× (and at all depths on HiFi); indel gaps are larger and more persistent, staying a few F1 points wide on ONT even at 50×, because phasing helps indels most. Chromosome 1 and the complete titration are in the Supplementary Materials (Note S2).

The light client also has a small local footprint (Table 1). It needs no GPU and no deep-learning runtime, completes a chromosome in tens of seconds to a few minutes, and peaks between about 4 and 11 GB of RAM using 8 threads, scaling with chromosome size and with platform error rate.

**Table 1.** Runtime and peak memory of the full-alignment-only Clair3-Connect client. Wall-clock time and peak resident set size (RSS) for calling variants across each whole chromosome of sample HG003 at 30× coverage, on Illumina, Oxford Nanopore, and PacBio HiFi reads. Values are mean ± standard deviation over ten repeats.

| Platform | Chromosome 1 |  | Chromosome 20 |  |
| --- | --- | --- | --- | --- |
|  | Wall time (s) | Peak memory (GB) | Wall time (s) | Peak memory (GB) |
| Illumina | 74.7 ± 2.9 | 5.01 ± 0.30 | 25.1 ± 1.1 | 3.83 ± 0.10 |
| Oxford Nanopore | 210.6 ± 21.1 | 10.66 ± 1.39 | 43.4 ± 0.3 | 7.15 ± 0.28 |
| PacBio HiFi | 70.5 ± 6.1 | 6.86 ± 0.14 | 22.1 ± 0.9 | 4.76 ± 0.13 |

### Transport security adds measurable but modest end-to-end overhead

A shared or remote inference server raises the question of data exposure in transit. Clair3-Connect addresses this risk at two levels. First, the client-server boundary limits what can leave the client. The server receives feature tensors, but not explicit sample identifiers, file paths, genomic coordinates, or other contextual metadata. Second, Clair3-Connect can protect the transmitted tensors with two standard mechanisms. Mutual TLS (mTLS) authenticates both endpoints and encrypts the channel. Per-payload AES-256-GCM encryption (AEAD) encrypts each tensor and binds it to its model, shape, and content. These mechanisms provide standard channel confidentiality, endpoint authentication, and payload integrity. Here, we measure the cost of applying them to Clair3-Connect’s tensor-based workflow.

We first measured the transport layers in isolation. The full-alignment tensor used in this benchmark had low byte-level entropy (0.82 bits/byte) and was highly compressible: ZSTD-1 reduced the 752 KB ONT tensor to ∼22 KB, corresponding to a 34.0× reduction and ∼3% of the original size. This benchmark used the full 752 KB tensor without compression and made inference the only server-side work, placing the tensor on the request critical path. Under plaintext transport, per-request latency was 13.4 ms. Latency increased to 20.8 ms with AEAD and to 21.6 ms with mTLS+AEAD (Figure 4a). The absolute AEAD cost was stable at approximately 7.4 ms.

**Figure 4.**
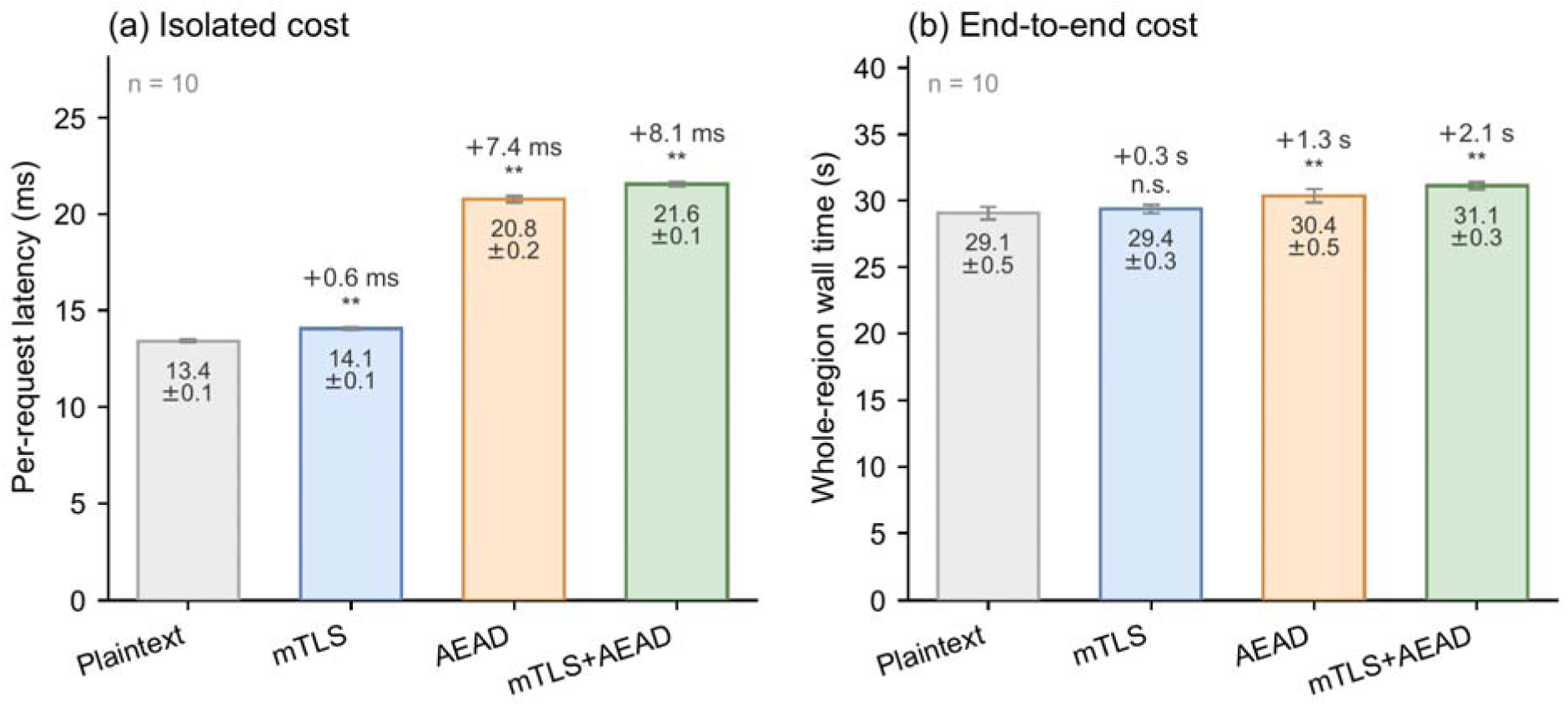
Transport-security cost (in absolute units), measured on an Oxford Nanopore whole-genome run (sample HG003, chromosome 20). **(a)** Isolated per-request latency (a deliberate worst case in which inference is the only work and the full 752 KB payload is encrypted on the critical path) and **(b)** end-to-end wall-clock time of a real full-alignment-only run (the deployment-relevant cost), each for an unencrypted (plaintext) channel and the two security layers. Bars are means over *n* = 10 paired rounds, whiskers are 95% confidence intervals, and each secured bar is labelled with its absolute increase over plaintext and its paired Wilcoxon significance (`**` *p* < 0.01, `n.s.` not significant). The per-request payload-encryption cost (≈7.4 ms) is largely absorbed by the CPU-bound candidate scan, so the whole-region cost of the secure default is only about 2 s. The confidentiality, mTLS access-control, and AEAD integrity checks are in the Supplementary Materials (Note S3).

The deployment-relevant cost was smaller in the full workflow. In a full-alignment-only run, client-side candidate scanning dominated runtime and absorbed much of the transport overhead. Enabling mTLS+AEAD increased end-to-end wall time by 7.2%, or about 2 s on a 29 s run. AEAD alone added 4.5% wall time, whereas mTLS alone added a non-significant 1.1% (Figure 4b; both AEAD-containing comparisons, *p* < 0.01). Transport security is therefore measurable in this workload, but it does not dominate the overall runtime.

## Discussion

In this study, we use Clair3-Connect to show that the design of a bioinformatics tool interface can strongly affect the efficiency and reliability of LLM-agent tool use. We demonstrated that the agentic interface made agent-driven variant calling more efficient and more stable than shell-based access, not by changing the biological task or the underlying model, but by changing what the agent could see and how it could act. This suggests that agent-facing interfaces should be treated as part of bioinformatics tool development, not as optional add-ons developed after the tool is released.

The need for such interfaces is especially clear in mature bioinformatics tools. Correct use often depends on defaults, flags, and output conventions that are familiar to developers but difficult to infer from final files alone. A command-line wrapper can expose the executable, but it cannot reliably by itself expose the algorithm’s operational semantics. Agentic tools allow developers to make these operational details explicit, including structured outputs, explicit defaults, error states, version information, and provenance. For agent-driven workflows, this reduces the need for fragile parsing and unsupported interpretation of logs, headers, and intermediate files.

Clair3-Connect also reflects the constraints of adapting existing standalone tools to this mode of use. Most bioinformatics software was designed for expert users and file-based workflows, not for agent orchestration through typed calls. In Clair3-Connect, keeping the client lightweight required simplifying the standard Clair3 workflow, and this introduced accuracy costs in the regimes where pileup and phasing are most useful. These costs define the coverage, read-quality, and variant-type settings in which the current design is most appropriate rather than a universal replacement for the original caller. Future implementations could recover part of this gap through lighter client-side filters, server-side preprocessing, models trained for lower-depth settings, or additional structured intermediate outputs.

More broadly, reliable agent-driven bioinformatics will require conventions that extend beyond a single caller. Aligners, assemblers, variant callers, structural-variant tools, and reference databases would all benefit from agent-facing interface schemas that specify inputs, outputs, defaults, provenance, and failure modes. Clair3-Connect provides one example of this direction: a validated algorithm delivered not only as a standalone program, but also as a semantically explicit interface that agents can call and compose with less ambiguity.

## Methods

### System architecture and inference workflow

Clair3-Connect is implemented in Rust and comprises three components, a client that performs the genomic stages of variant calling, a stateless server that runs the neural network, and the gRPC transport layer between them. The client takes an indexed BAM or CRAM alignment, the reference genome, and the regions to call, supplied as a single interval, a BED file, or a list of intervals. Within each region it selects candidate sites using Clair3’s default allele-frequency criteria, with minimum allele frequency of 0.08 for SNPs and 0.15 for indels at a minimum read depth of 2. We disabled the pileup model and the read phasing module, so the tensor’s haplotype channel in the resulting tensor carries no haplotype assignment. For every candidate the client builds the full-alignment feature tensor defined by Clair3, an int8 tensor with a per-platform read depth of 89 rows for Oxford Nanopore and 55 for PacBio HiFi and Illumina, across 33 flanking positions and 8 feature channels. This full-alignment-only configuration is the default configuration, and its accuracy relative to standard Clair3 is characterized in the Findings. The client sends the candidate tensors to the server in batches of 32, receives the per-site genotype probabilities, and decodes them into SNP and indel calls using Clair3’s output logic. It writes the calls to a Clair3-compatible VCF as a self-contained command-line caller or exposes them to an application or LLM agent through the typed tools described below.

The server performs only inference. It loads one or more Clair3 models as TorchScript modules using libtorch, on CPU or a specified GPU, and processes each request by running the selected model on a batch of feature tensors and returning the per-site probabilities. A dynamic batcher coalesces concurrent requestsuntil either 64 tensors are available or 4 ms has elapsed, so one server can accelerate inference for many clients. The server is stateless across requests apart from the loaded models. Its request schema has no fields for file paths, genomic coordinates, or sample or read names. Each request carries only a request identifier that does not encode sample, genomic, or file-system context, together with the tensor’s model identifier, data type, and shape. Thus, the server receives no explicit sample identifiers or genomic coordinates and can run on shared or remote hardware.

The same client operations are also exposed as typed tools, callable as OpenAI-style function tools or through the Model Context Protocol[18]. Through MCP, the agent can query genotypes at specified positions, inspect BAM metadata such as sequencing platform, and discover available server-side models, all through structured inputs and compact structured outputs rather than shell commands or parsed text files. Each tool returns a compact structured result; the full input and output schemas are in Supplementary Note S1.

### Evaluation for agent inference

To measure how the agent-facing interface affects an LLM agent, we fixed the agent model and the task and varied only the interface through which Clair3 was exposed. Three conditions were compared. The RawBinary condition gave the agent a single `run_shell` tool, which runs one bash command and returns its exit code and output, together with the path to the standard Clair3 command-line program. The Skills condition added the expert-authored Clair skill document (https://github.com/HKU-BAL/Clair-skills) to the system prompt of that same shell environment, with the skill’s reference notes and command templates readable through `run_shell`. The Tools condition removed the shell and exposed only the typed Clair3-Connect tools. The shell conditions were given the sample and reference locations and an invocation template for stock Clair3, whereas the Tools condition referenced datasets by sample name alone.

We used APOE diplotyping as a constrained but clinically interpretable task for evaluating agent-driven use of Clair3-Connect. *APOE* diplotyping determines the ε2/ε3/ε4 diplotype from the two SNPs rs429358 (chr19:44908684) and rs7412 (chr19:44908822) on two whole-genome samples, HG00136 and HG00150. Every condition used the same task prompt and the same agent model, `deepseek-v4-pro`. The prompt gave the two SNP coordinates and stated that, for this predefined task, a site with no called variant is homozygous reference, but it withheld each sample’s sequencing platform. To keep the comparison fair, every condition had to read the platform from the BAM header and select the matching model. The shell conditions did this with `samtools view -H`, and the Tools condition, which was not given a sample-metadata shortcut, had to call `bam_header`. The environment for this clinical task excluded the unrelated benchmark data.

The agent ran as an OpenAI-style function-calling loop. At each turn the model, decoding at temperature 0, either issued tool calls, whose results were appended to the conversation, or returned a final answer. The loop ended when the model stopped calling tools. A deliberately high turn cap of 50 and a 30-minute wall-clock budget let every condition complete without premature truncation, so the comparison reflects the cost of each interface. If a run reached the budget mid-analysis, a single follow-up request with tools disabled elicited its best conclusion from the evidence already gathered; this step was applied identically to every condition and added no task information. We ran each condition on each sample in ten independent repeats, recording the total tokens, wall-clock time, reasoning turns, and tool calls per run and reporting the mean and standard deviation across repeats. Answers were taken from a final `ANSWER:` line, with diplotypes scored as unordered allele sets after normalizing ε to e. The system prompts, task prompt, and per-repeat metrics are in Supplementary Note S1, and one representative round’s full transcript in Note S4.

### Variant-calling benchmark

We compared two configurations on sample HG003, which is held out from Clair3’s training data. Standard Clair3 was the official v2.0.1 GPU release, running its full pipeline of a pileup model, read phasing and haplotagging, and a full-alignment model over the remaining sites. The other configuration was the full-alignment-only client deployed in Clair3-Connect, run as the client-server system with the server performing inference on a GPU. Both used the same Clair3 model checkpoints, Clair3-Connect serving TorchScript exports of the checkpoints that standard Clair3 loads, so the comparison isolates the effect of the simplified pipeline rather than a difference in models. We used three platforms on GRCh38, Illumina NovaSeq, Oxford Nanopore, and PacBio HiFi Revio. From each native high-depth alignment we extracted chromosomes 1 and 20 and built a coverage titration at 10, 20, 30, 40, and 50× by subsampling reads. Calls from each configuration were compared against the GIAB v4.2.1 benchmark for HG003[22] with `hap.py` [23], restricted to each chromosome and the benchmark’s high-confidence regions. We report F1, recall, and precision for SNPs and indels. Chromosome 20 SNPs and indels are summarized in Figure 3, and the complete titration for both chromosomes is tabulated in Supplementary Note S2.

We also profiled the computational cost of the full-alignment-only client. For each platform, we called variants on chromosomes 1 and 20 at 30× coverage in ten repeats and recorded wall-clock time and peak memory usage. The client runs only the genomic stages, offloading inference to the server, whose GPU and host memory we recorded separately. The means are reported in Table 1 and the per-repeat values in Supplementary Note S2.

### Transport security

All client-server communication uses gRPC, secured by two independent layers that are each optional. Mutual TLS encrypts the channel and requires both endpoints to present a certificate chained to a configured authority. This gives channel confidentiality and integrity and admits only authenticated clients. Before payload encryption, tensor payloads can optionally be compressed with Zstandard (ZSTD).The second layer applies authenticated encryption to each tensor payload, independently of the channel. Each tensor is sealed with AES-256-GCM under a fresh 96-bit random nonce and a 16-byte authentication tag. The additional authenticated data covers the request’s model identifier, data type, tensor shape, and payload hash. This binds each ciphertext to these request metadata fields and prevents it from being accepted into a valid tensor under a different model or shape. Because the seal is end-to-end, a TLS-terminating intermediary that decrypts the channel still sees only the encrypted payload. Key management, including nonce limits and rotation, is documented with the released source code.

We measured the cost of these layers across four configurations, an unencrypted channel (plaintext), mTLS alone, AEAD alone, and the secure default of mTLS and AEAD together. To control for host drift, all four were served on separate ports simultaneously, given one warm-up run, and measured interleaved within each round. We report the mean and 95% confidence interval over ten rounds and test each configuration against plaintext with a paired Wilcoxon signed-rank test on the per-round differences. All measurements used sample HG003 on Oxford Nanopore chromosome 20, on a 128-core host with one RTX 4090, over loopback, so the reported costs are pure compute and exclude external network latency. The per-request payload was an int8 full-alignment tensor (≈752 KB).

Cost was measured in two settings. The isolated cost used a pure-inference generator that performs no candidate scan, placing the full payload encryption on the request critical path; for it we report per-request latency. The end-to-end cost was the wall-clock time of a real full-alignment-only run over chromosome 20:1-8 Mb, which is bound by the CPU candidate scan. We also verified the layers’ guarantees. For confidentiality we confirmed on a real full-alignment tensor that the AES-256-GCM ciphertext showed no detectable byte-level structure of the plaintext under our tests. Authentication was tested by presenting a valid client certificate, no certificate, and a certificate from an unrelated authority. Integrity was tested with a transport unit-test suite that tampers with the ciphertext, the additional authenticated data, the key, and the nonce length. The full statistics and outcomes are in Supplementary Note S3.

## Availability of supporting source code and requirements

- **Project name:** Clair3-Connect
- **Project home page:** https://github.com/HKU-BAL/Clair3-Connect
- **Operating system(s):** Linux
- **Programming language:** Rust (variant-calling engine); Python (agent interface and evaluation harness)
- **Other requirements:** Rust 1.85 or later and libtorch 2.5 to build the engine (a CUDA-capable GPU is optional and used only by the inference server); a TorchScript-exported Clair3 model; Python 3.11 or later for the agent and evaluation code; hap.py and samtools for the accuracy benchmark

## Data availability

All data supporting the results of this article are provided within the article and its Supplementary Materials. The accuracy benchmark uses the publicly available HG003 sequencing data and the GIAB v4.2.1 benchmark set[22]. The specific download paths for the sequencing samples, the reference genome, and the truth sets, together with the access paths for the tools and models used, are provided in the Supplementary Materials.

## List of abbreviations

AEAD: authenticated encryption with associated data
AES-256-GCM: Advanced Encryption Standard (256-bit key) in Galois/Counter Mode
BAM: Binary Alignment Map
CRAM: Compressed Reference-oriented Alignment Map
CV: coefficient of variation
GIAB: Genome in a Bottle
GPU: graphics processing unit
gRPC: gRPC Remote Procedure Call
HiFi: high-fidelity (PacBio) reads
LLM: large language model
MCP: Model Context Protocol
mTLS: mutual Transport Layer Security
ONT: Oxford Nanopore Technologies
PHI: protected health information
RPC: remote procedure call
SNP: single-nucleotide polymorphism
SV: structural variant
TLS: Transport Layer Security
VCF: Variant Call Format
WGS: whole-genome sequencing

## Authors’ contributions

R.L. conceived and supervised the study. X.Y. implemented the software. Z.Z. developed Clair3. X.Y. and Z.Z. wrote the paper. L.C., Z.Q., X.G., M.H., and R.L. reviewed and revised the manuscript.

## References

1. Ren, S., et al., Towards scientific intelligence: A survey of llm-based scientific agents. arXiv preprint arXiv:2503.24047, 2025.

2. Jimenez, C.E., et al. Swe-bench: Can language models resolve real-world github issues? In International Conference on Learning Representations. 2024.

3. Gridach, M., et al., Agentic ai for scientific discovery: A survey of progress, challenges, and future directions. arXiv preprint arXiv:2503.08979, 2025.

4. Huang, K., et al., Biomni: A general-purpose biomedical ai agent. biorxiv, 2025.

5. Su, H., W. Long, and Y. Zhang, BioMaster: Multi-agent system for automated bioinformatics analysis workflow. bioRxiv, 2025: p. 2025.01. 23.634608.

6. Xiao, Y., et al., Cellagent: An llm-driven multi-agent framework for automated single-cell data analysis. arXiv preprint arXiv:2407.09811, 2024.

7. Dip, S.A., et al., Large language model agents for biological intelligence across genomics, proteomics, spatial biology, and biomedicine. Briefings in Bioinformatics, 2026. 27(2): p. bbag110.

8. Li, H., Minimap2: pairwise alignment for nucleotide sequences. Bioinformatics, 2018. 34(18): p. 3094–3100.

9. Zhang, P., et al., HAlign-G: rapid and low-memory multiple-genome aligner for large-scale closely related genomes. Genome Biology, 2025. 26(1): p. 406.

10. Gao, Y., et al., abPOA: an SIMD-based C library for fast partial order alignment using adaptive band. Bioinformatics, 2021. 37(15): p. 2209–2211.

11. Cheng, H., et al., Haplotype-resolved de novo assembly using phased assembly graphs with hifiasm. Nature methods, 2021. 18(2): p. 170–175.

12. Kolmogorov, M., et al., Assembly of long, error-prone reads using repeat graphs. Nature biotechnology, 2019. 37(5): p. 540–546.

13. Jiang, T., et al., cuteFC: regenotyping structural variants through an accurate and efficient force-calling method. Genome biology, 2025. 26(1): p. 166.

14. Jiang, T., et al., Long-read-based human genomic structural variation detection with cuteSV. Genome biology, 2020. 21(1): p. 189.

15. Smolka, M., et al., Detection of mosaic and population-level structural variants with Sniffles2. Nature biotechnology, 2024. 42(10): p. 1571–1580.

16. Poplin, R., et al., A universal SNP and small-indel variant caller using deep neural networks. Nature biotechnology, 2018. 36(10): p. 983–987.

17. Zheng, Z., et al., Symphonizing pileup and full-alignment for deep learning-based long-read variant calling. Nature computational science, 2022. 2(12): p. 797–803.

18. Hou, X., et al., Model context protocol (mcp): Landscape, security threats, and future research directions. ACM Transactions on Software Engineering and Methodology, 2025.

19. Luebbert, L., et al., Paving the way for agents in biology. 2026.

20. Martin, Mb., et al., WhatsHap: fast and accurate read-based phasing. BioRxiv, 2016: p. 085050.

21. Lin, J.-H., et al., LongPhase: an ultra-fast chromosome-scale phasing algorithm for small and large variants. Bioinformatics, 2022. 38(7): p. 1816–1822.

22. Zook, J.M., et al., An open resource for accurately benchmarking small variant and reference calls. Nature biotechnology, 2019. 37(5): p. 561–566.

23. Krusche, P., et al., Best practices for benchmarking germline small-variant calls in human genomes. Nature biotechnology, 2019. 37(5): p. 555–560.

