## Supplementary materials for "Client-server interfaces enable efficient agent-driven variant calling"

### S1 Agent-based evaluation

This note collects the raw inputs and outputs of the agent-based evaluation: the three deployment conditions and their system prompts (§S1.1), the task prompt (§S1.2), the per-run metrics (§S1.3); the complete agent transcripts are reproduced in Supplementary Note S4.

#### S1.1 Experimental setup: three deployment conditions

The three conditions (Table S1) all drive the same Clair3 engine on the same task; they differ only in what the agent is handed.

**Table S1.** The three deployment conditions. All conditions use the identical agent model and task prompt; only the interface to Clair3 differs.

| Condition | What the agent is given | What the agent must do |
|---|---|---|
| **RawBinary** | A shell (`run_shell`) + the stock Clair3 CLI | Directly invoke Clair3: determine each sample's platform from its BAM header, choose the matching model, build a region BED, assemble and run the command, and parse the output VCF by hand. |
| **Skills** | A shell (`run_shell`) + our Clair-series skill document | Drive the same shell, but guided by our `clair-variant-caller` skill (model catalogs, platform/model selection, ready-to-run command templates, result analysis). |
| **Tools** | Our packaged Clair3-Connect tools only | Reference data by `(sample, platform)` and call typed tools (`bam_header`, `discover_models`, `genotypes_at`); never handle file paths or parse VCFs. |

**(a) RawBinary.** System prompt (`run_shell` + the stock Clair3 CLI):

```
You are a genomics analysis assistant on a Linux host with a single tool: run_shell,
    and a pointer to the stock (monolithic) Clair3 CLI.
Local execution environment (NO Docker image here; run stock Clair3 locally):
  conda run -n clair3-torch python /autofs/nas5/xyu2/projects/clair3-api/Clair3/
    run_clair3.py --bam_fn BAM --ref_fn REF --model_path MODELDIR --platform {ont|hifi
    |ilmn} --threads 8 --output OUTDIR --bed_fn REGION.bed --device cpu --longphase /
    autofs/nas5/xyu2/projects/clair3-api/Clair3/longphase --sample_name SAMPLE
Model dirs (each holds pileup.pt + full_alignment.pt):
  ont=/autofs/nas5/xyu2/softwares/clair3_models_pytorch/r1041_e82_400bps_hac_v500
  hifi=/autofs/nas5/xyu2/softwares/clair3_models_pytorch/hifi_revio
  ilmn=/autofs/nas5/xyu2/softwares/clair3_models_pytorch/ilmn
Registered clinical samples (human WGS; determine each one's platform yourself from
    its BAM @PG/@RG header, then pick the matching model/ref):
  HG00136 bam=/autofs/nas5/xyu2/projects/clair3-api/clair3r/paper_experiments/.cache/
    apoe/HG00136_apoe.bam ref=/autofs/bal19/zxzheng/testData/ilmn/data/
    GRCh38_full_analysis_set_plus_decoy_hla.fa
```

```
  HG00150 bam=/autofs/nas5/xyu2/projects/clair3-api/clair3r/paper_experiments/.cache/
    apoe/HG00150_apoe.bam ref=/autofs/bal19/zxzheng/testData/ilmn/data/
    GRCh38_full_analysis_set_plus_decoy_hla.fa
Output is OUTDIR/merge_output.vcf.gz (bgzipped). Build any region BED yourself.
Read genotypes with bcftools (/autofs/nas5/xyu2/softwares/miniforge3/envs/clair3-torch
    /bin/bcftools).
Finish with a single final line 'ANSWER: <value>'.
```

**(b) Skills.** System prompt (`run_shell` + our `clair-variant-caller` skill; full skill at `https://github.com/HKU-BAL/Clair-skills`):

```
You are a genomics analysis assistant equipped with the SKILL below
(read and follow it) and a single tool: run_shell.

===== BEGIN SKILL.md =====
  clair-variant-caller -- the full skill document is provided at
  https://github.com/HKU-BAL/Clair-skills
===== END SKILL.md =====
```

**(c) Tools.** System prompt (only the packaged Clair3-Connect tools; their operations are listed in Table S2):

```
You are a genomics analysis assistant. You can call germline variants and inspect
results ONLY through the provided Clair3-Connect tools. Datasets are referenced by
(sample, platform); you never handle file paths. Think step by step, call tools
as needed, and finish with a single final line of the exact form 'ANSWER: <value>'
(a bare number when a count/fraction is asked).
```

**Table S2.** The Clair3-Connect tools the agent used in the Tools condition, and the operation each performs.

| Tool | Operation |
|---|---|
| `bam_header` | Returns the BAM header (`@HD`, `@PG`, `@RG`, `@SQ` lines) for a registered sample, so the agent determines the sequencing platform itself from the header rather than being given it. |
| `discover_models` | Lists the served Clair3-Connect models with their sequencing platform and calling stage, so the agent can select the model matching a sample's platform without being told model names. |
| `genotypes_at` | Given a `region` and a list of `positions`, calls variants and returns the genotype at each requested site; it is reference-aware: a site with no called variant is reported as `0/0` (homozygous reference), not omitted. |

#### S1.2 The task prompt

All conditions receive the same task prompt, on samples `HG00136` and `HG00150`. The `HG00136` prompt is reproduced verbatim below; the `HG00150` prompt differs only in the sample name.

**Task prompt (identical across all conditions)**

Clinical genomics. Sample HG00136 is a human whole-genome sequencing dataset registered with the service. Determine its APOE diplotype — the $\varepsilon 2/\varepsilon 3/\varepsilon 4$ combination that stratifies Alzheimer'sdisease risk — defined by two SNPs on chr19 (GRCh38): rs429358 at chr19:44908684 (ref T, alt C) and rs7412 at chr19:44908822 (ref C, alt T). The three common haplotypes are $\varepsilon 4$ =(rs429358 C, rs7412 C), $\varepsilon 3$ =(T, C), $\varepsilon 2$ =(T, T). Steps: (1) determine which sequencing platform this sample is and select the matching Clair3 model; (2) call variants and read the genotype at BOTH SNP sites — a site with NO called variant is homozygous reference; (3) combine the two genotypes into the standard clinical diplotype. Finish with one line `ANSWER: <diplotype>`, for example `ANSWER: e4/e4`.

##### S1.3 Results

Agent model: `deepseek-v4-pro`. Each condition was run in ten independent repeats per sample; all 60 runs returned the correct diplotype with no forced answers and no tool errors. The per-repeat metrics (total tokens, wall-clock time, reasoning turns, tool calls) for every run are listed in Table S3, with the three conditions stacked and the two samples side by side. One representative round's full transcript is in §S4.

#### S2 Performance of the full-alignment-only Clair3-Connect client

##### S2.1 Variant-calling accuracy

The following tables report F1, recall, and precision for two configurations across three sequencing platforms (Illumina, Oxford Nanopore, PacBio HiFi) and coverages of 10 to 50×, on chromosomes 20 and 1, for SNPs and indels. Standard Clair3 runs the complete pileup-and-full-alignment pipeline. Full-alignment only is the configuration deployed in the Clair3-Connect client, with the pileup and phasing stages removed. The benchmark uses sample HG003 against the GIAB v4.2.1 truth set within its high-confidence regions, evaluated with `hap.py`.

**Table S3.** Per-repeat agent metrics across the three deployment conditions, with the two whole-genome samples side by side. Tokens are prompt + completion and Wall is wall-clock seconds; Turns and Calls are reasoning turns and tool calls. Every repeat returned the correct diplotype with no forced answer. Mean and SD are over the ten repeats.

| Condition | Rep | HG00136 | | | | HG00150 | | | |
|---|---|---|---|---|---|---|---|---|---|
| | | Tokens | Wall | Turns | Calls | Tokens | Wall | Turns | Calls |
| RawBinary | 1 | 91,150 | 92.09 | 11 | 10 | 88,770 | 93.25 | 11 | 10 |
| | 2 | 43,349 | 75.07 | 7 | 6 | 136,504 | 137.35 | 14 | 13 |
| | 3 | 56,824 | 83.23 | 8 | 7 | 83,032 | 122.71 | 10 | 9 |
| | 4 | 29,119 | 66.58 | 6 | 5 | 74,738 | 103.17 | 9 | 8 |
| | 5 | 127,041 | 118.03 | 14 | 13 | 78,178 | 106.41 | 10 | 9 |
| | 6 | 119,169 | 101.60 | 13 | 12 | 76,049 | 86.90 | 10 | 9 |
| | 7 | 87,461 | 91.67 | 11 | 10 | 51,849 | 75.40 | 8 | 7 |
| | 8 | 113,695 | 83.40 | 12 | 11 | 94,367 | 108.25 | 10 | 9 |
| | 9 | 73,080 | 71.36 | 10 | 9 | 71,666 | 78.77 | 9 | 8 |
| | 10 | 42,350 | 57.60 | 7 | 6 | 77,473 | 74.78 | 10 | 9 |
| | **Mean** | 78,324 | 84.06 | 9.9 | 8.9 | 83,263 | 98.70 | 10.1 | 9.1 |
| | **SD** | 34,911 | 17.74 | 2.8 | 2.8 | 21,842 | 20.88 | 1.6 | 1.6 |
| Skills | 1 | 120,602 | 76.13 | 6 | 5 | 146,002 | 89.67 | 7 | 6 |
| | 2 | 128,282 | 76.99 | 6 | 8 | 96,769 | 82.80 | 5 | 4 |
| | 3 | 150,141 | 89.22 | 7 | 9 | 291,819 | 159.89 | 12 | 11 |
| | 4 | 188,284 | 87.13 | 9 | 8 | 132,134 | 94.45 | 6 | 8 |
| | 5 | 201,354 | 101.61 | 9 | 8 | 254,009 | 115.12 | 11 | 12 |
| | 6 | 167,183 | 100.05 | 8 | 10 | 164,460 | 102.26 | 8 | 7 |
| | 7 | 106,464 | 78.13 | 5 | 8 | 122,703 | 98.49 | 6 | 5 |
| | 8 | 137,356 | 68.49 | 7 | 8 | 119,841 | 67.73 | 6 | 7 |
| | 9 | 145,485 | 67.52 | 7 | 6 | 233,484 | 77.80 | 10 | 11 |
| | 10 | 236,769 | 74.62 | 11 | 10 | 107,926 | 67.44 | 5 | 8 |
| | **Mean** | 158,192 | 81.99 | 7.5 | 8.0 | 166,915 | 95.56 | 7.6 | 7.9 |
| | **SD** | 40,370 | 12.06 | 1.8 | 1.6 | 68,184 | 27.19 | 2.5 | 2.7 |
| Tools | 1 | 11,667 | 20.32 | 3 | 3 | 11,933 | 21.33 | 3 | 3 |
| | 2 | 11,527 | 16.71 | 3 | 3 | 12,515 | 29.85 | 3 | 3 |
| | 3 | 12,122 | 21.11 | 3 | 4 | 11,951 | 23.50 | 3 | 3 |
| | 4 | 11,447 | 19.13 | 3 | 3 | 12,246 | 28.16 | 3 | 3 |
| | 5 | 11,137 | 21.24 | 3 | 2 | 11,825 | 23.05 | 3 | 3 |
| | 6 | 11,352 | 18.78 | 3 | 3 | 12,230 | 29.03 | 3 | 3 |
| | 7 | 11,686 | 16.75 | 3 | 3 | 12,217 | 30.16 | 3 | 3 |
| | 8 | 11,546 | 19.41 | 3 | 3 | 12,218 | 22.61 | 3 | 3 |
| | 9 | 11,456 | 19.28 | 3 | 3 | 13,532 | 51.07 | 3 | 3 |
| | 10 | 12,179 | 20.14 | 3 | 3 | 11,749 | 21.34 | 3 | 3 |
| | **Mean** | 11,612 | 19.29 | 3.0 | 3.0 | 12,242 | 28.01 | 3.0 | 3.0 |
| | **SD** | 324 | 1.57 | 0.0 | 0.5 | 510 | 8.84 | 0.0 | 0.0 |

**Table S4.** Variant-calling accuracy on chromosome 20 for sample HG003. TRUTH.TP, TRUTH.FN, and QUERY.FP are `hap.py` counts against the GIAB v4.2.1 truth set within the high-confidence region, which contains TRUTH.TOTAL = 70,166 SNP and 10,628 indel truth variants. Recall, Precision, and F1 are percentages to two decimals. Clair3 is standard Clair3 (the full pileup-and-full-alignment pipeline); FA-only is the full-alignment-only configuration deployed in the Clair3-Connect client.

| Type | Platform | Config. | Depth | TRUTH.TP | TRUTH.FN | QUERY.FP | Recall | Precision | F1 |
|---|---|---|---|---|---|---|---|---|---|
| SNP | Illumina | Clair3 | 10× | 67,214 | 2,952 | 3,848 | 95.79 | 94.59 | 95.19 |
| | | | 20× | 69,669 | 497 | 511 | 99.29 | 99.27 | 99.28 |
| | | | 30× | 69,883 | 283 | 279 | 99.60 | 99.60 | 99.60 |
| | | | 40× | 69,919 | 247 | 255 | 99.65 | 99.64 | 99.64 |
| | | | 50× | 69,929 | 237 | 238 | 99.66 | 99.66 | 99.66 |
| | | FA-only | 10× | 61,733 | 8,433 | 12,138 | 87.98 | 83.58 | 85.72 |
| | | | 20× | 68,108 | 2,058 | 2,213 | 97.07 | 96.86 | 96.96 |
| | | | 30× | 69,326 | 840 | 809 | 98.80 | 98.85 | 98.83 |
| | | | 40× | 69,613 | 553 | 568 | 99.21 | 99.19 | 99.20 |
| | | | 50× | 69,689 | 477 | 434 | 99.32 | 99.38 | 99.35 |
| | ONT | Clair3 | 10× | 66,697 | 3,469 | 7,358 | 95.06 | 90.07 | 92.49 |
| | | | 20× | 69,746 | 420 | 682 | 99.40 | 99.03 | 99.22 |
| | | | 30× | 69,974 | 192 | 247 | 99.73 | 99.65 | 99.69 |
| | | | 40× | 70,010 | 156 | 169 | 99.78 | 99.76 | 99.77 |
| | | | 50× | 70,061 | 105 | 133 | 99.85 | 99.81 | 99.83 |
| | | FA-only | 10× | 66,244 | 3,922 | 29,136 | 94.41 | 69.46 | 80.03 |
| | | | 20× | 69,446 | 720 | 4,565 | 98.97 | 93.83 | 96.34 |
| | | | 30× | 69,891 | 275 | 882 | 99.61 | 98.75 | 99.18 |
| | | | 40× | 70,023 | 143 | 386 | 99.80 | 99.45 | 99.62 |
| | | | 50× | 70,069 | 97 | 290 | 99.86 | 99.59 | 99.72 |
| | HiFi | Clair3 | 10× | 68,038 | 2,128 | 712 | 96.97 | 98.97 | 97.96 |
| | | | 20× | 69,923 | 243 | 87 | 99.65 | 99.88 | 99.76 |
| | | | 30× | 70,068 | 98 | 49 | 99.86 | 99.93 | 99.90 |
| | | | 40× | 70,099 | 67 | 37 | 99.90 | 99.95 | 99.93 |
| | | | 50× | 70,141 | 25 | 31 | 99.96 | 99.96 | 99.96 |
| | | FA-only | 10× | 68,122 | 2,044 | 831 | 97.09 | 98.80 | 97.93 |
| | | | 20× | 70,015 | 151 | 118 | 99.78 | 99.83 | 99.81 |
| | | | 30× | 70,119 | 47 | 67 | 99.93 | 99.90 | 99.92 |
| | | | 40× | 70,141 | 25 | 52 | 99.96 | 99.93 | 99.95 |
| | | | 50× | 70,149 | 17 | 44 | 99.98 | 99.94 | 99.96 |
| INDEL | Illumina | Clair3 | 10× | 9,761 | 867 | 644 | 91.84 | 94.04 | 92.93 |
| | | | 20× | 10,432 | 196 | 126 | 98.16 | 98.85 | 98.50 |
| | | | 30× | 10,544 | 84 | 49 | 99.21 | 99.56 | 99.38 |
| | | | 40× | 10,567 | 61 | 30 | 99.43 | 99.73 | 99.58 |
| | | | 50× | 10,576 | 52 | 20 | 99.51 | 99.82 | 99.66 |
| | | FA-only | 10× | 7,839 | 2,789 | 1,224 | 73.76 | 86.85 | 79.77 |
| | | | 20× | 9,501 | 1,127 | 634 | 89.40 | 93.95 | 91.62 |
| | | | 30× | 9,987 | 641 | 377 | 93.97 | 96.49 | 95.21 |
| | | | 40× | 10,155 | 473 | 267 | 95.55 | 97.53 | 96.53 |
| | | | 50× | 10,249 | 379 | 222 | 96.43 | 97.96 | 97.19 |
| | ONT | Clair3 | 10× | 7,327 | 3,301 | 7,668 | 68.94 | 49.33 | 57.51 |
| | | | 20× | 8,432 | 2,196 | 4,220 | 79.34 | 67.20 | 72.76 |
| | | | 30× | 8,744 | 1,884 | 2,632 | 82.27 | 77.34 | 79.73 |
| | | | 40× | 8,868 | 1,760 | 1,893 | 83.44 | 82.80 | 83.12 |
| | | | 50× | 8,972 | 1,656 | 1,545 | 84.42 | 85.66 | 85.03 |
| | | FA-only | 10× | 6,792 | 3,836 | 10,331 | 63.91 | 40.11 | 49.28 |
| | | | 20× | 7,827 | 2,801 | 4,807 | 73.65 | 62.53 | 67.64 |
| | | | 30× | 8,131 | 2,497 | 2,986 | 76.51 | 73.64 | 75.05 |
| | | | 40× | 8,290 | 2,338 | 2,185 | 78.00 | 79.59 | 78.79 |
| | | | 50× | 8,381 | 2,247 | 1,810 | 78.86 | 82.64 | 80.70 |
| | HiFi | Clair3 | 10× | 9,066 | 1,562 | 649 | 85.30 | 93.52 | 89.22 |
| | | | 20× | 10,245 | 383 | 271 | 96.40 | 97.52 | 96.95 |
| | | | 30× | 10,449 | 179 | 167 | 98.32 | 98.49 | 98.40 |
| | | | 40× | 10,518 | 110 | 114 | 98.97 | 98.97 | 98.97 |
| | | | 50× | 10,540 | 88 | 101 | 99.17 | 99.09 | 99.13 |
| | | FA-only | 10× | 8,402 | 2,226 | 874 | 79.06 | 90.84 | 84.54 |
| | | | 20× | 9,934 | 694 | 436 | 93.47 | 95.94 | 94.69 |
| | | | 30× | 10,286 | 342 | 284 | 96.78 | 97.41 | 97.10 |
| | | | 40× | 10,441 | 187 | 166 | 98.24 | 98.50 | 98.37 |
| | | | 50× | 10,513 | 115 | 118 | 98.92 | 98.93 | 98.93 |

**Table S5.** Variant-calling accuracy on chromosome 1 for sample HG003. TRUTH.TP, TRUTH.FN, and QUERY.FP are `hap.py` counts against the GIAB v4.2.1 truth set within the high-confidence region, which contains TRUTH.TOTAL = 256,313 SNP and 40,645 indel truth variants. Recall, Precision, and F1 are percentages to two decimals. Clair3 is standard Clair3 (the full pileup-and-full-alignment pipeline); FA-only is the full-alignment-only configuration deployed in the Clair3-Connect client.

| Type | Platform | Config. | Depth | TRUTH.TP | TRUTH.FN | QUERY.FP | Recall | Precision | F1 |
|---|---|---|---|---|---|---|---|---|---|
| SNP | Illumina | Clair3 | 10× | 244,788 | 11,525 | 14,682 | 95.50 | 94.34 | 94.92 |
| | | | 20× | 253,311 | 3,002 | 2,229 | 98.83 | 99.13 | 98.98 |
| | | | 30× | 254,214 | 2,099 | 1,285 | 99.18 | 99.50 | 99.34 |
| | | | 40× | 254,393 | 1,920 | 989 | 99.25 | 99.61 | 99.43 |
| | | | 50× | 254,440 | 1,873 | 907 | 99.27 | 99.64 | 99.46 |
| | | FA-only | 10× | 225,911 | 30,402 | 43,598 | 88.14 | 83.83 | 85.93 |
| | | | 20× | 247,481 | 8,832 | 8,610 | 96.55 | 96.64 | 96.60 |
| | | | 30× | 251,783 | 4,530 | 3,342 | 98.23 | 98.69 | 98.46 |
| | | | 40× | 252,964 | 3,349 | 2,155 | 98.69 | 99.16 | 98.92 |
| | | | 50× | 253,198 | 3,115 | 1,715 | 98.78 | 99.33 | 99.06 |
| | ONT | Clair3 | 10× | 244,611 | 11,702 | 28,909 | 95.43 | 89.43 | 92.34 |
| | | | 20× | 254,565 | 1,748 | 3,433 | 99.32 | 98.67 | 98.99 |
| | | | 30× | 255,445 | 868 | 1,266 | 99.66 | 99.51 | 99.58 |
| | | | 40× | 255,598 | 715 | 897 | 99.72 | 99.65 | 99.69 |
| | | | 50× | 255,679 | 634 | 644 | 99.75 | 99.75 | 99.75 |
| | | FA-only | 10× | 243,108 | 13,205 | 103,891 | 94.85 | 70.07 | 80.59 |
| | | | 20× | 253,389 | 2,924 | 17,225 | 98.86 | 93.64 | 96.18 |
| | | | 30× | 255,076 | 1,237 | 4,072 | 99.52 | 98.43 | 98.97 |
| | | | 40× | 255,562 | 751 | 2,107 | 99.71 | 99.18 | 99.44 |
| | | | 50× | 255,725 | 588 | 1,481 | 99.77 | 99.42 | 99.60 |
| | HiFi | Clair3 | 10× | 250,008 | 6,305 | 2,320 | 97.54 | 99.08 | 98.30 |
| | | | 20× | 254,798 | 1,515 | 472 | 99.41 | 99.82 | 99.61 |
| | | | 30× | 255,397 | 916 | 319 | 99.64 | 99.88 | 99.76 |
| | | | 40× | 255,630 | 683 | 300 | 99.73 | 99.88 | 99.81 |
| | | | 50× | 255,712 | 601 | 263 | 99.77 | 99.90 | 99.83 |
| | | FA-only | 10× | 250,436 | 5,877 | 2,771 | 97.71 | 98.91 | 98.30 |
| | | | 20× | 255,372 | 941 | 552 | 99.63 | 99.78 | 99.71 |
| | | | 30× | 255,655 | 658 | 450 | 99.74 | 99.82 | 99.78 |
| | | | 40× | 255,735 | 578 | 360 | 99.77 | 99.86 | 99.82 |
| | | | 50× | 255,774 | 539 | 334 | 99.79 | 99.87 | 99.83 |
| INDEL | Illumina | Clair3 | 10× | 37,220 | 3,425 | 2,388 | 91.57 | 94.19 | 92.87 |
| | | | 20× | 39,740 | 905 | 514 | 97.77 | 98.78 | 98.27 |
| | | | 30× | 40,192 | 453 | 205 | 98.89 | 99.51 | 99.20 |
| | | | 40× | 40,328 | 317 | 159 | 99.22 | 99.62 | 99.42 |
| | | | 50× | 40,369 | 276 | 121 | 99.32 | 99.71 | 99.52 |
| | | FA-only | 10× | 29,941 | 10,704 | 4,921 | 73.66 | 86.26 | 79.47 |
| | | | 20× | 36,078 | 4,567 | 2,490 | 88.76 | 93.77 | 91.20 |
| | | | 30× | 38,036 | 2,609 | 1,487 | 93.58 | 96.38 | 94.96 |
| | | | 40× | 38,739 | 1,906 | 1,055 | 95.31 | 97.45 | 96.37 |
| | | | 50× | 39,098 | 1,547 | 840 | 96.19 | 97.98 | 97.08 |
| | ONT | Clair3 | 10× | 28,105 | 12,540 | 28,957 | 69.15 | 49.77 | 57.88 |
| | | | 20× | 32,407 | 8,238 | 15,456 | 79.73 | 68.27 | 73.56 |
| | | | 30× | 33,589 | 7,056 | 9,660 | 82.64 | 78.16 | 80.34 |
| | | | 40× | 34,131 | 6,514 | 6,883 | 83.97 | 83.63 | 83.80 |
| | | | 50× | 34,491 | 6,154 | 5,598 | 84.86 | 86.41 | 85.63 |
| | | FA-only | 10× | 26,123 | 14,522 | 39,351 | 64.27 | 40.39 | 49.61 |
| | | | 20× | 30,010 | 10,635 | 18,146 | 73.83 | 62.91 | 67.94 |
| | | | 30× | 31,134 | 9,511 | 11,025 | 76.60 | 74.38 | 75.47 |
| | | | 40× | 31,752 | 8,893 | 8,100 | 78.12 | 80.14 | 79.12 |
| | | | 50× | 32,144 | 8,501 | 6,849 | 79.08 | 82.85 | 80.92 |
| | HiFi | Clair3 | 10× | 35,403 | 5,242 | 2,324 | 87.10 | 94.03 | 90.43 |
| | | | 20× | 39,392 | 1,253 | 870 | 96.92 | 97.92 | 97.42 |
| | | | 30× | 40,110 | 535 | 469 | 98.68 | 98.89 | 98.79 |
| | | | 40× | 40,311 | 334 | 330 | 99.18 | 99.22 | 99.20 |
| | | | 50× | 40,403 | 242 | 249 | 99.40 | 99.41 | 99.41 |
| | | FA-only | 10× | 33,089 | 7,556 | 3,165 | 81.41 | 91.52 | 86.17 |
| | | | 20× | 38,292 | 2,353 | 1,488 | 94.21 | 96.40 | 95.29 |
| | | | 30× | 39,606 | 1,039 | 795 | 97.44 | 98.11 | 97.78 |
| | | | 40× | 40,026 | 619 | 527 | 98.48 | 98.75 | 98.61 |
| | | | 50× | 40,251 | 394 | 363 | 99.03 | 99.14 | 99.09 |

#### S2.2 Runtime and peak memory

These tables list the wall-clock runtime and peak resident memory of the full-alignment-only client over each whole chromosome at 30× coverage, for sample HG003 on the three sequencing platforms. Each platform and chromosome was run in ten independent repeats. The per-repeat values are given below; their means and standard deviations are summarized in main-text Table 1.

**Table S6.** Per-repeat wall-clock runtime (seconds) of the full-alignment-only Clair3-Connect client, for sample HG003 at 30× coverage, over each whole chromosome on the three platforms. Ten independent repeats per platform and chromosome. The means and standard deviations in main-text Table 1 are computed from these values.

| Platform | Chr | Repeat | | | | | | | | | |
|---|---|---|---|---|---|---|---|---|---|---|---|
| | | 1 | 2 | 3 | 4 | 5 | 6 | 7 | 8 | 9 | 10 |
| Illumina | | 81.26 | 69.76 | 73.77 | 75.03 | 74.00 | 74.67 | 76.04 | 72.66 | 73.65 | 75.66 |
| Oxford Nanopore | 1 | 241.81 | 213.44 | 220.73 | 214.37 | 230.45 | 232.30 | 190.01 | 188.10 | 183.49 | 191.42 |
| PacBio HiFi | | 80.03 | 72.28 | 63.47 | 73.70 | 73.95 | 61.64 | 70.67 | 76.01 | 62.76 | 70.09 |
| Illumina | | 27.28 | 24.64 | 23.79 | 24.41 | 25.68 | 24.84 | 26.66 | 24.54 | 24.48 | 24.54 |
| Oxford Nanopore | 20 | 43.65 | 43.35 | 43.75 | 42.99 | 43.18 | 43.92 | 43.11 | 43.14 | 43.09 | 43.76 |
| PacBio HiFi | | 24.47 | 22.12 | 21.72 | 21.79 | 21.53 | 21.79 | 22.10 | 21.79 | 21.72 | 21.63 |

**Table S7.** Per-repeat peak resident memory (GB) of the full-alignment-only Clair3-Connect client, for sample HG003 at 30× coverage, over each whole chromosome on the three platforms. Ten independent repeats per platform and chromosome. The means and standard deviations in main-text Table 1 are computed from these values.

| Platform | Chr | Repeat | | | | | | | | | |
|---|---|---|---|---|---|---|---|---|---|---|---|
| | | 1 | 2 | 3 | 4 | 5 | 6 | 7 | 8 | 9 | 10 |
| Illumina | | 5.28 | 4.85 | 5.56 | 4.78 | 4.67 | 5.04 | 4.73 | 5.20 | 5.26 | 4.77 |
| Oxford Nanopore | 1 | 11.75 | 8.91 | 8.96 | 9.78 | 11.33 | 12.46 | 10.01 | 11.87 | 9.39 | 12.16 |
| PacBio HiFi | | 7.07 | 6.94 | 6.76 | 6.91 | 6.88 | 7.01 | 6.71 | 6.70 | 6.96 | 6.66 |
| Illumina | | 3.79 | 3.80 | 3.75 | 3.91 | 3.84 | 3.89 | 3.83 | 3.61 | 3.95 | 3.89 |
| Oxford Nanopore | 20 | 7.07 | 7.32 | 7.07 | 7.16 | 7.53 | 7.28 | 6.77 | 7.58 | 6.97 | 6.75 |
| PacBio HiFi | | 4.66 | 4.80 | 4.85 | 4.86 | 4.68 | 4.73 | 4.54 | 4.70 | 4.78 | 4.99 |

### S3 Transport security

This note collects the raw transport-security measurements: the per-configuration performance overhead (plaintext, mTLS, AEAD, mTLS+AEAD) is summarized in Table S8, with the per-round values in Tables S9 and S10. All measurements use sample HG003 on ONT chromosome 20, on a 128-core host with one RTX 4090 over loopback.

**Table S8.** Performance overhead of Clair3-Connect's two transport-security layers relative to plaintext. Mean ± 95% CI over $n = 10$ paired rounds; $\Delta$ is the paired change vs plaintext; $p$ is a paired Wilcoxon signed-rank test (ns: $p \geq 0.05$). The isolated latency puts the full payload encryption on the request critical path; the wall is a real full-alignment-only run.

| Configuration | Mean ± 95% CI | $\Delta$ vs plaintext | Wilcoxon $p$ |
|---|---|---|---|
| *Isolated per-request latency, concurrency 1 (ms)* | | | |
| Plaintext | 13.41 ± 0.06 | – | – |
| mTLS | 14.05 ± 0.09 | +4.8 % ± 0.7 | 0.002 ** |
| AEAD | 20.78 ± 0.17 | +54.9 % ± 1.0 | 0.002 ** |
| mTLS+AEAD | 21.55 ± 0.12 | +60.6 % ± 0.9 | 0.002 ** |
| *End-to-end whole-region wall (s)* | | | |
| Plaintext | 29.05 ± 0.49 | – | – |
| mTLS | 29.36 ± 0.32 | +1.1 % ± 2.0 | 0.232 ns |
| AEAD | 30.36 ± 0.52 | +4.5 % ± 2.1 | 0.004 ** |
| mTLS+AEAD | 31.12 ± 0.29 | +7.2 % ± 2.3 | 0.002 ** |

**Table S9.** Per-round isolated per-request latency at concurrency 1 (ms), by configuration. Ten paired rounds; means summarized in Table S8.

| Round | Plaintext | mTLS | AEAD | mTLS+AEAD |
|---|---|---|---|---|
| 1 | 13.36 | 14.07 | 20.82 | 21.63 |
| 2 | 13.28 | 13.89 | 20.55 | 21.30 |
| 3 | 13.41 | 14.32 | 20.76 | 21.81 |
| 4 | 13.52 | 14.06 | 21.06 | 21.63 |
| 5 | 13.44 | 14.05 | 20.51 | 21.46 |
| 6 | 13.54 | 14.11 | 20.84 | 21.50 |
| 7 | 13.50 | 14.05 | 21.27 | 21.74 |
| 8 | 13.38 | 13.98 | 20.77 | 21.66 |
| 9 | 13.39 | 13.89 | 20.57 | 21.37 |
| 10 | 13.35 | 14.11 | 20.67 | 21.36 |
| **Mean** | 13.41 | 14.05 | 20.78 | 21.55 |

**Table S10.** Per-round end-to-end whole-region wall-clock time (s), by configuration. Ten paired rounds; means summarized in Table S8.

| Round | Plaintext | mTLS | AEAD | mTLS+AEAD |
|---|---|---|---|---|
| 1 | 28.04 | 28.95 | 29.41 | 31.37 |
| 2 | 29.75 | 28.50 | 30.86 | 31.46 |
| 3 | 29.61 | 29.27 | 30.64 | 30.10 |
| 4 | 29.47 | 29.90 | 29.37 | 31.25 |
| 5 | 29.61 | 29.51 | 30.64 | 31.33 |
| 6 | 28.36 | 29.79 | 31.02 | 31.55 |
| 7 | 29.34 | 29.93 | 30.94 | 30.91 |
| 8 | 29.59 | 29.20 | 30.58 | 31.02 |
| 9 | 28.26 | 29.20 | 30.89 | 31.01 |
| 10 | 28.44 | 29.36 | 29.23 | 31.17 |
| **Mean** | 29.05 | 29.36 | 30.36 | 31.12 |

#### S4 Full agent transcripts

The complete round-by-round trajectory of every run is reproduced below, verbatim and without truncation. Each turn shows, in colour-coded boxes, the model's chain-of-thought (`reasoning_content`), the tool call (name and JSON arguments), and the returned result, ending with the final answer. System prompts are in §S1.1; the task prompt in §S1.2.

chain-of-thought | tool call | tool result | final answer

#### RawBinary

##### RawBinary ▪ HG00136

13 turns, 12 tool calls, 112,850 tokens, 90.3 s; answer: $\varepsilon 4/\varepsilon 4$

*System prompt: §S1.1 (RawBinary); task prompt: §S1.2 (sample HG00136).*

**Turn 1 ▪ reasoning**

Let me work through this step by step.
1. First, I need to determine the sequencing platform for HG00136 by checking the BAM header.
2. Then select the appropriate Clair3 model.
3. Create a BED file for the region containing both SNPs (chr19:44908684 and chr19:44908822).
4. Run Clair3 on that region.
5. Read the VCF output for both SNP sites.
6. Determine the diplotype.
Let's start by checking the BAM header to determine the platform.

**Turn 1 ▪ tool call: run_shell**

```
run_shell:
samtools view -H /autofs/nas5/xyu2/projects/clair3-api/clair3r/paper_experiments/.
    cache/apoe/HG00136_apoe.bam | head -50
```

**Result ▪ run_shell**

```
returncode: 0
stdout:
et_plus_decoy_hla.fa
@SQ     SN:chr11     LN:135086622     M5:1511375dc2dd1b633af8cf439ae90cec     UR:/gpfs/
    internal/sweng/production/Resources/GRCh38_1000genomes/
    GRCh38_full_analysis_set_plus_decoy_hla.fa
@SQ     SN:chr12     LN:133275309     M5:96e414eace405d8c27a6d35ba19df56f     UR:/gpfs/
    internal/sweng/production/Resources/GRCh38_1000genomes/
    GRCh38_full_analysis_set_plus_decoy_hla.fa
@SQ     SN:chr13     LN:114364328     M5:a5437debe2ef9c9ef8f3ea2874ae1d82     UR:/gpfs/
    internal/sweng/production/Resources/GRCh38_1000genomes/
    GRCh38_full_analysis_set_plus_decoy_hla.fa
@SQ     SN:chr14     LN:107043718     M5:e0f0eecc3bcab6178c62b6211565c807     UR:/gpfs/
    internal/sweng/production/Resources/GRCh38_1000genomes/
    GRCh38_full_analysis_set_plus_decoy_hla.fa
@SQ     SN:chr15     LN:101991189     M5:f036bd11158407596ca6bf3581454706     UR:/gpfs/
    internal/sweng/production/Resources/GRCh38_1000genomes/
    GRCh38_full_analysis_set_plus_decoy_hla.fa
```

```
@SQ    SN:chr16    LN:90338345    M5:db2d37c8b7d019caaf2dd64ba3a6f33a    UR:/gpfs/
    internal/sweng/production/Resources/GRCh38_1000genomes/
    GRCh38_full_analysis_set_plus_decoy_hla.fa
@SQ    SN:chr17    LN:83257441    M5:f9a0fb01553adb183568e3eb9d8626db    UR:/gpfs/
    internal/sweng/production/Resources/GRCh38_1000genomes/
    GRCh38_full_analysis_set_plus_decoy_hla.fa
@SQ    SN:chr18    LN:80373285    M5:11eeaa801f6b0e2e36a1138616b8ee9a    UR:/gpfs/
    internal/sweng/production/Resources/GRCh38_1000genomes/
    GRCh38_full_analysis_set_plus_decoy_hla.fa
@SQ    SN:chr19    LN:58617616    M5:85f9f4fc152c58cb7913c06d6b98573a    UR:/gpfs/
    internal/sweng/production/Resources/GRCh38_1000genomes/
    GRCh38_full_analysis_set_plus_decoy_hla.fa
@SQ    SN:chr20    LN:64444167    M5:b18e6c531b0bd70e949a7fc20859cb01    UR:/gpfs/
    internal/sweng/production/Resources/GRCh38_1000genomes/
    GRCh38_full_analysis_set_plus_decoy_hla.fa
@SQ    SN:chr21    LN:46709983    M5:974dc7aec0b755b19f031418fdedf293    UR:/gpfs/
    internal/sweng/production/Resources/GRCh38_1000genomes/
    GRCh38_full_analysis_set_plus_decoy_hla.fa
@SQ    SN:chr22    LN:50818468    M5:ac37ec46683600f808cdd41eac1d55cd    UR:/gpfs/
    internal/sweng/production/Resources/GRCh38_1000genomes/
    GRCh38_full_analysis_set_plus_decoy_hla.fa
@SQ    SN:chrX    LN:156040895    M5:2b3a55ff7f58eb308420c8a9b11cac50    UR:/gpfs/
    internal/sweng/production/Resources/GRCh38_1000genomes/
    GRCh38_full_analysis_set_plus_decoy_hla.fa
@SQ    SN:chrY    LN:57227415    M5:ce3e31103314a704255f3cd90369ecce    UR:/gpfs/
    internal/sweng/production/Resources/GRCh38_1000genomes/
    GRCh38_full_analysis_set_plus_decoy_hla.fa
@SQ    SN:chrM    LN:16569    M5:c68f52674c9fb33aef52dcf399755519    UR:/gpfs/internal
    /sweng/production/Resources/GRCh38_1000genomes/
    GRCh38_full_analysis_set_plus_decoy_hla.fa
@SQ    SN:chr1_KI270706v1_random    LN:175055    M5:62def1a794b3e18192863d187af956e6
     UR:/gpfs/internal/sweng/production/Resources/GRCh38_1000genomes/
    GRCh38_full_analysis_set_plus_decoy_hla.fa
@SQ    SN:chr1_KI270707v1_random    LN:32032    M5:78135804eb15220565483b7cdd02f3be
     UR:/gpfs/internal/sweng/production/Resources/GRCh38_1000genomes/
    GRCh38_full_analysis_set_plus_decoy_hla.fa
@SQ    SN:chr1_KI270708v1_random    LN:127682    M5:1e95e047b98ed92148dd84d6c037158c
     UR:/gpfs/internal/sweng/production/Resources/GRCh38_1000genomes/
    GRCh38_full_analysis_set_plus_decoy_hla.fa
@SQ    SN:chr1_KI270709v1_random    LN:66860    M5:4e2db2933ea96aee8dab54af60ecb37d
     UR:/gpfs/internal/sweng/production/Resources/GRCh38_1000genomes/
    GRCh38_full_analysis_set_plus_decoy_hla.fa
@SQ    SN:chr1_KI270710v1_random    LN:40176    M5:9949f776680c6214512ee738ac5da289
     UR:/gpfs/internal/sweng/production/Resources/GRCh38_1000genomes/
    GRCh38_full_analysis_set_plus_decoy_hla.fa
@SQ    SN:chr1_KI270711v1_random    LN:42210    M5:af383f98cf4492c1f1c4e750c26cbb40
     UR:/gpfs/internal/sweng/production/Resources/GRCh38_1000genomes/
    GRCh38_full_analysis_set_plus_decoy_hla.fa
@SQ    SN:chr1_KI270712v1_random    LN:176043    M5:c38a0fecae6a1838a405406f724d6838
     UR:/gpfs/internal/sweng/production/Resources/GRCh38_1000genomes/
    GRCh38_full_analysis_set_plus_decoy_hla.fa
@SQ    SN:chr1_KI270713v1_random    LN:40745    M5:cb78d48cc0adbc58822a1c6fe89e3569
     UR:/gpfs/internal/sweng/production/Resources/GRCh38_1000genomes/
    GRCh38_full_analysis_set_plus_decoy_hla.fa
```

**Turn 2 ▪ reasoning**

The output got cut off. Let me look for the @RG and @PG lines that contain platform information.

**Turn 2 ▪ tool call: run_shell**

```
run_shell:
samtools view -H /autofs/nas5/xyu2/projects/clair3-api/clair3r/paper_experiments/.
    cache/apoe/HG00136_apoe.bam | grep -E '@RG|@PG'
```

**Result ▪ run_shell**

```
returncode: 0
stdout:
PL:illumina\tPM:Unknown\tLB:HG00136\tDS:GRCh38\tSM:HG00136\tCN:NYGenome\tPU:HFJFFDSXX
    .3.CCAAGTCT /gpfs/internal/sweng/production/Resources/GRCh38_1000genomes/
    GRCh38_full_analysis_set_plus_decoy_hla.fa /gpfs/internal/sequence/demux/
    ILLUMINA_NOVASEQ/SNA00297/181121_SNA00297_0049_BHFJFFDSXX/01.2018-11-23_11.09.41/
    CCDG_13607_B01_GRM_WGS/NYGC/Project_CCDG_13607_B01_GRM_WGS/Sample_HG00136/
    HG00136_CCAAGTCT-AAGGATGA_HFJFFDSXX_L003/fastq/HG00136_CCAAGTCT-
    AAGGATGA_HFJFFDSXX_L003_001.R1.fastq.gz /gpfs/internal/sequence/demux/
    ILLUMINA_NOVASEQ/SNA00297/181121_SNA00297_0049_BHFJFFDSXX/01.2018-11-23_11.09.41/
    CCDG_13607_B01_GRM_WGS/NYGC/Project_CCDG_13607_B01_GRM_WGS/Sample_HG00136/
    HG00136_CCAAGTCT-AAGGATGA_HFJFFDSXX_L003/fastq/HG00136_CCAAGTCT-
    AAGGATGA_HFJFFDSXX_L003_001.R2.fastq.gz
@PG    ID:bwa.B    PN:bwa    VN:0.7.15-r1140    CL:/data/NYGC/Software/bwa/bwa-0.7.15/
    bwa mem -Y -K 100000000 -t 16 -R @RG\tID:HG00136_CCAAGTCT-AAGGATGA_HFM5KDSXX_L002\
    tPL:illumina\tPM:Unknown\tLB:HG00136\tDS:GRCh38\tSM:HG00136\tCN:NYGenome\tPU:
    HFM5KDSXX.2.CCAAGTCT /gpfs/internal/sweng/production/Resources/GRCh38_1000genomes/
    GRCh38_full_analysis_set_plus_decoy_hla.fa /gpfs/internal/sequence/demux/
    ILLUMINA_NOVASEQ/SNA00297/181121_SNA00297_0048_AHFM5KDSXX/01.2018-11-26_15.18.32/
    CCDG_13607_B01_GRM_WGS/NYGC/Project_CCDG_13607_B01_GRM_WGS/Sample_HG00136/
    HG00136_CCAAGTCT-AAGGATGA_HFM5KDSXX_L002/fastq/HG00136_CCAAGTCT-
    AAGGATGA_HFM5KDSXX_L002_001.R1.fastq.gz /gpfs/internal/sequence/demux/
    ILLUMINA_NOVASEQ/SNA00297/181121_SNA00297_0048_AHFM5KDSXX/01.2018-11-26_15.18.32/
    CCDG_13607_B01_GRM_WGS/NYGC/Project_CCDG_13607_B01_GRM_WGS/Sample_HG00136/
    HG00136_CCAAGTCT-AAGGATGA_HFM5KDSXX_L002/fastq/HG00136_CCAAGTCT-
    AAGGATGA_HFM5KDSXX_L002_001.R2.fastq.gz
@PG    ID:GATK PrintReads    VN:3.5-0-g36282e4    CL:readGroup=null platform=null
    number=-1 sample_file=[] sample_name=[] simplify=false no_pg_tag=false
@PG    ID:samtools    PN:samtools    PP:GATK PrintReads    VN:1.23    CL:samtools view
     -T /autofs/bal19/zxzheng/testData/ilmn/data/
    GRCh38_full_analysis_set_plus_decoy_hla.fa -b -o paper_experiments/.cache/apoe/
    HG00136_apoe.bam https://ftp.sra.ebi.ac.uk/vol1/run/ERR324/ERR3240142/HG00136.
    final.cram chr19:44905000-44912000
@PG    ID:samtools.1    PN:samtools    PP:samtools    VN:1.22.1    CL:samtools view -H
     /autofs/nas5/xyu2/projects/clair3-api/clair3r/paper_experiments/.cache/apoe/
    HG00136_apoe.bam
@RG    ID:HG00136_CCAAGTCT-AAGGATGA_HFJFFDSXX_L001    PL:illumina    PM:Unknown    LB:
    HG00136    DS:GRCh38    SM:HG00136    CN:NYGenome    PU:HFJFFDSXX.1.CCAAGTCT
@RG    ID:HG00136_CCAAGTCT-AAGGATGA_HFJFFDSXX_L002    PL:illumina    PM:Unknown    LB:
    HG00136    DS:GRCh38    SM:HG00136    CN:NYGenome    PU:HFJFFDSXX.2.CCAAGTCT
@RG    ID:HG00136_CCAAGTCT-AAGGATGA_HFJFFDSXX_L003    PL:illumina    PM:Unknown    LB:
    HG00136    DS:GRCh38    SM:HG00136    CN:NYGenome    PU:HFJFFDSXX.3.CCAAGTCT
@RG    ID:HG00136_CCAAGTCT-AAGGATGA_HFJFFDSXX_L004    PL:illumina    PM:Unknown    LB:
    HG00136    DS:GRCh38    SM:HG00136    CN:NYGenome    PU:HFJFFDSXX.4.CCAAGTCT
@RG    ID:HG00136_CCAAGTCT-AAGGATGA_HFKLNDSXX_L001    PL:illumina    PM:Unknown    LB:
    HG00136    DS:GRCh38    SM:HG00136    CN:NYGenome    PU:HFKLNDSXX.1.CCAAGTCT
```

```
@RG    ID:HG00136_CCAAGTCT-AAGGATGA_HFKLNDSXX_L002    PL:illumina    PM:Unknown    LB:
    HG00136    DS:GRCh38    SM:HG00136    CN:NYGenome    PU:HFKLNDSXX.2.CCAAGTCT
@RG    ID:HG00136_CCAAGTCT-AAGGATGA_HFKLNDSXX_L003    PL:illumina    PM:Unknown    LB:
    HG00136    DS:GRCh38    SM:HG00136    CN:NYGenome    PU:HFKLNDSXX.3.CCAAGTCT
@RG    ID:HG00136_CCAAGTCT-AAGGATGA_HFKLNDSXX_L004    PL:illumina    PM:Unknown    LB:
    HG00136    DS:GRCh38    SM:HG00136    CN:NYGenome    PU:HFKLNDSXX.4.CCAAGTCT
@RG    ID:HG00136_CCAAGTCT-AAGGATGA_HFM5KDSXX_L001    PL:illumina    PM:Unknown    LB:
    HG00136    DS:GRCh38    SM:HG00136    CN:NYGenome    PU:HFM5KDSXX.1.CCAAGTCT
@RG    ID:HG00136_CCAAGTCT-AAGGATGA_HFM5KDSXX_L002    PL:illumina    PM:Unknown    LB:
    HG00136    DS:GRCh38    SM:HG00136    CN:NYGenome    PU:HFM5KDSXX.2.CCAAGTCT
@RG    ID:HG00136_CCAAGTCT-AAGGATGA_HFM5KDSXX_L003    PL:illumina    PM:Unknown    LB:
    HG00136    DS:GRCh38    SM:HG00136    CN:NYGenome    PU:HFM5KDSXX.3.CCAAGTCT
@RG    ID:HG00136_CCAAGTCT-AAGGATGA_HFM5KDSXX_L004    PL:illumina    PM:Unknown    LB:
    HG00136    DS:GRCh38    SM:HG00136    CN:NYGenome    PU:HFM5KDSXX.4.CCAAGTCT
```

**Turn 3 ▪ reasoning**

PL:illumina — so this is an Illumina platform sample. The model is ‘ilmn‘ at ‘/autofs/nas5/xyu2/softwares/clair3_models_pytorch/ilmn‘.
Now I need to:
1. Create a BED file for chr19 covering both SNPs (rs429358 at chr19:44908684, rs7412 at chr19:44908822). I’ll make a region that includes both, say chr19:44908680-44908830.
2. Run Clair3 with the ilmn model.
3. Read the output VCF.
Let me create a working directory and the BED file.

**Turn 3 ▪ tool call: run_shell**

```
run_shell:
mkdir -p /tmp/hg00136_apoe && echo -e "chr19\t44908680\t44908830" > /tmp/hg00136_apoe/
    region.bed && cat /tmp/hg00136_apoe/region.bed
```

**Result ▪ run_shell**

```
returncode: 0
stdout:
chr19    44908680    44908830
```

**Turn 4 ▪ reasoning**

Now let’s run Clair3. The model for ilmn is at ‘/autofs/nas5/xyu2/softwares/clair3_models_pytorch/ilmn‘. The reference is ‘/autofs/bal19/zxzheng/testData/ilmn/data/GRCh38_full_analysis_set_plus_decoy_hla.fa‘.

**Turn 4 ▪ tool call: run_shell**

```
run_shell:
conda run -n clair3-torch python /autofs/nas5/xyu2/projects/clair3-api/Clair3/
    run_clair3.py --bam_fn /autofs/nas5/xyu2/projects/clair3-api/clair3r/
    paper_experiments/.cache/apoe/HG00136_apoe.bam --ref_fn /autofs/bal19/zxzheng/
    testData/ilmn/data/GRCh38_full_analysis_set_plus_decoy_hla.fa --model_path /autofs
    /nas5/xyu2/softwares/clair3_models_pytorch/ilmn --platform ilmn --threads 8 --
    output /tmp/hg00136_apoe --bed_fn /tmp/hg00136_apoe/region.bed --device cpu --
    sample_name HG00136 2>&1
```

**Result ▪ run_shell**

```
returncode: 0
stdout:
tput/pileup_chr19_9.vcf, remove empty file
[INFO] Delay 1 seconds before starting variant calling ...
[mpileup] 1 samples in 1 input files
Calling variants ...
Total processed positions in chr19 (chunk 10/12) : 0
Total time elapsed: 0.00 s
[INFO] No vcf output for file /tmp/hg00136_apoe/tmp/pileup_output/pileup_chr19_10.vcf,
     remove empty file
[INFO] Delay 3 seconds before starting variant calling ...
[mpileup] 1 samples in 1 input files
Calling variants ...
Total processed positions in chr19 (chunk 8/12) : 0
Total time elapsed: 0.00 s
[INFO] No vcf output for file /tmp/hg00136_apoe/tmp/pileup_output/pileup_chr19_8.vcf,
    remove empty file
[INFO] Delay 2 seconds before starting variant calling ...
[mpileup] 1 samples in 1 input files
Calling variants ...
Total processed positions in chr19 (chunk 11/12) : 0
Total time elapsed: 0.00 s
[INFO] No vcf output for file /tmp/hg00136_apoe/tmp/pileup_output/pileup_chr19_11.vcf,
     remove empty file
[INFO] Delay 3 seconds before starting variant calling ...
[mpileup] 1 samples in 1 input files
Calling variants ...
Total processed positions in chr19 (chunk 12/12) : 0
Total time elapsed: 0.00 s
[INFO] No vcf output for file /tmp/hg00136_apoe/tmp/pileup_output/pileup_chr19_12.vcf,
     remove empty file

[INFO] 2/7 Select heterozygous SNP variants for Whatshap phasing and haplotagging
[93m[WARNING] Cannot find any 0/1 variant in pileup output using variant quality cut-
    off proportion: 0.30000000000000004, total heterozygous variants: 0[0m
[93m[WARNING] Set low variant quality score cut-off to 0.0[0m
[INFO] Select heterozygous pileup variants exceeding phasing quality cutoff 0
[INFO] Total heterozygous SNP positions selected: chr19: 0

[INFO] 3/7 Phase VCF file using Whatshap
This is WhatsHap 2.8 running under Python 3.11.14
Working on 1 sample from 1 family

# Resource usage
Maximum memory usage: 0.071 GB
Time spent reading BAM/CRAM:                    0.0 s
Time spent parsing VCF:                         0.0 s
Time spent selecting reads:                     0.0 s
Time spent phasing:                             0.0 s
Time spent writing VCF:                         0.0 s
Time spent finding components:                  0.0 s
Time spent on rest:                             0.1 s
Total elapsed time:                             0.1 s

[INFO] 4/7 Haplotag input BAM file using Whatshap
Found 1 sample(s) in input VCF
Found 1 sample(s) in BAM file
```

```
Number of supplementary alignments: 0
Number of non-singleton groups: 0
Skipped 0 groups
Found 0 reads covering 0 variants

== SUMMARY ==
Total alignments processed:                    1978
Alignments that could be tagged:                  0
Alignments spanning multiple phase sets:          0
Finished in 0.5 s

[INFO] 5/7 Select candidates for full-alignment calling
[93m[WARNING] Cannot find any low-quality 0/1 or 1/1 variant in pileup output using
    variant quality cut-off proportion: 0.3, total variants: 1[0m
[93m[WARNING] Set low variant quality score cut-off to 0.0[0m
[93m[WARNING] Cannot find any low-quality 0/0 reference calls in pileup output using
    reference quality cut-off proportion: 0.3, total reference calls: 0[0m
[93m[WARNING] Set low reference quality score cut-off to 0.0[0m
[INFO] Set variants quality cutoff 0.0
[INFO] Set reference calls quality cutoff 0.0
[93m[WARNING] Cannot find any low-quality 0/0, 0/1 or 1/1 variant in pileup output in
    contig chr19[0m
[93m[WARNING] Copying pileup.vcf.gz to /tmp/hg00136_apoe/merge_output.vcf.gz[0m
[INFO] Low quality reference calls to be processed in chr19: 0
[INFO] Low quality variants to be processed in chr19: 0

[INFO] 6/7 Call low-quality variants using full-alignment model
[INFO] Delay 1 seconds before starting variant calling ...
[93m[WARNING] Skip full-alignment variant calling for empty full-alignment regions[0m
[93m[WARNING] No vcf file found with prefix:/tmp/hg00136_apoe/tmp/
    full_alignment_output/full_alignment, output empty vcf file[0m
[INFO] Exit in full-alignment variant calling
```

**Turn 5 ▪ reasoning**

The run completed. Let me check the output VCF.

**Turn 5 ▪ tool call: run_shell**

```
run_shell:
zcat /tmp/hg00136_apoe/merge_output.vcf.gz 2>/dev/null || cat /tmp/hg00136_apoe/
    merge_output.vcf.gz
```

**Result ▪ run_shell**

```
returncode: 0
stdout:
tig=<ID=HLA-C*15:05:01,length=3349>
##contig=<ID=HLA-C*15:05:02,length=3349>
##contig=<ID=HLA-C*15:13,length=2895>
##contig=<ID=HLA-C*15:16,length=3066>
##contig=<ID=HLA-C*15:17,length=3349>
##contig=<ID=HLA-C*15:96Q,length=3349>
##contig=<ID=HLA-C*16:01:01,length=3349>
##contig=<ID=HLA-C*16:02:01,length=2895>
##contig=<ID=HLA-C*16:04:01,length=3349>
##contig=<ID=HLA-C*17:01:01:01,length=3368>
##contig=<ID=HLA-C*17:01:01:02,length=3368>
```

```
##contig=<ID=HLA-C*17:01:01:03,length=3368>
##contig=<ID=HLA-C*17:03,length=3197>
##contig=<ID=HLA-C*18:01,length=3346>
##contig=<ID=HLA-DQA1*01:01:02,length=6489>
##contig=<ID=HLA-DQA1*01:02:01:01,length=6484>
##contig=<ID=HLA-DQA1*01:02:01:02,length=6485>
##contig=<ID=HLA-DQA1*01:02:01:03,length=6485>
##contig=<ID=HLA-DQA1*01:02:01:04,length=6492>
##contig=<ID=HLA-DQA1*01:03:01:01,length=6485>
##contig=<ID=HLA-DQA1*01:03:01:02,length=6492>
##contig=<ID=HLA-DQA1*01:04:01:01,length=6484>
##contig=<ID=HLA-DQA1*01:04:01:02,length=6485>
##contig=<ID=HLA-DQA1*01:05:01,length=6485>
##contig=<ID=HLA-DQA1*01:07,length=5959>
##contig=<ID=HLA-DQA1*01:10,length=5790>
##contig=<ID=HLA-DQA1*01:11,length=5926>
##contig=<ID=HLA-DQA1*02:01,length=6403>
##contig=<ID=HLA-DQA1*03:01:01,length=6437>
##contig=<ID=HLA-DQA1*03:02,length=6437>
##contig=<ID=HLA-DQA1*03:03:01,length=6437>
##contig=<ID=HLA-DQA1*04:01:02:01,length=5853>
##contig=<ID=HLA-DQA1*04:01:02:02,length=5666>
##contig=<ID=HLA-DQA1*04:02,length=6210>
##contig=<ID=HLA-DQA1*05:01:01:01,length=5806>
##contig=<ID=HLA-DQA1*05:01:01:02,length=6529>
##contig=<ID=HLA-DQA1*05:03,length=6121>
##contig=<ID=HLA-DQA1*05:05:01:01,length=6593>
##contig=<ID=HLA-DQA1*05:05:01:02,length=6597>
##contig=<ID=HLA-DQA1*05:05:01:03,length=6393>
##contig=<ID=HLA-DQA1*05:11,length=6589>
##contig=<ID=HLA-DQA1*06:01:01,length=5878>
##contig=<ID=HLA-DQB1*02:01:01,length=7480>
##contig=<ID=HLA-DQB1*02:02:01,length=7471>
##contig=<ID=HLA-DQB1*03:01:01:01,length=7231>
##contig=<ID=HLA-DQB1*03:01:01:02,length=7230>
##contig=<ID=HLA-DQB1*03:01:01:03,length=7231>
##contig=<ID=HLA-DQB1*03:02:01,length=7126>
##contig=<ID=HLA-DQB1*03:03:02:01,length=7126>
##contig=<ID=HLA-DQB1*03:03:02:02,length=7126>
##contig=<ID=HLA-DQB1*03:03:02:03,length=6800>
##contig=<ID=HLA-DQB1*03:05:01,length=6934>
##contig=<ID=HLA-DQB1*05:01:01:01,length=7090>
##contig=<ID=HLA-DQB1*05:01:01:02,length=7090>
##contig=<ID=HLA-DQB1*05:03:01:01,length=7089>
##contig=<ID=HLA-DQB1*05:03:01:02,length=7089>
##contig=<ID=HLA-DQB1*06:01:01,length=7111>
##contig=<ID=HLA-DQB1*06:02:01,length=7102>
##contig=<ID=HLA-DQB1*06:03:01,length=7103>
##contig=<ID=HLA-DQB1*06:09:01,length=7102>
##contig=<ID=HLA-DRB1*01:01:01,length=10741>
##contig=<ID=HLA-DRB1*01:02:01,length=11229>
##contig=<ID=HLA-DRB1*03:01:01:01,length=13908>
##contig=<ID=HLA-DRB1*03:01:01:02,length=13426>
##contig=<ID=HLA-DRB1*04:03:01,length=15246>
##contig=<ID=HLA-DRB1*07:01:01:01,length=16110>
##contig=<ID=HLA-DRB1*07:01:01:02,length=16120>
##contig=<ID=HLA-DRB1*08:03:02,length=13562>
##contig=<ID=HLA-DRB1*09:21,length=16039>
```

```
##contig=<ID=HLA-DRB1*10:01:01,length=13501>
##contig=<ID=HLA-DRB1*11:01:01,length=13921>
##contig=<ID=HLA-DRB1*11:01:02,length=13931>
##contig=<ID=HLA-DRB1*11:04:01,length=13919>
##contig=<ID=HLA-DRB1*12:01:01,length=13404>
##contig=<ID=HLA-DRB1*12:17,length=11260>
##contig=<ID=HLA-DRB1*13:01:01,length=13935>
##contig=<ID=HLA-DRB1*13:02:01,length=13941>
##contig=<ID=HLA-DRB1*14:05:01,length=13933>
##contig=<ID=HLA-DRB1*14:54:01,length=13936>
##contig=<ID=HLA-DRB1*15:01:01:01,length=11080>
##contig=<ID=HLA-DRB1*15:01:01:02,length=11571>
##contig=<ID=HLA-DRB1*15:01:01:03,length=11056>
##contig=<ID=HLA-DRB1*15:01:01:04,length=11056>
##contig=<ID=HLA-DRB1*15:02:01,length=10313>
##contig=<ID=HLA-DRB1*15:03:01:01,length=11567>
##contig=<ID=HLA-DRB1*15:03:01:02,length=11569>
##contig=<ID=HLA-DRB1*16:02:01,length=11005>
#CHROM    POS     ID    REF    ALT    QUAL    FILTER    INFO    FORMAT    HG00136
chr19    44908684    .    T    C    30.89    PASS    P    GT:GQ:DP:AD:AF
    1/1:30:59:0,59:1.0000
```

**Turn 6 ▪ reasoning**

I see only one variant at chr19:44908684 (rs429358). Let me also check rs7412 at chr19:44908822. The VCF shows only the first position. Let me also look at the whole output more carefully.
The VCF shows:
- chr19:44908684: REF=T, ALT=C, GT=1/1 (homozygous ALT)
But I need rs7412 at chr19:44908822 as well. It seems like there's no call for rs7412. Let me check if it's in the VCF or if it's missing (meaning homozygous reference).
Let me search the VCF more thoroughly.

**Turn 6 ▪ tool call: run_shell**

```
run_shell:
zcat /tmp/hg00136_apoe/merge_output.vcf.gz | grep -v '^#'
```

**Result ▪ run_shell**

```
returncode: 0
stdout:
chr19    44908684    .    T    C    30.89    PASS    P    GT:GQ:DP:AD:AF
    1/1:30:59:0,59:1.0000
```

**Turn 7 ▪ reasoning**

So there's only one variant called: at chr19:44908684 (rs429358), the genotype is 1/1 (homozygous C).
For rs7412 at chr19:44908822, there is no call, which means it's homozygous reference (C/C).
Now let's determine the diplotypes:
rs429358 (chr19:44908684): ref=T, alt=C. Genotype is 1/1 → C/C
rs7412 (chr19:44908822): ref=C, alt=T. No variant called → C/C
Haplotypes:
- $\varepsilon$4: (rs429358=C, rs7412=C)
- $\varepsilon$3: (rs429358=T, rs7412=C)
- $\varepsilon$2: (rs429358=T, rs7412=T)
With rs429358=C/C and rs7412=C/C, both chromosomes carry C at rs429358 and C at rs7412.

So both haplotypes are ε4/ε4.
Wait, but I should double-check that there really is no variant at rs7412. The BAM only covers chr19:44905000-44912000 based on the samtools command used to create it. Both SNPs should be within range (44908684 and 44908822). Let me verify by checking the pileup output or using bcftools to query.
Actually, the VCF only has one variant. In Clair3 output, if a position is not in the VCF, it's homozygous reference. So rs7412 is C/C (homozygous reference).
So:
- rs429358: C/C → ε4 allele on both chromosomes
- rs7412: C/C → consistent with ε4 on both chromosomes
Diplotype: ε4/ε4
Let me also verify using bcftools to be thorough.

**Turn 7 ▪ tool call: run_shell**

```
run_shell:
/autofs/nas5/xyu2/softwares/miniforge3/envs/clair3-torch/bin/bcftools view /tmp/
    hg00136_apoe/merge_output.vcf.gz 2>/dev/null | grep -v '^#'
```

**Result ▪ run_shell**

```
returncode: 0
stdout:
chr19    44908684    .    T    C    30.89    PASS    P    GT:GQ:DP:AD:AF
    1/1:30:59:0,59:1
```

**Turn 8 ▪ reasoning**

Let me also verify by looking at the raw pileup VCF to make sure rs7412 was examined.

**Turn 8 ▪ tool call: run_shell**

```
run_shell:
ls /tmp/hg00136_apoe/tmp/pileup_output/ 2>/dev/null && zcat /tmp/hg00136_apoe/tmp/
    pileup_output/pileup_chr19.vcf.gz 2>/dev/null | grep -v '^#'
```

**Result ▪ run_shell**

```
returncode: 1
stdout:
pileup_chr19_2.vcf
pileup_chr19_3.vcf
```

**Turn 9 ▪ reasoning**

Only two chunk files for pileup. Let me check them both.

**Turn 9 ▪ tool call: run_shell**

```
run_shell:
cat /tmp/hg00136_apoe/tmp/pileup_output/pileup_chr19_2.vcf /tmp/hg00136_apoe/tmp/
    pileup_output/pileup_chr19_3.vcf | grep -v '^#'
```

**Result ▪ run_shell**

```
returncode: 0
stdout:
chr19     44908684     .     T     C     30.89     PASS     P     GT:GQ:DP:AD:AF
    1/1:30:59:0,59:1.0000
chr19     44908684     .     T     C     30.89     PASS     P     GT:GQ:DP:AD:AF
    1/1:30:59:0,59:1.0000
```

**Turn 10 ▪ reasoning**

Both chunks show the same variant - rs429358. No call at rs7412. This confirms that rs7412 is homozygous reference (C/C).
So the data:
- rs429358 (chr19:44908684): C/C → both alleles are $\varepsilon 4$ (since $\varepsilon 4$ requires C at rs429358)
- rs7412 (chr19:44908822): C/C → both alleles are C at rs7412

Combining these:
- Chromosome 1: rs429358=C, rs7412=C → $\varepsilon 4$
- Chromosome 2: rs429358=C, rs7412=C → $\varepsilon 4$

Diplotype: $\varepsilon 4/\varepsilon 4$
I'm confident in this answer. Let me also quickly double-check by looking at the BAM directly at these positions using samtools mpileup to verify.

**Turn 10 ▪ tool call: run_shell**

```
run_shell:
samtools mpileup -f /autofs/bal19/zxzheng/testData/ilmn/data/
    GRCh38_full_analysis_set_plus_decoy_hla.fa -r chr19:44908684-44908822 /autofs/nas5
    /xyu2/projects/clair3-api/clair3r/paper_experiments/.cache/apoe/HG00136_apoe.bam
    2>/dev/null
```

**Result ▪ run_shell**

```
returncode: 0
stdout:
??????????????????????????
chr19     44908793     T     58
    ..,,,.,,,.,..,...,.,.,.,,.,..,,,.,,,,.,,,.,.,...,.,..,.....
    ??????????????????????????????????????????????????????????
chr19     44908794     C     59
    ..,,,.,,,.,..,...,.,.,.,,.,..,,,.,,,,.,,,.,.,...,.,..,...,..
    ???????????????????????????????????????????????????????????
chr19     44908795     C     59     .$
    .,,,.,,,.,..,...,.,.,.,,.,..,,,.,,,,.,,,.,.,...,.,..,...,..
    ???????????????????????????????????????????????????????????
chr19     44908796     T     58     .$,$
    ,,.,,,.,..,...,.,.,.,,.,..,,,.,,,,.,,,.,.,...,.,..,...,..
    ???????????????????????????5??????????????????????????????
chr19     44908797     C     56     ,$
    ,.,,,.,..,...,.,.,.,,.,..,,,.,,,,.,,,.,.,...,.,..,...,..
    >???????????????????????????????????????????????????????
chr19     44908798     C     55
    ,.,,,.,..,...,.,.,.,,.,..,,,.,,,,.,,,.,.,...,.,..,...,..
    ???????????????????????????????????????????????????????
chr19     44908799     G     55
    ,.,,,.,..,...,.,.,.,,.,..,,,.,,,,.,,,.,.,...,.,..,...,..
    ???????????????????????????????????????????????????????
chr19     44908800     C     56     ,$
    .,,,.,..,...,.,.,.,,.,..,,,.,,,,.,,,.,.,...,.,..,...,..^].
```

```
    ??55????????????????????????????????????????????????????
chr19    44908801    G    56    .$
    ,,,.,..,...,.,.,.,,.,..,,,.,,,,.,,,.,.,...,.,..,..,...^].
    ??5???????????????????????????????????????????????????5?
chr19    44908802    A    56
    ,,,.,..,...,.,.,.,,.,..,,,.,,,,.,,,.,.,...,.,..,..,....^].
    ?5??????????????????????????????????????????????????????
chr19    44908803    T    55    ,$,$
    ,.,..,...,.,.,.,,.,..,,,.,,,,.,,,.,.,...,...,......^],
    ???????????????????????????????????????????????????????
chr19    44908804    G    55
    ,.,..,...,.,.,.,,.,..,,,.,,,,.,,,.,.,...,.,..,..,......,
    ???????????????????????????????????????????????????????
chr19    44908805    C    57
    ,.,..,...,.,.,.,,.,..,,,.,,,,.,,,.,.,...,.,..,..,.....,^].^].
    ?????????????????????????????????????????????????????????
chr19    44908806    C    57    ,$
    .,..,...,.,.,.,,.,..,,,.,,,,.,,,.,.,...,.,..,..,.....,..
    ??????????????????????????????????????????????????????5??
chr19    44908807    G    56    .$
    ,..,...,.,.,.,,.,..,,,.,,,,.,,,.,.,...,.,..,..,.....,..
    5?????????????????????????????????????????????????????5?
chr19    44908808    A    54    ,$
    ..,...,.,.,.,,.,..,,,.,,,,.,,,..,...,.,..,..,.....,..
    ??????????????????????????????????????????????????????
chr19    44908809    T    53
    ..,...,.,.,.,,.,..,,,.,,,,.,,,.,.,..,.,..,..,.....,..^].
    ????????????????????????????????????????5????????????
chr19    44908810    G    55    .$
    .,...,.,.,.,,.,..,,,.,,,,.,,,.,.,...,.,..,..,.....,...
    ???????????????????????????????????????????????????????
chr19    44908811    A    51    .,...,.,.,.,,.,..,,,.,,,,.,,,.,.,...,.,....,....,...
     ???????????????????????????????????????????????????
chr19    44908812    C    54    .,...,.,.,.,,.,..,,,.,,,,.,,,.,.,...,.,..,..,.....,...
        ??????????????????????????????????????????????????????
chr19    44908813    C    54    .$
    ,...,.,.,.,,.,..,,,.,,,,.,,,.,.,...,.,..,..,.....,...
    ??????????????????????????????????????????????????????
chr19    44908814    T    53    ,...,.,.,.,,.,..,,,.,,,,.,,,.,.,...,.,..,..,.....,...
       ?????????????????????????????????????????????????????
chr19    44908815    G    53    ,...,.,.,.,,.,..,,,.,,,,.,,,.,.,...,.,..,..,.....,...
       ?????????????????????????????????????????????????????
chr19    44908816    C    53    ,...,.,.,.,,.,..,,,.,,,,.,,,.,.,...,.,..,..,.....,...
       ?????????????????????????????????????????????????????
chr19    44908817    A    53    ,...,.,.,.,,.,..,,,.,,,,.,,,.,.,...,.,..,..,.....,...
       ????????????????????????????????????????5????????????
chr19    44908818    G    53    ,...,.,.,.,,.,..,,,.,,,,.,,,.,.,...,.,..,..,.....,...
       ?????????????????????????????????????????????????????
chr19    44908819    A    51    ,$...,.,.,.,,.,..,,,.,,,,.,,,.,.,...,.,..,..,.....,...
      >????????????????????????5?????????????????????????
chr19    44908820    A    52    .$..,.,.,.,,.,..,,,.,,,,.,,,.,.,...,.,..,..,.....,...
       ????????????5???????????????????????????????????????
chr19    44908821    G    52    ..,.,.,.,,.,..,,,.,,,,.,,,.,.,...,.,..,..,.....,...^],
        ????????????????????????????????????????????????????
chr19    44908822    C    52    ..,.,.,.,,.,..,,,.,,,,.,,,.,.,...,.,..,..,.....,...,
      ????????????????????????????????????????????????????
```

**Turn 11 ▪ reasoning**

Interesting - the mpileup output at chr19:44908684 shows all "?" characters, which seems odd. But I notice the first line doesn't show properly. Let me check the specific positions more carefully.
Actually, I notice the first line of the mpileup output seems truncated/wrong. The mpileup starts from the position after the first one. Let me try to get the positions more specifically.
But actually, the key positions I care about are 44908684 and 44908822. Let me check them specifically.

**Turn 11 ▪ tool call: run_shell**

```
run_shell:
samtools mpileup -f /autofs/bal19/zxzheng/testData/ilmn/data/
    GRCh38_full_analysis_set_plus_decoy_hla.fa -r chr19:44908684-44908684 /autofs/nas5
    /xyu2/projects/clair3-api/clair3r/paper_experiments/.cache/apoe/HG00136_apoe.bam
```

**Result ▪ run_shell**

```
returncode: 0
stdout:
chr19    44908684    T    50    CcCCcCCcCCcCCcCCCCcccCcCCcccCcCCCcCCcccCcccCcCCcCC
    ??????????????????????????????????????????????????
stderr:
[mpileup] 1 samples in 1 input files
```

**Turn 12 ▪ reasoning**

At position 44908684, the reference base is T, and all 50 reads show C. So the genotype is clearly C/C (homozygous alternate). This matches the Clair3 call of 1/1.
Now let me check rs7412 at chr19:44908822:

**Turn 12 ▪ tool call: run_shell**

```
run_shell:
samtools mpileup -f /autofs/bal19/zxzheng/testData/ilmn/data/
    GRCh38_full_analysis_set_plus_decoy_hla.fa -r chr19:44908822-44908822 /autofs/nas5
    /xyu2/projects/clair3-api/clair3r/paper_experiments/.cache/apoe/HG00136_apoe.bam
```

**Result ▪ run_shell**

```
returncode: 0
stdout:
chr19    44908822    C    52    ..,.,.,.,,.,..,,,.,,,,.,,,.,.,...,.,..,..,.....,...,
      ????????????????????????????????????????????????????
stderr:
[mpileup] 1 samples in 1 input files
```

**Turn 13 ▪ reasoning**

At position 44908822 (rs7412), the reference base is C, and all reads show either "." (match to reference) or "," (match on reverse strand). All bases are C. No T variants are present. So it's C/C (homozygous reference).
This confirms:
- rs429358: C/C
- rs7412: C/C
Now combining into haplotypes:
- $\varepsilon 4$ = (rs429358=C, rs7412=C)
- $\varepsilon 3$ = (rs429358=T, rs7412=C)

- ε2 = (rs429358=T, rs7412=T)
Both chromosomes have (C, C) = ε4. So diplotype is ε4/ε4.
The answer is ε4/ε4.

**Final answer**

**Platform:** Illumina (‘PL:illumina‘ in all ‘@RG‘ headers). Used the ‘ilmn‘ model.
**Clair3 calls:**
- **rs429358** (chr19:44908684, ref T→alt C): **1/1** (homozygous ALT = C/C)
- **rs7412** (chr19:44908822, ref C→alt T): no variant called → homozygous REF = C/C
**Haplotype deduction:**
| Haplotype | rs429358 | rs7412 |
|———–|———-|——–|
| ε4 | C | C |
| ε3 | T | C |
| ε2 | T | T |
Both chromosomes carry C at rs429358 and C at rs7412 → both are ε4 haplotypes.
ANSWER: ε4/ε4

#### RawBinary ▪ HG00150

7 turns, 6 tool calls, 40,642 tokens, 62.5 s; answer: e4/e2

*System prompt: §S1.1 (RawBinary); task prompt: §S1.2 (sample HG00150).*

**Turn 1 ▪ reasoning**

Let me work through this step by step.
1. First, I need to determine the sequencing platform for HG00150. The sample is registered with BAM at ‘/autofs/nas5/xyu2/projects/clair3-api/clair3r/paper_experiments/.cache/apoe/HG00150_apoe.bam‘ and ref at ‘/autofs/bal19/zxzheng/testData/ilmn/data/GRCh38_full_analysis_set_plus_decoy_hla.fa‘.
2. Let me check the BAM header to determine the platform.

##### Turn 1 ▪ tool call: run_shell

```
run_shell:
samtools view -H /autofs/nas5/xyu2/projects/clair3-api/clair3r/paper_experiments/.
    cache/apoe/HG00150_apoe.bam 2>&1 | head -80
```

**Result ▪ run_shell**

```
returncode: 0
stdout:
5:cd5e932cfc4c74d05bb64e2126873a3a    UR:/gpfs/internal/sweng/production/Resources/
    GRCh38_1000genomes/GRCh38_full_analysis_set_plus_decoy_hla.fa
@SQ    SN:chr9_KI270720v1_random    LN:39050    M5:8c2683400a4aeeb40abff96652b9b127
     UR:/gpfs/internal/sweng/production/Resources/GRCh38_1000genomes/
    GRCh38_full_analysis_set_plus_decoy_hla.fa
@SQ    SN:chr11_KI270721v1_random    LN:100316    M5:9654b5d3f36845bb9d19a6dbd15d2f22
       UR:/gpfs/internal/sweng/production/Resources/GRCh38_1000genomes/
    GRCh38_full_analysis_set_plus_decoy_hla.fa
@SQ    SN:chr14_GL000009v2_random    LN:201709    M5:862f555045546733591ff7ab15bcecbe
       UR:/gpfs/internal/sweng/production/Resources/GRCh38_1000genomes/
    GRCh38_full_analysis_set_plus_decoy_hla.fa
@SQ    SN:chr14_GL000225v1_random    LN:211173    M5:63945c3e6962f28ffd469719a747e73c
       UR:/gpfs/internal/sweng/production/Resources/GRCh38_1000genomes/
    GRCh38_full_analysis_set_plus_decoy_hla.fa
```

```
@SQ	SN:chr14_KI270722v1_random	LN:194050	M5:51f46c9093929e6edc3b4dfd50d803fc
	UR:/gpfs/internal/sweng/production/Resources/GRCh38_1000genomes/
    GRCh38_full_analysis_set_plus_decoy_hla.fa
@SQ	SN:chr14_GL000194v1_random	LN:191469	M5:6ac8f815bf8e845bb3031b73f812c012
	UR:/gpfs/internal/sweng/production/Resources/GRCh38_1000genomes/
    GRCh38_full_analysis_set_plus_decoy_hla.fa
@SQ	SN:chr14_KI270723v1_random	LN:38115	M5:74a4b480675592095fb0c577c515b5df
	UR:/gpfs/internal/sweng/production/Resources/GRCh38_1000genomes/
    GRCh38_full_analysis_set_plus_decoy_hla.fa
@SQ	SN:chr14_KI270724v1_random	LN:39555	M5:c3fcb15dddf45f91ef7d94e2623ce13b
	UR:/gpfs/internal/sweng/production/Resources/GRCh38_1000genomes/
    GRCh38_full_analysis_set_plus_decoy_hla.fa
@SQ	SN:chr14_KI270725v1_random	LN:172810	M5:edc6402e58396b90b8738a5e37bf773d
	UR:/gpfs/internal/sweng/production/Resources/GRCh38_1000genomes/
    GRCh38_full_analysis_set_plus_decoy_hla.fa
@SQ	SN:chr14_KI270726v1_random	LN:43739	M5:fbe54a3197e2b469ccb2f4b161cfbe86
	UR:/gpfs/internal/sweng/production/Resources/GRCh38_1000genomes/
    GRCh38_full_analysis_set_plus_decoy_hla.fa
@SQ	SN:chr15_KI270727v1_random	LN:448248	M5:84fe18a7bf03f3b7fc76cbac8eb583f1
	UR:/gpfs/internal/sweng/production/Resources/GRCh38_1000genomes/
    GRCh38_full_analysis_set_plus_decoy_hla.fa
@SQ	SN:chr16_KI270728v1_random	LN:1872759	M5:369ff74cf36683b3066a2ca929d9c40d
	UR:/gpfs/internal/sweng/production/Resources/GRCh38_1000genomes/
    GRCh38_full_analysis_set_plus_decoy_hla.fa
@SQ	SN:chr17_GL000205v2_random	LN:185591	M5:458e71cd53dd1df4083dc7983a6c82c4
	UR:/gpfs/internal/sweng/production/Resources/GRCh38_1000genomes/
    GRCh38_full_analysis_set_plus_decoy_hla.fa
@SQ	SN:chr17_KI270729v1_random	LN:280839	M5:2756f6ee4f5780acce31e995443508b6
	UR:/gpfs/internal/sweng/production/Resources/GRCh38_1000genomes/
    GRCh38_full_analysis_set_plus_decoy_hla.fa
@SQ	SN:chr17_KI270730v1_random	LN:112551	M5:48f98ede8e28a06d241ab2e946c15e07
	UR:/gpfs/internal/sweng/production/Resources/GRCh38_1000genomes/
    GRCh38_full_analysis_set_plus_decoy_hla.fa
@SQ	SN:chr22_KI270731v1_random	LN:150754	M5:8176d9a20401e8d9f01b7ca8b51d9c08
	UR:/gpfs/internal/sweng/production/Resources/GRCh38_1000genomes/
    GRCh38_full_analysis_set_plus_decoy_hla.fa
@SQ	SN:chr22_KI270732v1_random	LN:41543	M5:d837bab5e416450df6e1038ae6cd0817
	UR:/gpfs/internal/sweng/production/Resources/GRCh38_1000genomes/
    GRCh38_full_analysis_set_plus_decoy_hla.fa
@SQ	SN:chr22_KI270733v1_random	LN:179772	M5:f1fa05d48bb0c1f87237a28b66f0be0b
	UR:/gpfs/internal/sweng/production/Resources/GRCh38_1000genomes/
    GRCh38_full_analysis_set_plus_decoy_hla.fa
@SQ	SN:chr22_KI270734v1_random	LN:165050	M5:1d17410ae2569c758e6dd51616412d32
	UR:/gpfs/internal/sweng/production/Resources/GRCh38_1000genomes/
    GRCh38_full_analysis_set_plus_decoy_hla.fa
@SQ	SN:chr22_KI270735v1_random	LN:42811	M5:eb6b07b73dd9a47252098ed3d9fb78b8
	UR:/gpfs/internal/sweng/production/Resources/GRCh38_1000genomes/
    GRCh38_full_analysis_set_plus_decoy_hla.fa
@SQ	SN:chr22_KI270736v1_random	LN:181920	M5:2ff189f33cfa52f321accddf648c5616
	UR:/gpfs/internal/sweng/production/Resources/GRCh38_1000genomes/
    GRCh38_full_analysis_set_plus_decoy_hla.fa
```

**Turn 2 ▪ reasoning**

I got the @SQ lines but I need to see @PG and @RG lines to determine the platform.

**Turn 2 ▪ tool call: run_shell**

```
run_shell:
samtools view -H /autofs/nas5/xyu2/projects/clair3-api/clair3r/paper_experiments/.
    cache/apoe/HG00150_apoe.bam 2>&1 | grep -E "^@RG|^@PG"
```

**Result ▪ run_shell**

```
returncode: 0
stdout:
rnal/sweng/production/Resources/GRCh38_1000genomes/
    GRCh38_full_analysis_set_plus_decoy_hla.fa /gpfs/internal/sequence/demux/
    ILLUMINA_NOVASEQ/SNA00404/181107_SNA00404_0041_AHFH25DSXX/01.2018-11-09_10.10.35/
    CCDG_13607_B01_GRM_WGS/NYGC/Project_CCDG_13607_B01_GRM_WGS/Sample_HG00150/
    HG00150_CATAGAGT-TGCCACCA_HFH25DSXX_L002/fastq/HG00150_CATAGAGT-
    TGCCACCA_HFH25DSXX_L002_001.R1.fastq.gz /gpfs/internal/sequence/demux/
    ILLUMINA_NOVASEQ/SNA00404/181107_SNA00404_0041_AHFH25DSXX/01.2018-11-09_10.10.35/
    CCDG_13607_B01_GRM_WGS/NYGC/Project_CCDG_13607_B01_GRM_WGS/Sample_HG00150/
    HG00150_CATAGAGT-TGCCACCA_HFH25DSXX_L002/fastq/HG00150_CATAGAGT-
    TGCCACCA_HFH25DSXX_L002_001.R2.fastq.gz
@PG    ID:bwa.B    PN:bwa    VN:0.7.15-r1140    CL:/data/NYGC/Software/bwa/bwa-0.7.15/
    bwa mem -Y -K 100000000 -t 16 -R @RG\tID:HG00150_CATAGAGT-TGCCACCA_HFKWWDSXX_L002\
    tPL:illumina\tPM:Unknown\tLB:HG00150\tDS:GRCh38\tSM:HG00150\tCN:NYGenome\tPU:
    HFKWWDSXX.2.CATAGAGT /gpfs/internal/sweng/production/Resources/GRCh38_1000genomes/
    GRCh38_full_analysis_set_plus_decoy_hla.fa /gpfs/internal/sequence/demux/
    ILLUMINA_NOVASEQ/SNA00297/181026_SNA00297_0035_BHFKWWDSXX/01.2018-10-28_12.05.06/
    CCDG_13607_B01_GRM_WGS/NYGC/Project_CCDG_13607_B01_GRM_WGS/Sample_HG00150/
    HG00150_CATAGAGT-TGCCACCA_HFKWWDSXX_L002/fastq/HG00150_CATAGAGT-
    TGCCACCA_HFKWWDSXX_L002_001.R1.fastq.gz /gpfs/internal/sequence/demux/
    ILLUMINA_NOVASEQ/SNA00297/181026_SNA00297_0035_BHFKWWDSXX/01.2018-10-28_12.05.06/
    CCDG_13607_B01_GRM_WGS/NYGC/Project_CCDG_13607_B01_GRM_WGS/Sample_HG00150/
    HG00150_CATAGAGT-TGCCACCA_HFKWWDSXX_L002/fastq/HG00150_CATAGAGT-
    TGCCACCA_HFKWWDSXX_L002_001.R2.fastq.gz
@PG    ID:GATK PrintReads    VN:3.5-0-g36282e4    CL:readGroup=null platform=null
    number=-1 sample_file=[] sample_name=[] simplify=false no_pg_tag=false
@PG    ID:samtools    PN:samtools    PP:GATK PrintReads    VN:1.23    CL:/autofs/nas5/
    xyu2/softwares/miniforge3/envs/clair3-torch/bin/samtools view -T /autofs/bal19/
    zxzheng/testData/ilmn/data/GRCh38_full_analysis_set_plus_decoy_hla.fa -b -o /
    autofs/nas5/xyu2/projects/clair3-api/clair3r/paper_experiments/.cache/apoe/
    HG00150_apoe.bam https://ftp.sra.ebi.ac.uk/vol1/run/ERR324/ERR3240154/HG00150.
    final.cram chr19:44903000-44913000
@PG    ID:samtools.1    PN:samtools    PP:samtools    VN:1.22.1    CL:samtools view -H
     /autofs/nas5/xyu2/projects/clair3-api/clair3r/paper_experiments/.cache/apoe/
    HG00150_apoe.bam
@RG    ID:HG00150_CATAGAGT-TGCCACCA_HFGYNDSXX_L001    PL:illumina    PM:Unknown    LB:
    HG00150    DS:GRCh38    SM:HG00150    CN:NYGenome    PU:HFGYNDSXX.1.CATAGAGT
@RG    ID:HG00150_CATAGAGT-TGCCACCA_HFGYNDSXX_L002    PL:illumina    PM:Unknown    LB:
    HG00150    DS:GRCh38    SM:HG00150    CN:NYGenome    PU:HFGYNDSXX.2.CATAGAGT
@RG    ID:HG00150_CATAGAGT-TGCCACCA_HFGYNDSXX_L003    PL:illumina    PM:Unknown    LB:
    HG00150    DS:GRCh38    SM:HG00150    CN:NYGenome    PU:HFGYNDSXX.3.CATAGAGT
@RG    ID:HG00150_CATAGAGT-TGCCACCA_HFGYNDSXX_L004    PL:illumina    PM:Unknown    LB:
    HG00150    DS:GRCh38    SM:HG00150    CN:NYGenome    PU:HFGYNDSXX.4.CATAGAGT
@RG    ID:HG00150_CATAGAGT-TGCCACCA_HFH25DSXX_L001    PL:illumina    PM:Unknown    LB:
    HG00150    DS:GRCh38    SM:HG00150    CN:NYGenome    PU:HFH25DSXX.1.CATAGAGT
@RG    ID:HG00150_CATAGAGT-TGCCACCA_HFH25DSXX_L002    PL:illumina    PM:Unknown    LB:
    HG00150    DS:GRCh38    SM:HG00150    CN:NYGenome    PU:HFH25DSXX.2.CATAGAGT
@RG    ID:HG00150_CATAGAGT-TGCCACCA_HFH25DSXX_L003    PL:illumina    PM:Unknown    LB:
    HG00150    DS:GRCh38    SM:HG00150    CN:NYGenome    PU:HFH25DSXX.3.CATAGAGT
@RG    ID:HG00150_CATAGAGT-TGCCACCA_HFH25DSXX_L004    PL:illumina    PM:Unknown    LB:
    HG00150    DS:GRCh38    SM:HG00150    CN:NYGenome    PU:HFH25DSXX.4.CATAGAGT
```

```
@RG    ID:HG00150_CATAGAGT-TGCCACCA_HFKWWDSXX_L001    PL:illumina    PM:Unknown    LB:
    HG00150    DS:GRCh38    SM:HG00150    CN:NYGenome    PU:HFKWWDSXX.1.CATAGAGT
@RG    ID:HG00150_CATAGAGT-TGCCACCA_HFKWWDSXX_L002    PL:illumina    PM:Unknown    LB:
    HG00150    DS:GRCh38    SM:HG00150    CN:NYGenome    PU:HFKWWDSXX.2.CATAGAGT
@RG    ID:HG00150_CATAGAGT-TGCCACCA_HFKWWDSXX_L003    PL:illumina    PM:Unknown    LB:
    HG00150    DS:GRCh38    SM:HG00150    CN:NYGenome    PU:HFKWWDSXX.3.CATAGAGT
@RG    ID:HG00150_CATAGAGT-TGCCACCA_HFKWWDSXX_L004    PL:illumina    PM:Unknown    LB:
    HG00150    DS:GRCh38    SM:HG00150    CN:NYGenome    PU:HFKWWDSXX.4.CATAGAGT
```

**Turn 3 ▪ reasoning**

The platform is clearly "illumina" (PL:illumina in all @RG lines). So I'll use the ilmn model.
Now I need to:
1. Create a BED file covering the region with the two SNPs: rs429358 at chr19:44908684 and rs7412 at chr19:44908822
2. Run Clair3 with the ilmn model
3. Read the genotypes from the output VCF
Let me create the output directory and BED file, then run Clair3.

**Turn 3 ▪ tool call: run_shell**

```
run_shell:
mkdir -p /tmp/hg00150_apoe && cat > /tmp/hg00150_apoe/region.bed << 'EOF'
chr19    44908600    44908900
EOF
echo "BED file created"
```

**Result ▪ run_shell**

```
returncode: 0
stdout:
BED file created
```

**Turn 4 ▪ reasoning**

Now run Clair3 with the ilmn model.

**Turn 4 ▪ tool call: run_shell**

```
run_shell:
conda run -n clair3-torch python /autofs/nas5/xyu2/projects/clair3-api/Clair3/
    run_clair3.py \
  --bam_fn /autofs/nas5/xyu2/projects/clair3-api/clair3r/paper_experiments/.cache/apoe
    /HG00150_apoe.bam \
  --ref_fn /autofs/bal19/zxzheng/testData/ilmn/data/
    GRCh38_full_analysis_set_plus_decoy_hla.fa \
  --model_path /autofs/nas5/xyu2/softwares/clair3_models_pytorch/ilmn \
  --platform ilmn \
  --threads 8 \
  --output /tmp/hg00150_apoe \
  --bed_fn /tmp/hg00150_apoe/region.bed \
  --device cpu \
  --sample_name HG00150 \
  2>&1 | tail -30
```

**Result ▪ run_shell**

```
returncode: 0
stdout:
Found 1 sample(s) in input VCF
Found 1 sample(s) in BAM file
Number of supplementary alignments: 0
Number of non-singleton groups: 16
Skipped 0 groups
Found 124 reads covering 2 variants

== SUMMARY ==
Total alignments processed:                    3453
Alignments that could be tagged:                248
Alignments spanning multiple phase sets:          0
Finished in 0.5 s

[INFO] 5/7 Select candidates for full-alignment calling
[93m[WARNING] Cannot find any low-quality 0/1 or 1/1 variant in pileup output using
    variant quality cut-off proportion: 0.3, total variants: 2[0m
[93m[WARNING] Set low variant quality score cut-off to 0.0[0m
[93m[WARNING] Cannot find any low-quality 0/0 reference calls in pileup output using
    reference quality cut-off proportion: 0.3, total reference calls: 0[0m
[93m[WARNING] Set low reference quality score cut-off to 0.0[0m
[INFO] Set variants quality cutoff 0.0
[INFO] Set reference calls quality cutoff 0.0
[93m[WARNING] Cannot find any low-quality 0/0, 0/1 or 1/1 variant in pileup output in
    contig chr19[0m
[93m[WARNING] Copying pileup.vcf.gz to /tmp/hg00150_apoe/merge_output.vcf.gz[0m
[INFO] Low quality reference calls to be processed in chr19: 0
[INFO] Low quality variants to be processed in chr19: 0

[INFO] 6/7 Call low-quality variants using full-alignment model
[INFO] Delay 0 seconds before starting variant calling ...
[93m[WARNING] Skip full-alignment variant calling for empty full-alignment regions[0m
[93m[WARNING] No vcf file found with prefix:/tmp/hg00150_apoe/tmp/
    full_alignment_output/full_alignment, output empty vcf file[0m
[INFO] Exit in full-alignment variant calling
```

**Turn 5 ▪ reasoning**

Clair3 completed. Now let me check the output VCF.

**Turn 5 ▪ tool call: run_shell**

```
run_shell:
ls -la /tmp/hg00150_apoe/merge_output.vcf.gz 2>&1 && /autofs/nas5/xyu2/softwares/
    miniforge3/envs/clair3-torch/bin/bcftools view /tmp/hg00150_apoe/merge_output.vcf.
    gz 2>&1
```

**Result ▪ run_shell**

```
returncode: 0
stdout:
15:96Q,length=3349>
##contig=<ID=HLA-C*16:01:01,length=3349>
##contig=<ID=HLA-C*16:02:01,length=2895>
##contig=<ID=HLA-C*16:04:01,length=3349>
##contig=<ID=HLA-C*17:01:01:01,length=3368>
```

```
##contig=<ID=HLA-C*17:01:01:02,length=3368>
##contig=<ID=HLA-C*17:01:01:03,length=3368>
##contig=<ID=HLA-C*17:03,length=3197>
##contig=<ID=HLA-C*18:01,length=3346>
##contig=<ID=HLA-DQA1*01:01:02,length=6489>
##contig=<ID=HLA-DQA1*01:02:01:01,length=6484>
##contig=<ID=HLA-DQA1*01:02:01:02,length=6485>
##contig=<ID=HLA-DQA1*01:02:01:03,length=6485>
##contig=<ID=HLA-DQA1*01:02:01:04,length=6492>
##contig=<ID=HLA-DQA1*01:03:01:01,length=6485>
##contig=<ID=HLA-DQA1*01:03:01:02,length=6492>
##contig=<ID=HLA-DQA1*01:04:01:01,length=6484>
##contig=<ID=HLA-DQA1*01:04:01:02,length=6485>
##contig=<ID=HLA-DQA1*01:05:01,length=6485>
##contig=<ID=HLA-DQA1*01:07,length=5959>
##contig=<ID=HLA-DQA1*01:10,length=5790>
##contig=<ID=HLA-DQA1*01:11,length=5926>
##contig=<ID=HLA-DQA1*02:01,length=6403>
##contig=<ID=HLA-DQA1*03:01:01,length=6437>
##contig=<ID=HLA-DQA1*03:02,length=6437>
##contig=<ID=HLA-DQA1*03:03:01,length=6437>
##contig=<ID=HLA-DQA1*04:01:02:01,length=5853>
##contig=<ID=HLA-DQA1*04:01:02:02,length=5666>
##contig=<ID=HLA-DQA1*04:02,length=6210>
##contig=<ID=HLA-DQA1*05:01:01:01,length=5806>
##contig=<ID=HLA-DQA1*05:01:01:02,length=6529>
##contig=<ID=HLA-DQA1*05:03,length=6121>
##contig=<ID=HLA-DQA1*05:05:01:01,length=6593>
##contig=<ID=HLA-DQA1*05:05:01:02,length=6597>
##contig=<ID=HLA-DQA1*05:05:01:03,length=6393>
##contig=<ID=HLA-DQA1*05:11,length=6589>
##contig=<ID=HLA-DQA1*06:01:01,length=5878>
##contig=<ID=HLA-DQB1*02:01:01,length=7480>
##contig=<ID=HLA-DQB1*02:02:01,length=7471>
##contig=<ID=HLA-DQB1*03:01:01:01,length=7231>
##contig=<ID=HLA-DQB1*03:01:01:02,length=7230>
##contig=<ID=HLA-DQB1*03:01:01:03,length=7231>
##contig=<ID=HLA-DQB1*03:02:01,length=7126>
##contig=<ID=HLA-DQB1*03:03:02:01,length=7126>
##contig=<ID=HLA-DQB1*03:03:02:02,length=7126>
##contig=<ID=HLA-DQB1*03:03:02:03,length=6800>
##contig=<ID=HLA-DQB1*03:05:01,length=6934>
##contig=<ID=HLA-DQB1*05:01:01:01,length=7090>
##contig=<ID=HLA-DQB1*05:01:01:02,length=7090>
##contig=<ID=HLA-DQB1*05:03:01:01,length=7089>
##contig=<ID=HLA-DQB1*05:03:01:02,length=7089>
##contig=<ID=HLA-DQB1*06:01:01,length=7111>
##contig=<ID=HLA-DQB1*06:02:01,length=7102>
##contig=<ID=HLA-DQB1*06:03:01,length=7103>
##contig=<ID=HLA-DQB1*06:09:01,length=7102>
##contig=<ID=HLA-DRB1*01:01:01,length=10741>
##contig=<ID=HLA-DRB1*01:02:01,length=11229>
##contig=<ID=HLA-DRB1*03:01:01:01,length=13908>
##contig=<ID=HLA-DRB1*03:01:01:02,length=13426>
##contig=<ID=HLA-DRB1*04:03:01,length=15246>
##contig=<ID=HLA-DRB1*07:01:01:01,length=16110>
##contig=<ID=HLA-DRB1*07:01:01:02,length=16120>
##contig=<ID=HLA-DRB1*08:03:02,length=13562>
```

```
##contig=<ID=HLA-DRB1*09:21,length=16039>
##contig=<ID=HLA-DRB1*10:01:01,length=13501>
##contig=<ID=HLA-DRB1*11:01:01,length=13921>
##contig=<ID=HLA-DRB1*11:01:02,length=13931>
##contig=<ID=HLA-DRB1*11:04:01,length=13919>
##contig=<ID=HLA-DRB1*12:01:01,length=13404>
##contig=<ID=HLA-DRB1*12:17,length=11260>
##contig=<ID=HLA-DRB1*13:01:01,length=13935>
##contig=<ID=HLA-DRB1*13:02:01,length=13941>
##contig=<ID=HLA-DRB1*14:05:01,length=13933>
##contig=<ID=HLA-DRB1*14:54:01,length=13936>
##contig=<ID=HLA-DRB1*15:01:01:01,length=11080>
##contig=<ID=HLA-DRB1*15:01:01:02,length=11571>
##contig=<ID=HLA-DRB1*15:01:01:03,length=11056>
##contig=<ID=HLA-DRB1*15:01:01:04,length=11056>
##contig=<ID=HLA-DRB1*15:02:01,length=10313>
##contig=<ID=HLA-DRB1*15:03:01:01,length=11567>
##contig=<ID=HLA-DRB1*15:03:01:02,length=11569>
##contig=<ID=HLA-DRB1*16:02:01,length=11005>
##bcftools_viewVersion=1.23+htslib-1.23
##bcftools_viewCommand=view /tmp/hg00150_apoe/merge_output.vcf.gz; Date=Mon Jun  8
    12:57:46 2026
#CHROM    POS    ID    REF    ALT    QUAL    FILTER    INFO    FORMAT    HG00150
chr19    44908684    .    T    C    24.68    PASS    P    GT:GQ:DP:AD:AF
    0/1:24:67:37,30:0.4478
chr19    44908822    .    C    T    21.57    PASS    P    GT:GQ:DP:AD:AF
    0/1:21:77:33,44:0.5714
```

**Turn 6 ▪ reasoning**

Let me analyze the results:
rs429358 at chr19:44908684 (ref T, alt C):
- Genotype: 0/1 (heterozygous) - one T allele, one C allele
- This means one allele has C ($\varepsilon$4-linked) and one has T

rs7412 at chr19:44908822 (ref C, alt T):
- Genotype: 0/1 (heterozygous) - one C allele, one T allele
- This means one allele has C and one has T

Now let me figure out the haplotypes/phasing. The genotypes are unphased (0/1 for both, no phasing information from Clair3). But we can deduce the diplotypes:
We have two SNPs, each heterozygous. The possible combinations:
The three common haplotypes:
- $\varepsilon$4 = (rs429358=C, rs7412=C)
- $\varepsilon$3 = (rs429358=T, rs7412=C)
- $\varepsilon$2 = (rs429358=T, rs7412=T)

The sample has:
- rs429358: T/C (one T, one C)
- rs7412: C/T (one C, one T)

Since rs429358 has one C, that C must come from an $\varepsilon$4 haplotype ($\varepsilon$4 is the only one with C at rs429358). So one haplotype is $\varepsilon$4.
The other haplotype has T at rs429358, which means it's either $\varepsilon$3 or $\varepsilon$2. The sample has both C and T at rs7412. If the $\varepsilon$4 chromosome has C at rs7412, then the other chromosome must have T at rs7412. That means the other haplotype is $\varepsilon$2 (T at rs429358, T at rs7412).
So the diplotype is $\varepsilon 4/\varepsilon 2$.
Wait, let me double-check:
- $\varepsilon$4 = (C, C)
- $\varepsilon$3 = (T, C)
- $\varepsilon$2 = (T, T)

The sample: rs429358: T/C, rs7412: C/T
If the haplotypes are ε4 + ε2:
- ε4: C at rs429358, C at rs7412
- ε2: T at rs429358, T at rs7412
- Combined: rs429358 = C/T , rs7412 = C/T
If the haplotypes were ε4 + ε3:
- ε4: C at rs429358, C at rs7412
- ε3: T at rs429358, C at rs7412
- Combined: rs429358 = C/T , rs7412 = C/C (but we see C/T)
So ε4/ε2 is the only combination that fits.
The conventional notation: ε4/ε2 or e4/e2. Let me use the format requested: e4/e2 or similar.
Looking at the example: 'ANSWER: e4/e4'
So: ANSWER: e4/e2

**Turn 6 ▪ tool call: run_shell**

```
run_shell:
# Let me also verify by looking at the pileup VCF more carefully
zcat /tmp/hg00150_apoe/merge_output.vcf.gz | grep -v "^#"
```

**Result ▪ run_shell**

```
returncode: 0
stdout:
chr19    44908684    .    T    C    24.68    PASS    P    GT:GQ:DP:AD:AF
    0/1:24:67:37,30:0.4478
chr19    44908822    .    C    T    21.57    PASS    P    GT:GQ:DP:AD:AF
    0/1:21:77:33,44:0.5714
```

**Turn 7 ▪ reasoning**

The results are clear:
rs429358 (chr19:44908684): ref T, alt C → genotype 0/1 (heterozygous T/C)
rs7412 (chr19:44908822): ref C, alt T → genotype 0/1 (heterozygous C/T)
The haplotypes:
- ε4 = (rs429358=C, rs7412=C)
- ε3 = (rs429358=T, rs7412=C)
- ε2 = (rs429358=T, rs7412=T)
With both SNPs heterozygous, the only combination that matches is ε4/ε2:
- ε4 contributes C at rs429358 and C at rs7412
- ε2 contributes T at rs429358 and T at rs7412
- Together: rs429358=T/C, rs7412=C/T
So the diplotype is ε4/ε2.
ANSWER: e4/e2

**Final answer**

The sample HG00150 is Illumina (PL:illumina in all @RG lines).
Clair3 called both SNPs as heterozygous:
- **rs429358** (chr19:44908684): '0/1' — T/C
- **rs7412** (chr19:44908822): '0/1' — C/T
Haplotype resolution:
- ε4 = (rs429358 C, rs7412 C)
- ε3 = (rs429358 T, rs7412 C)
- ε2 = (rs429358 T, rs7412 T)
The only combination yielding heterozygous calls at both sites is ε4 (C at 429358, C at 08822) paired with ε2 (T at 429358, T at 08822).
ANSWER: e4/e2

### Skills

#### Skills ▪ HG00136

12 turns, 13 tool calls, 267,764 tokens, 148.2 s; answer: e4/e4

*System prompt: §S1.1 (Skills); task prompt: §S1.2 (sample HG00136).*

**Turn 1 ▪ reasoning**

Let me work through this step by step.
1. First, determine the sequencing platform for HG00136 by inspecting the BAM header.
2. Then call variants and read genotypes at both SNP sites.
3. Combine into the diplotype.
Let me start by inspecting the BAM file.

##### Turn 1 ▪ tool call: run_shell

```
run_shell:
samtools view -H /autofs/nas5/xyu2/projects/clair3-api/clair3r/paper_experiments/.
    cache/apoe/HG00136_apoe.bam | grep "@PG"
```

##### Turn 1 ▪ tool call: run_shell

```
run_shell:
samtools view -H /autofs/nas5/xyu2/projects/clair3-api/clair3r/paper_experiments/.
    cache/apoe/HG00136_apoe.bam | grep "@RG"
```

##### Result ▪ run_shell

```
returncode: 0
stdout:
fs/internal/sequence/demux/ILLUMINA_NOVASEQ/SNA00297/181118_SNA00297_0047_BHFKLNDSXX
    /01.2018-11-20_11.12.53/CCDG_13607_B01_GRM_WGS/NYGC/Project_CCDG_13607_B01_GRM_WGS
    /Sample_HG00136/HG00136_CCAAGTCT-AAGGATGA_HFKLNDSXX_L003/fastq/HG00136_CCAAGTCT-
    AAGGATGA_HFKLNDSXX_L003_001.R1.fastq.gz /gpfs/internal/sequence/demux/
    ILLUMINA_NOVASEQ/SNA00297/181118_SNA00297_0047_BHFKLNDSXX/01.2018-11-20_11.12.53/
    CCDG_13607_B01_GRM_WGS/NYGC/Project_CCDG_13607_B01_GRM_WGS/Sample_HG00136/
    HG00136_CCAAGTCT-AAGGATGA_HFKLNDSXX_L003/fastq/HG00136_CCAAGTCT-
    AAGGATGA_HFKLNDSXX_L003_001.R2.fastq.gz
@PG     ID:bwa.9     PN:bwa     VN:0.7.15-r1140     CL:/data/NYGC/Software/bwa/bwa-0.7.15/
    bwa mem -Y -K 100000000 -t 16 -R @RG\tID:HG00136_CCAAGTCT-AAGGATGA_HFM5KDSXX_L004\
    tPL:illumina\tPM:Unknown\tLB:HG00136\tDS:GRCh38\tSM:HG00136\tCN:NYGenome\tPU:
    HFM5KDSXX.4.CCAAGTCT /gpfs/internal/sweng/production/Resources/GRCh38_1000genomes/
    GRCh38_full_analysis_set_plus_decoy_hla.fa /gpfs/internal/sequence/demux/
    ILLUMINA_NOVASEQ/SNA00297/181121_SNA00297_0048_AHFM5KDSXX/01.2018-11-26_15.18.32/
    CCDG_13607_B01_GRM_WGS/NYGC/Project_CCDG_13607_B01_GRM_WGS/Sample_HG00136/
    HG00136_CCAAGTCT-AAGGATGA_HFM5KDSXX_L004/fastq/HG00136_CCAAGTCT-
    AAGGATGA_HFM5KDSXX_L004_001.R1.fastq.gz /gpfs/internal/sequence/demux/
    ILLUMINA_NOVASEQ/SNA00297/181121_SNA00297_0048_AHFM5KDSXX/01.2018-11-26_15.18.32/
    CCDG_13607_B01_GRM_WGS/NYGC/Project_CCDG_13607_B01_GRM_WGS/Sample_HG00136/
    HG00136_CCAAGTCT-AAGGATGA_HFM5KDSXX_L004/fastq/HG00136_CCAAGTCT-
    AAGGATGA_HFM5KDSXX_L004_001.R2.fastq.gz
@PG     ID:bwa.A     PN:bwa     VN:0.7.15-r1140     CL:/data/NYGC/Software/bwa/bwa-0.7.15/
    bwa mem -Y -K 100000000 -t 16 -R @RG\tID:HG00136_CCAAGTCT-AAGGATGA_HFJFFDSXX_L003\
    tPL:illumina\tPM:Unknown\tLB:HG00136\tDS:GRCh38\tSM:HG00136\tCN:NYGenome\tPU:
    HFJFFDSXX.3.CCAAGTCT /gpfs/internal/sweng/production/Resources/GRCh38_1000genomes/
    GRCh38_full_analysis_set_plus_decoy_hla.fa /gpfs/internal/sequence/demux/
```

```
    ILLUMINA_NOVASEQ/SNA00297/181121_SNA00297_0049_BHFJFFDSXX/01.2018-11-23_11.09.41/
    CCDG_13607_B01_GRM_WGS/NYGC/Project_CCDG_13607_B01_GRM_WGS/Sample_HG00136/
    HG00136_CCAAGTCT-AAGGATGA_HFJFFDSXX_L003/fastq/HG00136_CCAAGTCT-
    AAGGATGA_HFJFFDSXX_L003_001.R1.fastq.gz /gpfs/internal/sequence/demux/
    ILLUMINA_NOVASEQ/SNA00297/181121_SNA00297_0049_BHFJFFDSXX/01.2018-11-23_11.09.41/
    CCDG_13607_B01_GRM_WGS/NYGC/Project_CCDG_13607_B01_GRM_WGS/Sample_HG00136/
    HG00136_CCAAGTCT-AAGGATGA_HFJFFDSXX_L003/fastq/HG00136_CCAAGTCT-
    AAGGATGA_HFJFFDSXX_L003_001.R2.fastq.gz
@PG    ID:bwa.B    PN:bwa    VN:0.7.15-r1140    CL:/data/NYGC/Software/bwa/bwa-0.7.15/
    bwa mem -Y -K 100000000 -t 16 -R @RG\tID:HG00136_CCAAGTCT-AAGGATGA_HFM5KDSXX_L002\
    tPL:illumina\tPM:Unknown\tLB:HG00136\tDS:GRCh38\tSM:HG00136\tCN:NYGenome\tPU:
    HFM5KDSXX.2.CCAAGTCT /gpfs/internal/sweng/production/Resources/GRCh38_1000genomes/
    GRCh38_full_analysis_set_plus_decoy_hla.fa /gpfs/internal/sequence/demux/
    ILLUMINA_NOVASEQ/SNA00297/181121_SNA00297_0048_AHFM5KDSXX/01.2018-11-26_15.18.32/
    CCDG_13607_B01_GRM_WGS/NYGC/Project_CCDG_13607_B01_GRM_WGS/Sample_HG00136/
    HG00136_CCAAGTCT-AAGGATGA_HFM5KDSXX_L002/fastq/HG00136_CCAAGTCT-
    AAGGATGA_HFM5KDSXX_L002_001.R1.fastq.gz /gpfs/internal/sequence/demux/
    ILLUMINA_NOVASEQ/SNA00297/181121_SNA00297_0048_AHFM5KDSXX/01.2018-11-26_15.18.32/
    CCDG_13607_B01_GRM_WGS/NYGC/Project_CCDG_13607_B01_GRM_WGS/Sample_HG00136/
    HG00136_CCAAGTCT-AAGGATGA_HFM5KDSXX_L002/fastq/HG00136_CCAAGTCT-
    AAGGATGA_HFM5KDSXX_L002_001.R2.fastq.gz
@PG    ID:GATK PrintReads    VN:3.5-0-g36282e4    CL:readGroup=null platform=null
    number=-1 sample_file=[] sample_name=[] simplify=false no_pg_tag=false
@PG    ID:samtools    PN:samtools    PP:GATK PrintReads    VN:1.23    CL:samtools view
     -T /autofs/bal19/zxzheng/testData/ilmn/data/
    GRCh38_full_analysis_set_plus_decoy_hla.fa -b -o paper_experiments/.cache/apoe/
    HG00136_apoe.bam https://ftp.sra.ebi.ac.uk/vol1/run/ERR324/ERR3240142/HG00136.
    final.cram chr19:44905000-44912000
@PG    ID:samtools.1    PN:samtools    PP:samtools    VN:1.22.1    CL:samtools view -H
     /autofs/nas5/xyu2/projects/clair3-api/clair3r/paper_experiments/.cache/apoe/
    HG00136_apoe.bam
```

**Result ▪ run_shell**

```
returncode: 0
stdout:
607_B01_GRM_WGS/NYGC/Project_CCDG_13607_B01_GRM_WGS/Sample_HG00136/HG00136_CCAAGTCT-
    AAGGATGA_HFM5KDSXX_L004/fastq/HG00136_CCAAGTCT-AAGGATGA_HFM5KDSXX_L004_001.R1.
    fastq.gz /gpfs/internal/sequence/demux/ILLUMINA_NOVASEQ/SNA00297/181121
    _SNA00297_0048_AHFM5KDSXX/01.2018-11-26_15.18.32/CCDG_13607_B01_GRM_WGS/NYGC/
    Project_CCDG_13607_B01_GRM_WGS/Sample_HG00136/HG00136_CCAAGTCT-
    AAGGATGA_HFM5KDSXX_L004/fastq/HG00136_CCAAGTCT-AAGGATGA_HFM5KDSXX_L004_001.R2.
    fastq.gz
@PG    ID:bwa.A    PN:bwa    VN:0.7.15-r1140    CL:/data/NYGC/Software/bwa/bwa-0.7.15/
    bwa mem -Y -K 100000000 -t 16 -R @RG\tID:HG00136_CCAAGTCT-AAGGATGA_HFJFFDSXX_L003\
    tPL:illumina\tPM:Unknown\tLB:HG00136\tDS:GRCh38\tSM:HG00136\tCN:NYGenome\tPU:
    HFJFFDSXX.3.CCAAGTCT /gpfs/internal/sweng/production/Resources/GRCh38_1000genomes/
    GRCh38_full_analysis_set_plus_decoy_hla.fa /gpfs/internal/sequence/demux/
    ILLUMINA_NOVASEQ/SNA00297/181121_SNA00297_0049_BHFJFFDSXX/01.2018-11-23_11.09.41/
    CCDG_13607_B01_GRM_WGS/NYGC/Project_CCDG_13607_B01_GRM_WGS/Sample_HG00136/
    HG00136_CCAAGTCT-AAGGATGA_HFJFFDSXX_L003/fastq/HG00136_CCAAGTCT-
    AAGGATGA_HFJFFDSXX_L003_001.R1.fastq.gz /gpfs/internal/sequence/demux/
    ILLUMINA_NOVASEQ/SNA00297/181121_SNA00297_0049_BHFJFFDSXX/01.2018-11-23_11.09.41/
    CCDG_13607_B01_GRM_WGS/NYGC/Project_CCDG_13607_B01_GRM_WGS/Sample_HG00136/
    HG00136_CCAAGTCT-AAGGATGA_HFJFFDSXX_L003/fastq/HG00136_CCAAGTCT-
    AAGGATGA_HFJFFDSXX_L003_001.R2.fastq.gz
@PG    ID:bwa.B    PN:bwa    VN:0.7.15-r1140    CL:/data/NYGC/Software/bwa/bwa-0.7.15/
    bwa mem -Y -K 100000000 -t 16 -R @RG\tID:HG00136_CCAAGTCT-AAGGATGA_HFM5KDSXX_L002\
```

```
    tPL:illumina\tPM:Unknown\tLB:HG00136\tDS:GRCh38\tSM:HG00136\tCN:NYGenome\tPU:
    HFM5KDSXX.2.CCAAGTCT /gpfs/internal/sweng/production/Resources/GRCh38_1000genomes/
    GRCh38_full_analysis_set_plus_decoy_hla.fa /gpfs/internal/sequence/demux/
    ILLUMINA_NOVASEQ/SNA00297/181121_SNA00297_0048_AHFM5KDSXX/01.2018-11-26_15.18.32/
    CCDG_13607_B01_GRM_WGS/NYGC/Project_CCDG_13607_B01_GRM_WGS/Sample_HG00136/
    HG00136_CCAAGTCT-AAGGATGA_HFM5KDSXX_L002/fastq/HG00136_CCAAGTCT-
    AAGGATGA_HFM5KDSXX_L002_001.R1.fastq.gz /gpfs/internal/sequence/demux/
    ILLUMINA_NOVASEQ/SNA00297/181121_SNA00297_0048_AHFM5KDSXX/01.2018-11-26_15.18.32/
    CCDG_13607_B01_GRM_WGS/NYGC/Project_CCDG_13607_B01_GRM_WGS/Sample_HG00136/
    HG00136_CCAAGTCT-AAGGATGA_HFM5KDSXX_L002/fastq/HG00136_CCAAGTCT-
    AAGGATGA_HFM5KDSXX_L002_001.R2.fastq.gz
@RG    ID:HG00136_CCAAGTCT-AAGGATGA_HFJFFDSXX_L001    PL:illumina    PM:Unknown    LB:
    HG00136    DS:GRCh38    SM:HG00136    CN:NYGenome    PU:HFJFFDSXX.1.CCAAGTCT
@RG    ID:HG00136_CCAAGTCT-AAGGATGA_HFJFFDSXX_L002    PL:illumina    PM:Unknown    LB:
    HG00136    DS:GRCh38    SM:HG00136    CN:NYGenome    PU:HFJFFDSXX.2.CCAAGTCT
@RG    ID:HG00136_CCAAGTCT-AAGGATGA_HFJFFDSXX_L003    PL:illumina    PM:Unknown    LB:
    HG00136    DS:GRCh38    SM:HG00136    CN:NYGenome    PU:HFJFFDSXX.3.CCAAGTCT
@RG    ID:HG00136_CCAAGTCT-AAGGATGA_HFJFFDSXX_L004    PL:illumina    PM:Unknown    LB:
    HG00136    DS:GRCh38    SM:HG00136    CN:NYGenome    PU:HFJFFDSXX.4.CCAAGTCT
@RG    ID:HG00136_CCAAGTCT-AAGGATGA_HFKLNDSXX_L001    PL:illumina    PM:Unknown    LB:
    HG00136    DS:GRCh38    SM:HG00136    CN:NYGenome    PU:HFKLNDSXX.1.CCAAGTCT
@RG    ID:HG00136_CCAAGTCT-AAGGATGA_HFKLNDSXX_L002    PL:illumina    PM:Unknown    LB:
    HG00136    DS:GRCh38    SM:HG00136    CN:NYGenome    PU:HFKLNDSXX.2.CCAAGTCT
@RG    ID:HG00136_CCAAGTCT-AAGGATGA_HFKLNDSXX_L003    PL:illumina    PM:Unknown    LB:
    HG00136    DS:GRCh38    SM:HG00136    CN:NYGenome    PU:HFKLNDSXX.3.CCAAGTCT
@RG    ID:HG00136_CCAAGTCT-AAGGATGA_HFKLNDSXX_L004    PL:illumina    PM:Unknown    LB:
    HG00136    DS:GRCh38    SM:HG00136    CN:NYGenome    PU:HFKLNDSXX.4.CCAAGTCT
@RG    ID:HG00136_CCAAGTCT-AAGGATGA_HFM5KDSXX_L001    PL:illumina    PM:Unknown    LB:
    HG00136    DS:GRCh38    SM:HG00136    CN:NYGenome    PU:HFM5KDSXX.1.CCAAGTCT
@RG    ID:HG00136_CCAAGTCT-AAGGATGA_HFM5KDSXX_L002    PL:illumina    PM:Unknown    LB:
    HG00136    DS:GRCh38    SM:HG00136    CN:NYGenome    PU:HFM5KDSXX.2.CCAAGTCT
@RG    ID:HG00136_CCAAGTCT-AAGGATGA_HFM5KDSXX_L003    PL:illumina    PM:Unknown    LB:
    HG00136    DS:GRCh38    SM:HG00136    CN:NYGenome    PU:HFM5KDSXX.3.CCAAGTCT
@RG    ID:HG00136_CCAAGTCT-AAGGATGA_HFM5KDSXX_L004    PL:illumina    PM:Unknown    LB:
    HG00136    DS:GRCh38    SM:HG00136    CN:NYGenome    PU:HFM5KDSXX.4.CCAAGTCT
```

**Turn 2 ▪ reasoning**

Platform: Illumina (NovaSeq), bwa-mem alignment. So I need the ‘ilmn‘ model. The BAM is already limited to chr19:44905000-44912000 — this is great, it’s a small region around our SNPs.
Now I need to:
1. Create a BED file for the region of interest (the two SNPs)
2. Run Clair3 with the ilmn model
3. Read genotypes from the output VCF
Let me first check that the BAM has an index and the ref file.
The SNPs:
- rs429358: chr19:44908684 (1-based), ref T, alt C
- rs7412: chr19:44908822 (1-based), ref C, alt T
I need to create a BED region covering both SNPs. Let me make it broad enough for Clair3’s pileup window. Let’s say chr19:44908000-44909500.
Actually, Clair3 takes a BED file for calling regions. Let me create one covering both SNPs, and then run the caller.
Let me check the ref and bam index first.

**Turn 2 ▪ tool call: run_shell**

```
run_shell:
ls /autofs/nas5/xyu2/projects/clair3-api/clair3r/paper_experiments/.cache/apoe/
```

```
    HG00136_apoe.bam.bai 2>/dev/null || samtools index /autofs/nas5/xyu2/projects/
    clair3-api/clair3r/paper_experiments/.cache/apoe/HG00136_apoe.bam
```

**Turn 2 ▪ tool call: run_shell**

```
run_shell:
ls /autofs/bal19/zxzheng/testData/ilmn/data/GRCh38_full_analysis_set_plus_decoy_hla.fa
     /autofs/bal19/zxzheng/testData/ilmn/data/GRCh38_full_analysis_set_plus_decoy_hla.
    fa.fai
```

**Result ▪ run_shell**

```
returncode: 0
stdout:
/autofs/nas5/xyu2/projects/clair3-api/clair3r/paper_experiments/.cache/apoe/
    HG00136_apoe.bam.bai
```

**Result ▪ run_shell**

```
returncode: 0
stdout:
/autofs/bal19/zxzheng/testData/ilmn/data/GRCh38_full_analysis_set_plus_decoy_hla.fa
/autofs/bal19/zxzheng/testData/ilmn/data/GRCh38_full_analysis_set_plus_decoy_hla.fa.
    fai
```

**Turn 3 ▪ reasoning**

Good, BAM index and ref + index all exist. Now create the BED file and run Clair3.

**Turn 3 ▪ tool call: run_shell**

```
run_shell:
printf "chr19\t44908000\t44909500\n" > /tmp/apoe_region.bed
cat /tmp/apoe_region.bed
```

**Result ▪ run_shell**

```
returncode: 0
stdout:
chr19    44908000    44909500
```

**Turn 4 ▪ reasoning**

Now run Clair3 with the ilmn model.

**Turn 4 ▪ tool call: run_shell**

```
run_shell:
conda run -n clair3-torch python /autofs/nas5/xyu2/projects/clair3-api/Clair3/
    run_clair3.py \
  --bam_fn /autofs/nas5/xyu2/projects/clair3-api/clair3r/paper_experiments/.cache/apoe
    /HG00136_apoe.bam \
  --ref_fn /autofs/bal19/zxzheng/testData/ilmn/data/
    GRCh38_full_analysis_set_plus_decoy_hla.fa \
  --model_path /autofs/nas5/xyu2/softwares/clair3_models_pytorch/ilmn \
  --platform ilmn \
  --threads 8 \
```

```
  --output /tmp/clair3_hg00136_apoe \
  --bed_fn /tmp/apoe_region.bed \
  --device cpu \
  --longphase /autofs/nas5/xyu2/projects/clair3-api/Clair3/longphase \
  --sample_name HG00136 2>&1 | tail -30
```

Result ▪ run_shell

```
returncode: 0
stdout:
[INFO] 4/7 Haplotag input BAM file using Whatshap
Found 1 sample(s) in input VCF
Found 1 sample(s) in BAM file
Number of supplementary alignments: 0
Number of non-singleton groups: 0
Skipped 0 groups
Found 0 reads covering 0 variants

== SUMMARY ==
Total alignments processed:                      1978
Alignments that could be tagged:                    0
Alignments spanning multiple phase sets:            0
Finished in 0.9 s

[INFO] 5/7 Select candidates for full-alignment calling
[93m[WARNING] Cannot find any low-quality 0/1 or 1/1 variant in pileup output using
    variant quality cut-off proportion: 0.3, total variants: 1[0m
[93m[WARNING] Set low variant quality score cut-off to 0.0[0m
[INFO] Set variants quality cutoff 0.0
[INFO] Set reference calls quality cutoff 30.0
[93m[WARNING] Cannot find any low-quality 0/0, 0/1 or 1/1 variant in pileup output in
    contig chr19[0m
[93m[WARNING] Copying pileup.vcf.gz to /tmp/clair3_hg00136_apoe/merge_output.vcf.gz[0m
[INFO] Low quality reference calls to be processed in chr19: 0
[INFO] Low quality variants to be processed in chr19: 0

[INFO] 6/7 Call low-quality variants using full-alignment model
[INFO] Delay 2 seconds before starting variant calling ...
[93m[WARNING] Skip full-alignment variant calling for empty full-alignment regions[0m
[93m[WARNING] No vcf file found with prefix:/tmp/clair3_hg00136_apoe/tmp/
    full_alignment_output/full_alignment, output empty vcf file[0m
[INFO] Exit in full-alignment variant calling
```

**Turn 5 ▪ reasoning**

The run completed. Let me check the output VCF for the genotypes at the two SNP positions.

**Turn 5 ▪ tool call: run_shell**

```
run_shell:
ls -la /tmp/clair3_hg00136_apoe/merge_output.vcf.gz && bcftools view -H /tmp/
    clair3_hg00136_apoe/merge_output.vcf.gz
```

Result ▪ run_shell

```
returncode: 0
stdout:
```

```
-rw-r--r-- 1 xyu2 users 18175 Jun  8 12:53 /tmp/clair3_hg00136_apoe/merge_output.vcf.
    gz
chr19    44908475    .    T    .    29.81    RefCall    P    GT:GQ:DP:AD:AF
    0/0:29:42:36:0.8571
chr19    44908484    .    T    .    37.29    RefCall    P    GT:GQ:DP:AD:AF
    0/0:37:46:37:0.8043
chr19    44908501    .    T    .    36.9    RefCall    P    GT:GQ:DP:AD:AF
    0/0:36:47:40:0.8511
chr19    44908513    .    T    .    34.85    RefCall    P    GT:GQ:DP:AD:AF
    0/0:34:46:40:0.8696
chr19    44908521    .    T    .    37.59    RefCall    P    GT:GQ:DP:AD:AF
    0/0:37:49:42:0.8571
chr19    44908523    .    T    .    33.47    RefCall    P    GT:GQ:DP:AD:AF
    0/0:33:48:43:0.8958
chr19    44908684    .    T    C    30.89    PASS    P    GT:GQ:DP:AD:AF
    1/1:30:59:0,59:1
```

**Turn 6 ▪ reasoning**

I see rs429358 at chr19:44908684: genotype is 1/1 (homozygous alt C/C).
Now I need to check rs7412 at chr19:44908822. I don't see it in the output. Let me check if it's in the full VCF, maybe I need to look more specifically — the output might be truncated. Let me also check the pileup VCF.

**Turn 6 ▪ tool call: run_shell**

```
run_shell:
bcftools view -H -r chr19:44908800-44908850 /tmp/clair3_hg00136_apoe/merge_output.vcf.
    gz 2>/dev/null || echo "No variant at rs7412"
# Also check the pileup VCF
for f in /tmp/clair3_hg00136_apoe/tmp/*/pileup.vcf.gz; do echo "=== $f ==="; bcftools
    view -H -r chr19:44908822 "$f" 2>/dev/null || echo "(no call)"; done
```

**Result ▪ run_shell**

```
returncode: 0
stdout:
No variant at rs7412
=== /tmp/clair3_hg00136_apoe/tmp/*/pileup.vcf.gz ===
(no call)
```

**Turn 7 ▪ reasoning**

rs7412 isn't called by Clair3. The output only went to chr19:44908475-44908684. But the BED was 44908000-44909500, so rs7412 at 44908822 should be covered. Let me check if it's in the pileup output or if there's a RefCall. Let me look at all records in the merge output.

**Turn 7 ▪ tool call: run_shell**

```
run_shell:
bcftools view -H /tmp/clair3_hg00136_apoe/merge_output.vcf.gz | grep -c "^"
echo "---"
bcftools view -H /tmp/clair3_hg00136_apoe/merge_output.vcf.gz | awk '{print $1,$2,$4,
    $5,$6,$7,$10}'
```

**Result ▪ run_shell**

```
returncode: 0
stdout:
7
---
chr19 44908475 T . 29.81 RefCall 0/0:29:42:36:0.8571
chr19 44908484 T . 37.29 RefCall 0/0:37:46:37:0.8043
chr19 44908501 T . 36.9 RefCall 0/0:36:47:40:0.8511
chr19 44908513 T . 34.85 RefCall 0/0:34:46:40:0.8696
chr19 44908521 T . 37.59 RefCall 0/0:37:49:42:0.8571
chr19 44908523 T . 33.47 RefCall 0/0:33:48:43:0.8958
chr19 44908684 T C 30.89 PASS 1/1:30:59:0,59:1
```

**Turn 8 ▪ reasoning**

The VCF only has 7 records, and rs7412 at 44908822 is not among them. This is strange. The BED was 44908000-44909500 which covers both SNPs. Let me check if it's in the pileup.vcf.gz file, which might have all positions.
Let me find where the pileup vcf is.

**Turn 8 ▪ tool call: run_shell**

```
run_shell:
find /tmp/clair3_hg00136_apoe -name "pileup.vcf.gz" 2>/dev/null
```

**Result ▪ run_shell**

```
returncode: 0
stdout:
/tmp/clair3_hg00136_apoe/pileup.vcf.gz
```

**Turn 9 ▪ reasoning**

Let me check the pileup VCF for rs7412.

**Turn 9 ▪ tool call: run_shell**

```
run_shell:
bcftools view -H -r chr19:44908820-44908825 /tmp/clair3_hg00136_apoe/pileup.vcf.gz
```

**Result ▪ run_shell**

```
returncode: 0
```

**Turn 10 ▪ reasoning**

No call at rs7412 at all, even in the pileup VCF. Let me understand why. Let me check the full pileup VCF to see what positions were covered.
Actually, wait. Let me think about this more carefully. The warning said:
"Only 1 low-quality variant was found" and "Cannot find any low-quality 0/1 or 1/1 variant" — so Clair3 found only one variant (rs429358 at 44908684, which was called as 1/1) and no other candidates.
The fact that rs7412 is not even in the pileup means Clair3's pileup step didn't consider that position at all, or the position was so clearly homozygous reference that it wasn't even reported.
Let me look at what's in the pileup VCF more broadly.

**Turn 10 ▪ tool call: run_shell**

```
run_shell:
bcftools view -H /tmp/clair3_hg00136_apoe/pileup.vcf.gz | awk '{print $1,$2,$4,$5,$10
    }' | head -20
```

**Result ▪ run_shell**

```
returncode: 0
stdout:
chr19 44908475 T . 0/0:29:42:36:0.8571
chr19 44908484 T . 0/0:37:46:37:0.8043
chr19 44908501 T . 0/0:36:47:40:0.8511
chr19 44908513 T . 0/0:34:46:40:0.8696
chr19 44908521 T . 0/0:37:49:42:0.8571
chr19 44908523 T . 0/0:33:48:43:0.8958
chr19 44908684 T C 1/1:30:59:0,59:1
```

**Turn 11 ▪ reasoning**

The pileup VCF only has 7 records and none at rs7412 (44908822). This means Clair3's pileup model considered the position at chr19:44908822 to be so clearly homozygous reference that it wasn't even output as a low-quality candidate.
This is actually expected behavior for Illumina data - some positions are filtered out by Clair3's pileup model if the evidence strongly supports reference. Since rs7412 wasn't called as a variant, it means the genotype is homozygous reference: C/C at that position.
So let me verify by checking the BAM directly at rs7412 to confirm the genotype.

**Turn 11 ▪ tool call: run_shell**

```
run_shell:
# rs429358: chr19:44908684  (ref T, alt C)
echo "=== rs429358 (chr19:44908684) ==="
samtools mpileup -r chr19:44908684-44908684 -f /autofs/bal19/zxzheng/testData/ilmn/
    data/GRCh38_full_analysis_set_plus_decoy_hla.fa /autofs/nas5/xyu2/projects/clair3-
    api/clair3r/paper_experiments/.cache/apoe/HG00136_apoe.bam

echo ""
echo "=== rs7412 (chr19:44908822) ==="
samtools mpileup -r chr19:44908822-44908822 -f /autofs/bal19/zxzheng/testData/ilmn/
    data/GRCh38_full_analysis_set_plus_decoy_hla.fa /autofs/nas5/xyu2/projects/clair3-
    api/clair3r/paper_experiments/.cache/apoe/HG00136_apoe.bam
```

**Result ▪ run_shell**

```
returncode: 0
stdout:
=== rs429358 (chr19:44908684) ===
chr19    44908684    T    50    CcCCcCCcCCcCCcCCCCcccCcCCcccCcCCCcCCcccCcccCcCCcCC
    ??????????????????????????????????????????????????

=== rs7412 (chr19:44908822) ===
chr19    44908822    C    52    ..,.,.,.,,.,..,,,.,,,,.,,,.,.,...,.,..,..,.....,...,
      ????????????????????????????????????????????????????
stderr:
[mpileup] 1 samples in 1 input files
[mpileup] 1 samples in 1 input files
```

**Turn 12 ▪ reasoning**

Now I have the full picture:
**rs429358 (chr19:44908684)**:
- Ref = T
- Mpileup shows all C bases (50 reads, all C — "CcCCcCCcCCcCCcCCCCcccCcCCcccCcCCCcCCc-ccCcccCcCCcCC")
- So genotype is C/C (homozygous alternate)
**rs7412 (chr19:44908822)**:
- Ref = C
- Mpileup shows all reference matches (52 reads, all "." or "," - no variant base)
- So genotype is C/C (homozygous reference)
Now let's determine the haplotypes:
Each chromosome has two haplotypes (one from each parent), determined by the combination of alleles at rs429358 and rs7412:
- $\varepsilon 4$ = (rs429358=C, rs7412=C)
- $\varepsilon 3$ = (rs429358=T, rs7412=C)
- $\varepsilon 2$ = (rs429358=T, rs7412=T)
At rs429358: genotype is C/C → both chromosomes have C at rs429358
At rs7412: genotype is C/C → both chromosomes have C at rs7412
So each chromosome is (C, C) = $\varepsilon 4/\varepsilon 4$.
ANSWER: e4/e4

**Final answer**

Both SNPs are now clear:
- **rs429358** (chr19:44908684): 50 reads, all C → genotype **C/C** (homozygous alt)
- **rs7412** (chr19:44908822): 52 reads, all reference C → genotype **C/C** (homozygous ref)
Haplotype resolution: both chromosomes carry (rs429358=C, rs7412=C) = **$\varepsilon 4/\varepsilon 4$**.
ANSWER: e4/e4

#### Skills ▪ HG00150

10 turns, 9 tool calls, 205,487 tokens, 82.9 s; answer: e2/e4

*System prompt: §S1.1 (Skills); task prompt: §S1.2 (sample HG00150).*

**Turn 1 ▪ reasoning**

The user wants me to determine the APOE diplotype for sample HG00136. Let me follow the steps:
1. First, I need to check the BAM file to determine the sequencing platform
2. Then call variants using Clair3 with the appropriate model
3. Read the genotypes at the two SNP positions
4. Combine into the diplotype
Let me start by inspecting the BAM file for HG00136.
Wait, let me re-read. The user says HG00136 in the registered samples but the question header says "HG00150". Let me look more carefully...
The registered samples listed are:
- HG00136: bam=/autofs/nas5/xyu2/projects/clair3-api/clair3r/paper_experiments/.cache/apoe/HG00136_apoe.bam
- HG00150: bam=/autofs/nas5/xyu2/projects/clair3-api/clair3r/paper_experiments/.cache/apoe/HG00150_apoe.bam
The user says "Sample HG00150 is a human whole-genome sequencing dataset." So it's HG00150.
Let me first inspect the BAM header for HG00150 to determine the platform.

**Turn 1 ▪ tool call: run_shell**

```
run_shell:
samtools view -H /autofs/nas5/xyu2/projects/clair3-api/clair3r/paper_experiments/.
    cache/apoe/HG00150_apoe.bam 2>&1 | head -30
```

**Result ▪ run_shell**

```
returncode: 0
stdout:
C/Project_CCDG_13607_B01_GRM_WGS/Sample_HG00150/HG00150_CATAGAGT-
    TGCCACCA_HFH25DSXX_L002/fastq/HG00150_CATAGAGT-TGCCACCA_HFH25DSXX_L002_001.R2.
    fastq.gz
@PG    ID:bwa.B    PN:bwa    VN:0.7.15-r1140    CL:/data/NYGC/Software/bwa/bwa-0.7.15/
    bwa mem -Y -K 100000000 -t 16 -R @RG\tID:HG00150_CATAGAGT-TGCCACCA_HFKWWDSXX_L002\
    tPL:illumina\tPM:Unknown\tLB:HG00150\tDS:GRCh38\tSM:HG00150\tCN:NYGenome\tPU:
    HFKWWDSXX.2.CATAGAGT /gpfs/internal/sweng/production/Resources/GRCh38_1000genomes/
    GRCh38_full_analysis_set_plus_decoy_hla.fa /gpfs/internal/sequence/demux/
    ILLUMINA_NOVASEQ/SNA00297/181026_SNA00297_0035_BHFKWWDSXX/01.2018-10-28_12.05.06/
    CCDG_13607_B01_GRM_WGS/NYGC/Project_CCDG_13607_B01_GRM_WGS/Sample_HG00150/
    HG00150_CATAGAGT-TGCCACCA_HFKWWDSXX_L002/fastq/HG00150_CATAGAGT-
    TGCCACCA_HFKWWDSXX_L002_001.R1.fastq.gz /gpfs/internal/sequence/demux/
    ILLUMINA_NOVASEQ/SNA00297/181026_SNA00297_0035_BHFKWWDSXX/01.2018-10-28_12.05.06/
    CCDG_13607_B01_GRM_WGS/NYGC/Project_CCDG_13607_B01_GRM_WGS/Sample_HG00150/
    HG00150_CATAGAGT-TGCCACCA_HFKWWDSXX_L002/fastq/HG00150_CATAGAGT-
    TGCCACCA_HFKWWDSXX_L002_001.R2.fastq.gz
@PG    ID:GATK PrintReads    VN:3.5-0-g36282e4    CL:readGroup=null platform=null
    number=-1 sample_file=[] sample_name=[] simplify=false no_pg_tag=false
@PG    ID:samtools    PN:samtools    PP:GATK PrintReads    VN:1.23    CL:/autofs/nas5/
    xyu2/softwares/miniforge3/envs/clair3-torch/bin/samtools view -T /autofs/bal19/
    zxzheng/testData/ilmn/data/GRCh38_full_analysis_set_plus_decoy_hla.fa -b -o /
    autofs/nas5/xyu2/projects/clair3-api/clair3r/paper_experiments/.cache/apoe/
    HG00150_apoe.bam https://ftp.sra.ebi.ac.uk/vol1/run/ERR324/ERR3240154/HG00150.
    final.cram chr19:44903000-44913000
@PG    ID:samtools.1    PN:samtools    PP:samtools    VN:1.22.1    CL:samtools view -H
     /autofs/nas5/xyu2/projects/clair3-api/clair3r/paper_experiments/.cache/apoe/
    HG00150_apoe.bam
@SQ    SN:chr1    LN:248956422    M5:6aef897c3d6ff0c78aff06ac189178dd    UR:/gpfs/
    internal/sweng/production/Resources/GRCh38_1000genomes/
    GRCh38_full_analysis_set_plus_decoy_hla.fa
@SQ    SN:chr2    LN:242193529    M5:f98db672eb0993dcfdabafe2a882905c    UR:/gpfs/
    internal/sweng/production/Resources/GRCh38_1000genomes/
    GRCh38_full_analysis_set_plus_decoy_hla.fa
@SQ    SN:chr3    LN:198295559    M5:76635a41ea913a405ded820447d067b0    UR:/gpfs/
    internal/sweng/production/Resources/GRCh38_1000genomes/
    GRCh38_full_analysis_set_plus_decoy_hla.fa
@SQ    SN:chr4    LN:190214555    M5:3210fecf1eb92d5489da4346b3fddc6e    UR:/gpfs/
    internal/sweng/production/Resources/GRCh38_1000genomes/
    GRCh38_full_analysis_set_plus_decoy_hla.fa
@SQ    SN:chr5    LN:181538259    M5:a811b3dc9fe66af729dc0dddf7fa4f13    UR:/gpfs/
    internal/sweng/production/Resources/GRCh38_1000genomes/
    GRCh38_full_analysis_set_plus_decoy_hla.fa
@SQ    SN:chr6    LN:170805979    M5:5691468a67c7e7a7b5f2a3a683792c29    UR:/gpfs/
    internal/sweng/production/Resources/GRCh38_1000genomes/
    GRCh38_full_analysis_set_plus_decoy_hla.fa
@SQ    SN:chr7    LN:159345973    M5:cc044cc2256a1141212660fb07b6171e    UR:/gpfs/
    internal/sweng/production/Resources/GRCh38_1000genomes/
    GRCh38_full_analysis_set_plus_decoy_hla.fa
@SQ    SN:chr8    LN:145138636    M5:c67955b5f7815a9a1edfaa15893d3616    UR:/gpfs/
    internal/sweng/production/Resources/GRCh38_1000genomes/
    GRCh38_full_analysis_set_plus_decoy_hla.fa
@SQ    SN:chr9    LN:138394717    M5:6c198acf68b5af7b9d676dfdd531b5de    UR:/gpfs/
    internal/sweng/production/Resources/GRCh38_1000genomes/
    GRCh38_full_analysis_set_plus_decoy_hla.fa
```

```
@SQ	SN:chr10	LN:133797422	M5:c0eeee7acfdaf31b770a509bdaa6e51a	UR:/gpfs/
    internal/sweng/production/Resources/GRCh38_1000genomes/
    GRCh38_full_analysis_set_plus_decoy_hla.fa
@SQ	SN:chr11	LN:135086622	M5:1511375dc2dd1b633af8cf439ae90cec	UR:/gpfs/
    internal/sweng/production/Resources/GRCh38_1000genomes/
    GRCh38_full_analysis_set_plus_decoy_hla.fa
@SQ	SN:chr12	LN:133275309	M5:96e414eace405d8c27a6d35ba19df56f	UR:/gpfs/
    internal/sweng/production/Resources/GRCh38_1000genomes/
    GRCh38_full_analysis_set_plus_decoy_hla.fa
@SQ	SN:chr13	LN:114364328	M5:a5437debe2ef9c9ef8f3ea2874ae1d82	UR:/gpfs/
    internal/sweng/production/Resources/GRCh38_1000genomes/
    GRCh38_full_analysis_set_plus_decoy_hla.fa
```

**Turn 2 ▪ reasoning**

From the BAM header I can see:
- Platform: Illumina (bwa mem with PL:illumina tag, and "ILLUMINA_NOVASEQ")
- Aligner: bwa
- Reference: GRCh38_full_analysis_set_plus_decoy_hla.fa
- The BAM was created from a CRAM file for chr19:44903000-44913000 region (APOE region)

Since it's Illumina, I should use the 'ilmn' model. The reference is already specified at '/autofs/bal19/zxzheng/testData/ilmn/data/GRCh38_full_analysis_set_plus_decoy_hla.fa'.
Now let me create the output directory and run Clair3 with the ilmn model. The SNPs are at:
- rs429358: chr19:44908684
- rs7412: chr19:44908822

I need to create a BED file covering these positions. Let me create a region BED.
Let me also check if the BAM has an index.

**Turn 2 ▪ tool call: run_shell**

```
run_shell:
ls /autofs/nas5/xyu2/projects/clair3-api/clair3r/paper_experiments/.cache/apoe/
    HG00150_apoe.bam* 2>&1
```

**Result ▪ run_shell**

```
returncode: 0
stdout:
/autofs/nas5/xyu2/projects/clair3-api/clair3r/paper_experiments/.cache/apoe/
    HG00150_apoe.bam
/autofs/nas5/xyu2/projects/clair3-api/clair3r/paper_experiments/.cache/apoe/
    HG00150_apoe.bam.bai
```

**Turn 3 ▪ reasoning**

Good, index exists. Platform is Illumina → use 'ilmn' model. Let me now:
1. Create a BED file covering chr19:44908684-44908822 (the region with the two SNPs)
2. Run Clair3
3. Extract genotypes

The reference is at '/autofs/bal19/zxzheng/testData/ilmn/data/GRCh38_full_analysis_set_plus_decoy_hla.fa'. Let me verify it has a .fai index.

**Turn 3 ▪ tool call: run_shell**

```
run_shell:
ls /autofs/bal19/zxzheng/testData/ilmn/data/GRCh38_full_analysis_set_plus_decoy_hla.fa
    * 2>&1
```

**Result ▪ run_shell**

```
returncode: 0
stdout:
/autofs/bal19/zxzheng/testData/ilmn/data/GRCh38_full_analysis_set_plus_decoy_hla.fa
/autofs/bal19/zxzheng/testData/ilmn/data/GRCh38_full_analysis_set_plus_decoy_hla.fa.
    amb
/autofs/bal19/zxzheng/testData/ilmn/data/GRCh38_full_analysis_set_plus_decoy_hla.fa.
    ann
/autofs/bal19/zxzheng/testData/ilmn/data/GRCh38_full_analysis_set_plus_decoy_hla.fa.
    bwt
/autofs/bal19/zxzheng/testData/ilmn/data/GRCh38_full_analysis_set_plus_decoy_hla.fa.
    fai
/autofs/bal19/zxzheng/testData/ilmn/data/GRCh38_full_analysis_set_plus_decoy_hla.fa.
    pac
/autofs/bal19/zxzheng/testData/ilmn/data/GRCh38_full_analysis_set_plus_decoy_hla.fa.sa
```

**Turn 4 ▪ reasoning**

Good. Now let me create the BED file and run Clair3. Let me create a BED covering the two SNP positions. I'll make a region that spans both SNPs comfortably.
chr19:44908684-44908822. Let me create a slightly wider region.
Let me also create the output directory and the BED file, then run Clair3.

**Turn 4 ▪ tool call: run_shell**

```
run_shell:
mkdir -p /tmp/hg00150_apoe && printf 'chr19\t44908600\t44908900\n' > /tmp/hg00150_apoe
    /region.bed && cat /tmp/hg00150_apoe/region.bed
```

**Result ▪ run_shell**

```
returncode: 0
stdout:
chr19    44908600    44908900
```

**Turn 5 ▪ reasoning**

Now run Clair3 with the ilmn model.

**Turn 5 ▪ tool call: run_shell**

```
run_shell:
cd /tmp/hg00150_apoe && conda run -n clair3-torch python /autofs/nas5/xyu2/projects/
    clair3-api/Clair3/run_clair3.py \
  --bam_fn /autofs/nas5/xyu2/projects/clair3-api/clair3r/paper_experiments/.cache/apoe
    /HG00150_apoe.bam \
  --ref_fn /autofs/bal19/zxzheng/testData/ilmn/data/
    GRCh38_full_analysis_set_plus_decoy_hla.fa \
  --model_path /autofs/nas5/xyu2/softwares/clair3_models_pytorch/ilmn \
  --platform ilmn \
  --threads 8 \
  --output /tmp/hg00150_apoe/clair3_output \
  --bed_fn /tmp/hg00150_apoe/region.bed \
  --device cpu \
  --longphase /autofs/nas5/xyu2/projects/clair3-api/Clair3/longphase \
  --sample_name HG00150 2>&1
```

**Result ▪ run_shell**

```
returncode: 0
stdout:
processed positions in chr19 (chunk 7/12) : 0
Total time elapsed: 0.00 s
[INFO] No vcf output for file /tmp/hg00150_apoe/clair3_output/tmp/pileup_output/
    pileup_chr19_7.vcf, remove empty file
[INFO] Delay 3 seconds before starting variant calling ...
[mpileup] 1 samples in 1 input files
Calling variants ...
Total processed positions in chr19 (chunk 9/12) : 1
Total time elapsed: 0.01 s
[INFO] Delay 3 seconds before starting variant calling ...
[mpileup] 1 samples in 1 input files
Calling variants ...
Total processed positions in chr19 (chunk 11/12) : 0
Total time elapsed: 0.00 s
[INFO] No vcf output for file /tmp/hg00150_apoe/clair3_output/tmp/pileup_output/
    pileup_chr19_11.vcf, remove empty file

[INFO] 2/7 Select heterozygous SNP variants for Whatshap phasing and haplotagging
[93m[WARNING] Cannot find any 0/1 variant in pileup output using variant quality cut-
    off proportion: 0.30000000000000004, total heterozygous variants: 0[0m
[93m[WARNING] Set low variant quality score cut-off to 0.0[0m
[INFO] Select heterozygous pileup variants exceeding phasing quality cutoff 0
[INFO] Total heterozygous SNP positions selected: chr19: 2

[INFO] 3/7 Phase VCF file using Whatshap
This is WhatsHap 2.8 running under Python 3.11.14
Working on 1 sample from 1 family

# Working on contig chr19 in individual HG00150
Found 2 usable heterozygous variants (0 skipped due to missing genotypes)
Number of supplementary alignments: 0
Number of non-singleton groups: 14
Skipped 0 groups
Found 112 reads covering 2 variants
Kept 17 reads that cover at least two variants each
Selected 15 most phase-informative reads covering 2 variants
Phasing 1 sample by solving the MEC problem ...
Largest block contains 2 variants (100.0% of accessible variants) between position
    44908684 and 44908822

# Resource usage
Maximum memory usage: 0.188 GB
Time spent reading BAM/CRAM:                    0.3 s
Time spent parsing VCF:                         0.0 s
Time spent selecting reads:                     0.0 s
Time spent phasing:                             0.0 s
Time spent writing VCF:                         0.0 s
Time spent finding components:                  0.0 s
Time spent on rest:                             0.1 s
Total elapsed time:                             0.5 s

[INFO] 4/7 Haplotag input BAM file using Whatshap
Found 1 sample(s) in input VCF
Found 1 sample(s) in BAM file
```

```
Number of supplementary alignments: 0
Number of non-singleton groups: 16
Skipped 0 groups
Found 124 reads covering 2 variants

== SUMMARY ==
Total alignments processed:                        3453
Alignments that could be tagged:                    248
Alignments spanning multiple phase sets:              0
Finished in 0.5 s

[INFO] 5/7 Select candidates for full-alignment calling
[93m[WARNING] Cannot find any low-quality 0/1 or 1/1 variant in pileup output using
    variant quality cut-off proportion: 0.3, total variants: 2[0m
[93m[WARNING] Set low variant quality score cut-off to 0.0[0m
[93m[WARNING] Cannot find any low-quality 0/0 reference calls in pileup output using
    reference quality cut-off proportion: 0.3, total reference calls: 0[0m
[93m[WARNING] Set low reference quality score cut-off to 0.0[0m
[INFO] Set variants quality cutoff 0.0
[INFO] Set reference calls quality cutoff 0.0
[93m[WARNING] Cannot find any low-quality 0/0, 0/1 or 1/1 variant in pileup output in
    contig chr19[0m
[93m[WARNING] Copying pileup.vcf.gz to /tmp/hg00150_apoe/clair3_output/merge_output.
    vcf.gz[0m
[INFO] Low quality reference calls to be processed in chr19: 0
[INFO] Low quality variants to be processed in chr19: 0

[INFO] 6/7 Call low-quality variants using full-alignment model
[INFO] Delay 0 seconds before starting variant calling ...
[93m[WARNING] Skip full-alignment variant calling for empty full-alignment regions[0m
[93m[WARNING] No vcf file found with prefix:/tmp/hg00150_apoe/clair3_output/tmp/
    full_alignment_output/full_alignment, output empty vcf file[0m
[INFO] Exit in full-alignment variant calling
```

**Turn 6 ▪ reasoning**

Clair3 completed successfully. Now let me look at the output VCF to see the genotypes at the two SNP positions.

**Turn 6 ▪ tool call: run_shell**

```
run_shell:
bcftools view -H /tmp/hg00150_apoe/clair3_output/merge_output.vcf.gz 2>&1
```

**Result ▪ run_shell**

```
returncode: 0
stdout:
chr19    44908684    .    T    C    24.68    PASS    P    GT:GQ:DP:AD:AF
    0/1:24:67:37,30:0.4478
chr19    44908822    .    C    T    21.57    PASS    P    GT:GQ:DP:AD:AF
    0/1:21:77:33,44:0.5714
```

**Turn 7 ▪ reasoning**

Let me now interpret the genotypes:
rs429358 (chr19:44908684): ref T, alt C → genotype 0/1 (heterozygous T/C)

rs7412 (chr19:44908822): ref C, alt T → genotype 0/1 (heterozygous C/T)
Now let's determine haplotypes:
The SNPs:
- rs429358 at chr19:44908684 (ref T, alt C)
- rs7412 at chr19:44908822 (ref C, alt T)
The three common haplotypes:
- $\varepsilon4$ = (rs429358 C, rs7412 C)
- $\varepsilon3$ = (T, C)
- $\varepsilon2$ = (T, T)
The individual is heterozygous at both SNPs. Let me think about the possible diplotypes:
At rs429358: T/C → one allele has T, one has C
At rs7412: C/T → one allele has C, one has T
Now from the Whatshap phasing output, I saw it found 2 variants in one block and phased them. But I don't see the phase set information in the VCF output. Let me check if the phased VCF has PS tags.
Wait, the merge_output.vcf.gz might not have phasing info. Let me check the phased output. But actually, given the haplotype definitions, we can figure out:
Possible haplotypes given the genotypes:
- rs429358: one T, one C
- rs7412: one C, one T
Possible combinations for two haplotypes:
- Haplotype 1: (T at rs429358, C at rs7412) = $\varepsilon3$
- Haplotype 2: (C at rs429358, T at rs7412) = this is (C, T) — not one of the three common haplotypes!
Wait, let me re-read:
- $\varepsilon4$ = (rs429358 C, rs7412 C)
- $\varepsilon3$ = (T, C)
- $\varepsilon2$ = (T, T)
So (C, T) at these positions isn't one of the common haplotypes. But wait, could it be that they're phased differently?
Actually, let me check - the genotypes are both 0/1, so:
- At rs429358: one chromosome has T, one has C
- At rs7412: one chromosome has C, one has T
If the phasing from Whatshap tells us which combination goes together, we could determine the diplotypes. Let me check the phased VCF.
But actually, let me look at this more carefully. The two possible phased solutions are:
Solution A (if T at rs429358 goes with C at rs7412 on one chromosome):
- Chromosome 1: (T, C) = $\varepsilon3$
- Chromosome 2: (C, T) = this is not a standard $\varepsilon$ allele
Solution B (if T at rs429358 goes with T at rs7412 on one chromosome):
- Chromosome 1: (T, T) = $\varepsilon2$
- Chromosome 2: (C, C) = $\varepsilon4$
So if the phasing is T-T / C-C, the diplotype is $\varepsilon2/\varepsilon4$. If it's T-C / C-T, it would be $\varepsilon3$ / something unusual.
Let me check the phased VCF to see which alleles are on the same chromosome.

**Turn 7 ▪ tool call: run_shell**

```
run_shell:
zcat /tmp/hg00150_apoe/clair3_output/merge_output.vcf.gz 2>&1 | grep -v '^#'
```

**Result ▪ run_shell**

```
returncode: 0
stdout:
chr19    44908684    .    T    C    24.68    PASS    P    GT:GQ:DP:AD:AF
   0/1:24:67:37,30:0.4478
```

```
chr19    44908822    .    C    T    21.57    PASS    P    GT:GQ:DP:AD:AF
    0/1:21:77:33,44:0.5714
```

**Turn 8 ▪ reasoning**

The GT doesn't show phasing (no | separator). Let me check the phased VCF from Whatshap which would be at a different path. Let me look in the tmp directory for the phased VCF.

**Turn 8 ▪ tool call: run_shell**

```
run_shell:
find /tmp/hg00150_apoe/clair3_output -name "*.vcf*" -type f 2>&1
```

**Result ▪ run_shell**

```
returncode: 0
stdout:
/tmp/hg00150_apoe/clair3_output/full_alignment.vcf.gz
/tmp/hg00150_apoe/clair3_output/full_alignment.vcf.gz.tbi
/tmp/hg00150_apoe/clair3_output/pileup.vcf.gz.tbi
/tmp/hg00150_apoe/clair3_output/tmp/phase_output/phase_vcf/chr19.vcf
/tmp/hg00150_apoe/clair3_output/tmp/phase_output/phase_vcf/phased_chr19.vcf.gz
/tmp/hg00150_apoe/clair3_output/tmp/phase_output/phase_vcf/phased_chr19.vcf.gz.tbi
/tmp/hg00150_apoe/clair3_output/tmp/pileup_output/pileup_chr19_8.vcf
/tmp/hg00150_apoe/clair3_output/tmp/pileup_output/pileup_chr19_4.vcf
/tmp/hg00150_apoe/clair3_output/tmp/pileup_output/pileup_chr19_9.vcf
/tmp/hg00150_apoe/clair3_output/tmp/pileup_output/pileup_chr19_5.vcf
/tmp/hg00150_apoe/clair3_output/pileup.vcf.gz
/tmp/hg00150_apoe/clair3_output/merge_output.vcf.gz
```

**Turn 9 ▪ reasoning**

Let me look at the phased VCF from Whatshap.

**Turn 9 ▪ tool call: run_shell**

```
run_shell:
zcat /tmp/hg00150_apoe/clair3_output/tmp/phase_output/phase_vcf/phased_chr19.vcf.gz
    2>&1
```

**Result ▪ run_shell**

```
returncode: 0
stdout:
h=6485>
##contig=<ID=HLA-DQA1*01:02:01:03,length=6485>
##contig=<ID=HLA-DQA1*01:02:01:04,length=6492>
##contig=<ID=HLA-DQA1*01:03:01:01,length=6485>
##contig=<ID=HLA-DQA1*01:03:01:02,length=6492>
##contig=<ID=HLA-DQA1*01:04:01:01,length=6484>
##contig=<ID=HLA-DQA1*01:04:01:02,length=6485>
##contig=<ID=HLA-DQA1*01:05:01,length=6485>
##contig=<ID=HLA-DQA1*01:07,length=5959>
##contig=<ID=HLA-DQA1*01:10,length=5790>
##contig=<ID=HLA-DQA1*01:11,length=5926>
##contig=<ID=HLA-DQA1*02:01,length=6403>
##contig=<ID=HLA-DQA1*03:01:01,length=6437>
```

```
##contig=<ID=HLA-DQA1*03:02,length=6437>
##contig=<ID=HLA-DQA1*03:03:01,length=6437>
##contig=<ID=HLA-DQA1*04:01:02:01,length=5853>
##contig=<ID=HLA-DQA1*04:01:02:02,length=5666>
##contig=<ID=HLA-DQA1*04:02,length=6210>
##contig=<ID=HLA-DQA1*05:01:01:01,length=5806>
##contig=<ID=HLA-DQA1*05:01:01:02,length=6529>
##contig=<ID=HLA-DQA1*05:03,length=6121>
##contig=<ID=HLA-DQA1*05:05:01:01,length=6593>
##contig=<ID=HLA-DQA1*05:05:01:02,length=6597>
##contig=<ID=HLA-DQA1*05:05:01:03,length=6393>
##contig=<ID=HLA-DQA1*05:11,length=6589>
##contig=<ID=HLA-DQA1*06:01:01,length=5878>
##contig=<ID=HLA-DQB1*02:01:01,length=7480>
##contig=<ID=HLA-DQB1*02:02:01,length=7471>
##contig=<ID=HLA-DQB1*03:01:01:01,length=7231>
##contig=<ID=HLA-DQB1*03:01:01:02,length=7230>
##contig=<ID=HLA-DQB1*03:01:01:03,length=7231>
##contig=<ID=HLA-DQB1*03:02:01,length=7126>
##contig=<ID=HLA-DQB1*03:03:02:01,length=7126>
##contig=<ID=HLA-DQB1*03:03:02:02,length=7126>
##contig=<ID=HLA-DQB1*03:03:02:03,length=6800>
##contig=<ID=HLA-DQB1*03:05:01,length=6934>
##contig=<ID=HLA-DQB1*05:01:01:01,length=7090>
##contig=<ID=HLA-DQB1*05:01:01:02,length=7090>
##contig=<ID=HLA-DQB1*05:03:01:01,length=7089>
##contig=<ID=HLA-DQB1*05:03:01:02,length=7089>
##contig=<ID=HLA-DQB1*06:01:01,length=7111>
##contig=<ID=HLA-DQB1*06:02:01,length=7102>
##contig=<ID=HLA-DQB1*06:03:01,length=7103>
##contig=<ID=HLA-DQB1*06:09:01,length=7102>
##contig=<ID=HLA-DRB1*01:01:01,length=10741>
##contig=<ID=HLA-DRB1*01:02:01,length=11229>
##contig=<ID=HLA-DRB1*03:01:01:01,length=13908>
##contig=<ID=HLA-DRB1*03:01:01:02,length=13426>
##contig=<ID=HLA-DRB1*04:03:01,length=15246>
##contig=<ID=HLA-DRB1*07:01:01:01,length=16110>
##contig=<ID=HLA-DRB1*07:01:01:02,length=16120>
##contig=<ID=HLA-DRB1*08:03:02,length=13562>
##contig=<ID=HLA-DRB1*09:21,length=16039>
##contig=<ID=HLA-DRB1*10:01:01,length=13501>
##contig=<ID=HLA-DRB1*11:01:01,length=13921>
##contig=<ID=HLA-DRB1*11:01:02,length=13931>
##contig=<ID=HLA-DRB1*11:04:01,length=13919>
##contig=<ID=HLA-DRB1*12:01:01,length=13404>
##contig=<ID=HLA-DRB1*12:17,length=11260>
##contig=<ID=HLA-DRB1*13:01:01,length=13935>
##contig=<ID=HLA-DRB1*13:02:01,length=13941>
##contig=<ID=HLA-DRB1*14:05:01,length=13933>
##contig=<ID=HLA-DRB1*14:54:01,length=13936>
##contig=<ID=HLA-DRB1*15:01:01:01,length=11080>
##contig=<ID=HLA-DRB1*15:01:01:02,length=11571>
##contig=<ID=HLA-DRB1*15:01:01:03,length=11056>
##contig=<ID=HLA-DRB1*15:01:01:04,length=11056>
##contig=<ID=HLA-DRB1*15:02:01,length=10313>
##contig=<ID=HLA-DRB1*15:03:01:01,length=11567>
##contig=<ID=HLA-DRB1*15:03:01:02,length=11569>
##contig=<ID=HLA-DRB1*16:02:01,length=11005>
```

```
##FORMAT=<ID=AD,Number=.,Type=Integer,Description="Observed allele depths">
##commandline="(whatshap 2.8) phase --output /tmp/hg00150_apoe/clair3_output/tmp/
    phase_output/phase_vcf/phased_chr19.vcf.gz --reference /autofs/bal19/zxzheng/
    testData/ilmn/data/GRCh38_full_analysis_set_plus_decoy_hla.fa --chromosome chr19
    --distrust-genotypes --ignore-read-groups /tmp/hg00150_apoe/clair3_output/tmp/
    phase_output/phase_vcf/chr19.vcf /autofs/nas5/xyu2/projects/clair3-api/clair3r/
    paper_experiments/.cache/apoe/HG00150_apoe.bam"
##FORMAT=<ID=PS,Number=1,Type=Integer,Description="Phase set identifier">
#CHROM    POS    ID    REF    ALT    QUAL    FILTER    INFO    FORMAT    HG00150
chr19    44908684    .    T    C    24.68    PASS    P    GT:GQ:DP:AD:AF:PS
    1|0:24:67:37,30:0.4478:44908684
chr19    44908822    .    C    T    21.57    PASS    P    GT:GQ:DP:AD:AF:PS
    0|1:21:77:33,44:0.5714:44908684
```

**Turn 10 ▪ reasoning**

Now I can see the phased genotypes:
rs429358 (chr19:44908684): GT = 1|0 → Allele 1 (haplotype 1) has the ALT (C), Allele 0 (haplotype 2) has the REF (T)
rs7412 (chr19:44908822): GT = 0|1 → Allele 1 (haplotype 1) has the REF (C), Allele 0 (haplotype 2) has the ALT (T)
So:
- Haplotype 1 (PS phase set, allele 1): rs429358=C, rs7412=C → this is $\varepsilon 4$
- Haplotype 2 (PS phase set, allele 0): rs429358=T, rs7412=T → this is $\varepsilon 2$
The diplotype is $\varepsilon 2/\varepsilon 4$.
Wait, let me double-check. The GT format is:
At rs429358: 1|0 means:
- first haplotype (1) = alt allele = C
- second haplotype (0) = ref allele = T
At rs7412: 0|1 means:
- first haplotype (0) = ref allele = C
- second haplotype (1) = alt allele = T
So:
- First haplotype: (rs429358=C, rs7412=C) = $\varepsilon 4$
- Second haplotype: (rs429358=T, rs7412=T) = $\varepsilon 2$
Diplotype: $\varepsilon 2/\varepsilon 4$
Convention is usually to list them in order ($\varepsilon 2$ before $\varepsilon 4$), so $\varepsilon 2/\varepsilon 4$.
ANSWER: e2/e4

**Final answer**

The phased VCF reveals the two haplotypes clearly:
- **Haplotype 1**: rs429358=C, rs7412=C → **$\varepsilon 4$**
- **Haplotype 2**: rs429358=T, rs7412=T → **$\varepsilon 2$**
ANSWER: e2/e4

### Tools

#### Tools ▪ HG00136

3 turns, 2 tool calls, 11,443 tokens, 20.1 s; answer: e4/e4

*System prompt: §S1.1 (Tools); task prompt: §S1.2 (sample HG00136).*

**Turn 1 ▪ reasoning**

Let me break this down step by step:
1. First, I need to determine the sequencing platform for sample HG00136. I'll use 'bam_header' to get the BAM header and figure out the platform from the @RG PL tag or other hints.
2. Then I need to call variants at the two SNP positions: rs429358 (chr19:44908684) and rs7412 (chr19:44908822).
3. Finally, combine the genotypes to determine the APOE diplotype.
Let me start by getting the BAM header and also the gene region for APOE (though I already have the positions, but it might be useful).
Actually, I'll start with the BAM header to determine the platform.

**Turn 1 ▪ tool call: bam_header**

```
bam_header({"sample": "HG00136"})
```

**Result ▪ bam_header**

```
{
  "sample": "HG00136",
  "header": [
    "@HD\tVN:1.5\tGO:none\tSO:coordinate",
    "@PG\tID:MarkDuplicates\tVN:2.4.1(7
   c4d36e011df1aec4689b51efcada44e92d1817f_1464389670)\tCL:picard.sam.markduplicates.
   MarkDuplicates INPUT=[/data/analysis/DarnellR/Project_CCDG_13607_B01_GRM_WGS/
   Sample_HG00136/analysis/HG0013",
    "@PG\tID:bwa\tPN:bwa\tVN:0.7.15-r1140\tCL:/data/NYGC/Software/bwa/bwa-0.7.15/bwa
   mem -Y -K 100000000 -t 16 -R @RG\\tID:HG00136_CCAAGTCT-AAGGATGA_HFM5KDSXX_L001\\
   tPL:illumina\\tPM:Unknown\\tLB:HG00136\\tDS:GRCh38\\tSM:HG00136\\tCN:NY",
    "@PG\tID:bwa.1\tPN:bwa\tVN:0.7.15-r1140\tCL:/data/NYGC/Software/bwa/bwa-0.7.15/bwa
    mem -Y -K 100000000 -t 16 -R @RG\\tID:HG00136_CCAAGTCT-AAGGATGA_HFJFFDSXX_L001\\
   tPL:illumina\\tPM:Unknown\\tLB:HG00136\\tDS:GRCh38\\tSM:HG00136\\tCN:",
    "@PG\tID:bwa.2\tPN:bwa\tVN:0.7.15-r1140\tCL:/data/NYGC/Software/bwa/bwa-0.7.15/bwa
    mem -Y -K 100000000 -t 16 -R @RG\\tID:HG00136_CCAAGTCT-AAGGATGA_HFKLNDSXX_L002\\
   tPL:illumina\\tPM:Unknown\\tLB:HG00136\\tDS:GRCh38\\tSM:HG00136\\tCN:",
    "@PG\tID:bwa.3\tPN:bwa\tVN:0.7.15-r1140\tCL:/data/NYGC/Software/bwa/bwa-0.7.15/bwa
    mem -Y -K 100000000 -t 16 -R @RG\\tID:HG00136_CCAAGTCT-AAGGATGA_HFJFFDSXX_L004\\
   tPL:illumina\\tPM:Unknown\\tLB:HG00136\\tDS:GRCh38\\tSM:HG00136\\tCN:",
    "@PG\tID:bwa.4\tPN:bwa\tVN:0.7.15-r1140\tCL:/data/NYGC/Software/bwa/bwa-0.7.15/bwa
    mem -Y -K 100000000 -t 16 -R @RG\\tID:HG00136_CCAAGTCT-AAGGATGA_HFKLNDSXX_L004\\
   tPL:illumina\\tPM:Unknown\\tLB:HG00136\\tDS:GRCh38\\tSM:HG00136\\tCN:",
    "@PG\tID:bwa.5\tPN:bwa\tVN:0.7.15-r1140\tCL:/data/NYGC/Software/bwa/bwa-0.7.15/bwa
    mem -Y -K 100000000 -t 16 -R @RG\\tID:HG00136_CCAAGTCT-AAGGATGA_HFM5KDSXX_L003\\
   tPL:illumina\\tPM:Unknown\\tLB:HG00136\\tDS:GRCh38\\tSM:HG00136\\tCN:",
    "@PG\tID:bwa.6\tPN:bwa\tVN:0.7.15-r1140\tCL:/data/NYGC/Software/bwa/bwa-0.7.15/bwa
    mem -Y -K 100000000 -t 16 -R @RG\\tID:HG00136_CCAAGTCT-AAGGATGA_HFKLNDSXX_L001\\
   tPL:illumina\\tPM:Unknown\\tLB:HG00136\\tDS:GRCh38\\tSM:HG00136\\tCN:",
    "@PG\tID:bwa.7\tPN:bwa\tVN:0.7.15-r1140\tCL:/data/NYGC/Software/bwa/bwa-0.7.15/bwa
    mem -Y -K 100000000 -t 16 -R @RG\\tID:HG00136_CCAAGTCT-AAGGATGA_HFJFFDSXX_L002\\
   tPL:illumina\\tPM:Unknown\\tLB:HG00136\\tDS:GRCh38\\tSM:HG00136\\tCN:",
    "@PG\tID:bwa.8\tPN:bwa\tVN:0.7.15-r1140\tCL:/data/NYGC/Software/bwa/bwa-0.7.15/bwa
    mem -Y -K 100000000 -t 16 -R @RG\\tID:HG00136_CCAAGTCT-AAGGATGA_HFKLNDSXX_L003\\
   tPL:illumina\\tPM:Unknown\\tLB:HG00136\\tDS:GRCh38\\tSM:HG00136\\tCN:",
    "@PG\tID:bwa.9\tPN:bwa\tVN:0.7.15-r1140\tCL:/data/NYGC/Software/bwa/bwa-0.7.15/bwa
    mem -Y -K 100000000 -t 16 -R @RG\\tID:HG00136_CCAAGTCT-AAGGATGA_HFM5KDSXX_L004\\
   tPL:illumina\\tPM:Unknown\\tLB:HG00136\\tDS:GRCh38\\tSM:HG00136\\tCN:",
    "@PG\tID:bwa.A\tPN:bwa\tVN:0.7.15-r1140\tCL:/data/NYGC/Software/bwa/bwa-0.7.15/bwa
    mem -Y -K 100000000 -t 16 -R @RG\\tID:HG00136_CCAAGTCT-AAGGATGA_HFJFFDSXX_L003\\
   tPL:illumina\\tPM:Unknown\\tLB:HG00136\\tDS:GRCh38\\tSM:HG00136\\tCN:",
```

```
    "@PG\tID:bwa.B\tPN:bwa\tVN:0.7.15-r1140\tCL:/data/NYGC/Software/bwa/bwa-0.7.15/bwa
     mem -Y -K 100000000 -t 16 -R @RG\\tID:HG00136_CCAAGTCT-AAGGATGA_HFM5KDSXX_L002\\
    tPL:illumina\\tPM:Unknown\\tLB:HG00136\\tDS:GRCh38\\tSM:HG00136\\tCN:",
    "@PG\tID:GATK PrintReads\tVN:3.5-0-g36282e4\tCL:readGroup=null platform=null
    number=-1 sample_file=[] sample_name=[] simplify=false no_pg_tag=false",
    "@PG\tID:samtools\tPN:samtools\tPP:GATK PrintReads\tVN:1.23\tCL:samtools view -T /
    autofs/bal19/zxzheng/testData/ilmn/data/GRCh38_full_analysis_set_plus_decoy_hla.fa
     -b -o paper_experiments/.cache/apoe/HG00136_apoe.bam https://",
    "@PG\tID:samtools.1\tPN:samtools\tPP:samtools\tVN:1.23\tCL:/autofs/nas5/xyu2/
    softwares/miniforge3/envs/clair3-torch/bin/samtools view -H /autofs/nas5/xyu2/
    projects/clair3-api/clair3r/paper_experiments/.cache/apoe/HG00136_apoe.",
    "@RG\tID:HG00136_CCAAGTCT-AAGGATGA_HFJFFDSXX_L001\tPL:illumina\tPM:Unknown\tLB:
    HG00136\tDS:GRCh38\tSM:HG00136\tCN:NYGenome\tPU:HFJFFDSXX.1.CCAAGTCT",
    "@RG\tID:HG00136_CCAAGTCT-AAGGATGA_HFJFFDSXX_L002\tPL:illumina\tPM:Unknown\tLB:
    HG00136\tDS:GRCh38\tSM:HG00136\tCN:NYGenome\tPU:HFJFFDSXX.2.CCAAGTCT",
    "@RG\tID:HG00136_CCAAGTCT-AAGGATGA_HFJFFDSXX_L003\tPL:illumina\tPM:Unknown\tLB:
    HG00136\tDS:GRCh38\tSM:HG00136\tCN:NYGenome\tPU:HFJFFDSXX.3.CCAAGTCT",
    "@RG\tID:HG00136_CCAAGTCT-AAGGATGA_HFJFFDSXX_L004\tPL:illumina\tPM:Unknown\tLB:
    HG00136\tDS:GRCh38\tSM:HG00136\tCN:NYGenome\tPU:HFJFFDSXX.4.CCAAGTCT",
    "@RG\tID:HG00136_CCAAGTCT-AAGGATGA_HFKLNDSXX_L001\tPL:illumina\tPM:Unknown\tLB:
    HG00136\tDS:GRCh38\tSM:HG00136\tCN:NYGenome\tPU:HFKLNDSXX.1.CCAAGTCT",
    "@RG\tID:HG00136_CCAAGTCT-AAGGATGA_HFKLNDSXX_L002\tPL:illumina\tPM:Unknown\tLB:
    HG00136\tDS:GRCh38\tSM:HG00136\tCN:NYGenome\tPU:HFKLNDSXX.2.CCAAGTCT",
    "@RG\tID:HG00136_CCAAGTCT-AAGGATGA_HFKLNDSXX_L003\tPL:illumina\tPM:Unknown\tLB:
    HG00136\tDS:GRCh38\tSM:HG00136\tCN:NYGenome\tPU:HFKLNDSXX.3.CCAAGTCT",
    "@RG\tID:HG00136_CCAAGTCT-AAGGATGA_HFKLNDSXX_L004\tPL:illumina\tPM:Unknown\tLB:
    HG00136\tDS:GRCh38\tSM:HG00136\tCN:NYGenome\tPU:HFKLNDSXX.4.CCAAGTCT",
    "@RG\tID:HG00136_CCAAGTCT-AAGGATGA_HFM5KDSXX_L001\tPL:illumina\tPM:Unknown\tLB:
    HG00136\tDS:GRCh38\tSM:HG00136\tCN:NYGenome\tPU:HFM5KDSXX.1.CCAAGTCT",
    "@RG\tID:HG00136_CCAAGTCT-AAGGATGA_HFM5KDSXX_L002\tPL:illumina\tPM:Unknown\tLB:
    HG00136\tDS:GRCh38\tSM:HG00136\tCN:NYGenome\tPU:HFM5KDSXX.2.CCAAGTCT",
    "@RG\tID:HG00136_CCAAGTCT-AAGGATGA_HFM5KDSXX_L003\tPL:illumina\tPM:Unknown\tLB:
    HG00136\tDS:GRCh38\tSM:HG00136\tCN:NYGenome\tPU:HFM5KDSXX.3.CCAAGTCT",
    "@RG\tID:HG00136_CCAAGTCT-AAGGATGA_HFM5KDSXX_L004\tPL:illumina\tPM:Unknown\tLB:
    HG00136\tDS:GRCh38\tSM:HG00136\tCN:NYGenome\tPU:HFM5KDSXX.4.CCAAGTCT"
  ],
  "n_reference_contigs": 3366,
  "note": "infer the sequencing platform from these records, then select the matching
    model (see discover_models)"
}
```

**Turn 2 ▪ reasoning**

The platform is clearly Illumina (PL:illumina in all @RG records). Now I need to get the genotypes at the two SNP positions. Let me use ‘genotypes_at‘ to call variants and read the genotypes at both positions.
The region should cover both positions: chr19:44908684-44908822. Let me use a region that covers both, like chr19:44908684-44908822.
Positions:
- chr19:44908684 (rs429358)
- chr19:44908822 (rs7412)
Platform: ilmn

**Turn 2 ▪ tool call: genotypes_at**

```
genotypes_at({"sample": "HG00136", "platform": "ilmn", "region": "chr19
    :44908684-44908822", "positions": ["chr19:44908684", "chr19:44908822"]})
```

**Result ▪ genotypes_at**

```
{
  "sample": "HG00136",
  "platform": "ilmn",
  "region": "chr19:44908684-44908822",
  "vcf_path": "/autofs/nas5/xyu2/projects/clair3-api/clair3r/paper_experiments/.cache/
    e4_agent_work/HG00136_ilmn_chr19_44908684-44908822/merge_output.vcf",
  "genotypes": [
    {
      "position": "chr19:44908684",
      "ref": "T",
      "alt": "C",
      "gt": "1/1",
      "called": true
    },
    {
      "position": "chr19:44908822",
      "alt": ".",
      "gt": "0/0",
      "called": false,
      "note": "no variant called at this position - homozygous reference"
    }
  ]
}
```

**Turn 3 ▪ reasoning**

Let me interpret the results:
- rs429358 (chr19:44908684): genotype is 1/1, which means homozygous for the alt allele C. So both chromosomes have C at this position.
- rs7412 (chr19:44908822): genotype is 0/0, which means homozygous reference. The ref is C, so both chromosomes have C at this position.
Now let me map these to haplotypes:
- $\varepsilon 4$ haplotype: (rs429358 = C, rs7412 = C)
- $\varepsilon 3$ haplotype: (rs429358 = T, rs7412 = C)
- $\varepsilon 2$ haplotype: (rs429358 = T, rs7412 = T)
At rs429358: genotype 1/1 = C/C. Both alleles are C.
At rs7412: genotype 0/0 = C/C. Both alleles are C.
So each chromosome has: (rs429358=C, rs7412=C) $\rightarrow$ $\varepsilon 4/\varepsilon 4$.
The diplotype is $\varepsilon 4/\varepsilon 4$.

**Final answer**

Now let me interpret the genotypes:
- **rs429358** (chr19:44908684): GT = ‘1/1‘ → homozygous **C/C**
- **rs7412** (chr19:44908822): GT = ‘0/0‘ → homozygous **C/C** (reference)
Each chromosome carries (rs429358=C, rs7412=C), which is the **$\varepsilon 4$** haplotype. Both chromosomes are $\varepsilon 4$, so the diplotype is $\varepsilon 4/\varepsilon 4$.
ANSWER: e4/e4

#### Tools ▪ HG00150

3 turns, 3 tool calls, 12,151 tokens, 27.7 s; answer: e2/e4

*System prompt: §S1.1 (Tools); task prompt: §S1.2 (sample HG00150).*

**Turn 1 ▪ reasoning**

The user wants me to determine the APOE diplotype for sample HG00150. Let me follow the steps:
1. First, determine the sequencing platform by looking at the BAM header.
2. Then call variants at the two SNP positions.
3. Combine genotypes to determine the diplotype.
Let me start by getting the BAM header and discovering available models.

**Turn 1 ▪ tool call: bam_header**

```
bam_header({"sample": "HG00150"})
```

**Turn 1 ▪ tool call: discover_models**

```
discover_models({})
```

**Result ▪ bam_header**

```
{
  "sample": "HG00150",
  "header": [
    "@HD\tVN:1.5\tGO:none\tSO:coordinate",
    "@PG\tID:MarkDuplicates\tVN:2.4.1(7
    c4d36e011df1aec4689b51efcada44e92d1817f_1464389670)\tCL:picard.sam.markduplicates.
    MarkDuplicates INPUT=[/data/analysis/DarnellR/Project_CCDG_13607_B01_GRM_WGS/
    Sample_HG00150/analysis/HG0015",
    "@PG\tID:bwa\tPN:bwa\tVN:0.7.15-r1140\tCL:/data/NYGC/Software/bwa/bwa-0.7.15/bwa
    mem -Y -K 100000000 -t 16 -R @RG\\tID:HG00150_CATAGAGT-TGCCACCA_HFH25DSXX_L004\\
    tPL:illumina\\tPM:Unknown\\tLB:HG00150\\tDS:GRCh38\\tSM:HG00150\\tCN:NY",
    "@PG\tID:bwa.1\tPN:bwa\tVN:0.7.15-r1140\tCL:/data/NYGC/Software/bwa/bwa-0.7.15/bwa
     mem -Y -K 100000000 -t 16 -R @RG\\tID:HG00150_CATAGAGT-TGCCACCA_HFGYNDSXX_L004\\
    tPL:illumina\\tPM:Unknown\\tLB:HG00150\\tDS:GRCh38\\tSM:HG00150\\tCN:",
    "@PG\tID:bwa.2\tPN:bwa\tVN:0.7.15-r1140\tCL:/data/NYGC/Software/bwa/bwa-0.7.15/bwa
     mem -Y -K 100000000 -t 16 -R @RG\\tID:HG00150_CATAGAGT-TGCCACCA_HFH25DSXX_L001\\
    tPL:illumina\\tPM:Unknown\\tLB:HG00150\\tDS:GRCh38\\tSM:HG00150\\tCN:",
    "@PG\tID:bwa.3\tPN:bwa\tVN:0.7.15-r1140\tCL:/data/NYGC/Software/bwa/bwa-0.7.15/bwa
     mem -Y -K 100000000 -t 16 -R @RG\\tID:HG00150_CATAGAGT-TGCCACCA_HFGYNDSXX_L001\\
    tPL:illumina\\tPM:Unknown\\tLB:HG00150\\tDS:GRCh38\\tSM:HG00150\\tCN:",
    "@PG\tID:bwa.4\tPN:bwa\tVN:0.7.15-r1140\tCL:/data/NYGC/Software/bwa/bwa-0.7.15/bwa
     mem -Y -K 100000000 -t 16 -R @RG\\tID:HG00150_CATAGAGT-TGCCACCA_HFGYNDSXX_L003\\
    tPL:illumina\\tPM:Unknown\\tLB:HG00150\\tDS:GRCh38\\tSM:HG00150\\tCN:",
    "@PG\tID:bwa.5\tPN:bwa\tVN:0.7.15-r1140\tCL:/data/NYGC/Software/bwa/bwa-0.7.15/bwa
     mem -Y -K 100000000 -t 16 -R @RG\\tID:HG00150_CATAGAGT-TGCCACCA_HFKWWDSXX_L003\\
    tPL:illumina\\tPM:Unknown\\tLB:HG00150\\tDS:GRCh38\\tSM:HG00150\\tCN:",
    "@PG\tID:bwa.6\tPN:bwa\tVN:0.7.15-r1140\tCL:/data/NYGC/Software/bwa/bwa-0.7.15/bwa
     mem -Y -K 100000000 -t 16 -R @RG\\tID:HG00150_CATAGAGT-TGCCACCA_HFKWWDSXX_L001\\
    tPL:illumina\\tPM:Unknown\\tLB:HG00150\\tDS:GRCh38\\tSM:HG00150\\tCN:",
    "@PG\tID:bwa.7\tPN:bwa\tVN:0.7.15-r1140\tCL:/data/NYGC/Software/bwa/bwa-0.7.15/bwa
     mem -Y -K 100000000 -t 16 -R @RG\\tID:HG00150_CATAGAGT-TGCCACCA_HFGYNDSXX_L002\\
    tPL:illumina\\tPM:Unknown\\tLB:HG00150\\tDS:GRCh38\\tSM:HG00150\\tCN:",
    "@PG\tID:bwa.8\tPN:bwa\tVN:0.7.15-r1140\tCL:/data/NYGC/Software/bwa/bwa-0.7.15/bwa
     mem -Y -K 100000000 -t 16 -R @RG\\tID:HG00150_CATAGAGT-TGCCACCA_HFKWWDSXX_L004\\
    tPL:illumina\\tPM:Unknown\\tLB:HG00150\\tDS:GRCh38\\tSM:HG00150\\tCN:",
    "@PG\tID:bwa.9\tPN:bwa\tVN:0.7.15-r1140\tCL:/data/NYGC/Software/bwa/bwa-0.7.15/bwa
     mem -Y -K 100000000 -t 16 -R @RG\\tID:HG00150_CATAGAGT-TGCCACCA_HFH25DSXX_L003\\
    tPL:illumina\\tPM:Unknown\\tLB:HG00150\\tDS:GRCh38\\tSM:HG00150\\tCN:",
    "@PG\tID:bwa.A\tPN:bwa\tVN:0.7.15-r1140\tCL:/data/NYGC/Software/bwa/bwa-0.7.15/bwa
     mem -Y -K 100000000 -t 16 -R @RG\\tID:HG00150_CATAGAGT-TGCCACCA_HFH25DSXX_L002\\
    tPL:illumina\\tPM:Unknown\\tLB:HG00150\\tDS:GRCh38\\tSM:HG00150\\tCN:",
```

```
    "@PG\tID:bwa.B\tPN:bwa\tVN:0.7.15-r1140\tCL:/data/NYGC/Software/bwa/bwa-0.7.15/bwa
     mem -Y -K 100000000 -t 16 -R @RG\\tID:HG00150_CATAGAGT-TGCCACCA_HFKWWDSXX_L002\\
    tPL:illumina\\tPM:Unknown\\tLB:HG00150\\tDS:GRCh38\\tSM:HG00150\\tCN:",
    "@PG\tID:GATK PrintReads\tVN:3.5-0-g36282e4\tCL:readGroup=null platform=null
    number=-1 sample_file=[] sample_name=[] simplify=false no_pg_tag=false",
    "@PG\tID:samtools\tPN:samtools\tPP:GATK PrintReads\tVN:1.23\tCL:/autofs/nas5/xyu2/
    softwares/miniforge3/envs/clair3-torch/bin/samtools view -T /autofs/bal19/zxzheng/
    testData/ilmn/data/GRCh38_full_analysis_set_plus_decoy_hla.fa ",
    "@PG\tID:samtools.1\tPN:samtools\tPP:samtools\tVN:1.23\tCL:/autofs/nas5/xyu2/
    softwares/miniforge3/envs/clair3-torch/bin/samtools view -H /autofs/nas5/xyu2/
    projects/clair3-api/clair3r/paper_experiments/.cache/apoe/HG00150_apoe.",
    "@RG\tID:HG00150_CATAGAGT-TGCCACCA_HFGYNDSXX_L001\tPL:illumina\tPM:Unknown\tLB:
    HG00150\tDS:GRCh38\tSM:HG00150\tCN:NYGenome\tPU:HFGYNDSXX.1.CATAGAGT",
    "@RG\tID:HG00150_CATAGAGT-TGCCACCA_HFGYNDSXX_L002\tPL:illumina\tPM:Unknown\tLB:
    HG00150\tDS:GRCh38\tSM:HG00150\tCN:NYGenome\tPU:HFGYNDSXX.2.CATAGAGT",
    "@RG\tID:HG00150_CATAGAGT-TGCCACCA_HFGYNDSXX_L003\tPL:illumina\tPM:Unknown\tLB:
    HG00150\tDS:GRCh38\tSM:HG00150\tCN:NYGenome\tPU:HFGYNDSXX.3.CATAGAGT",
    "@RG\tID:HG00150_CATAGAGT-TGCCACCA_HFGYNDSXX_L004\tPL:illumina\tPM:Unknown\tLB:
    HG00150\tDS:GRCh38\tSM:HG00150\tCN:NYGenome\tPU:HFGYNDSXX.4.CATAGAGT",
    "@RG\tID:HG00150_CATAGAGT-TGCCACCA_HFH25DSXX_L001\tPL:illumina\tPM:Unknown\tLB:
    HG00150\tDS:GRCh38\tSM:HG00150\tCN:NYGenome\tPU:HFH25DSXX.1.CATAGAGT",
    "@RG\tID:HG00150_CATAGAGT-TGCCACCA_HFH25DSXX_L002\tPL:illumina\tPM:Unknown\tLB:
    HG00150\tDS:GRCh38\tSM:HG00150\tCN:NYGenome\tPU:HFH25DSXX.2.CATAGAGT",
    "@RG\tID:HG00150_CATAGAGT-TGCCACCA_HFH25DSXX_L003\tPL:illumina\tPM:Unknown\tLB:
    HG00150\tDS:GRCh38\tSM:HG00150\tCN:NYGenome\tPU:HFH25DSXX.3.CATAGAGT",
    "@RG\tID:HG00150_CATAGAGT-TGCCACCA_HFH25DSXX_L004\tPL:illumina\tPM:Unknown\tLB:
    HG00150\tDS:GRCh38\tSM:HG00150\tCN:NYGenome\tPU:HFH25DSXX.4.CATAGAGT",
    "@RG\tID:HG00150_CATAGAGT-TGCCACCA_HFKWWDSXX_L001\tPL:illumina\tPM:Unknown\tLB:
    HG00150\tDS:GRCh38\tSM:HG00150\tCN:NYGenome\tPU:HFKWWDSXX.1.CATAGAGT",
    "@RG\tID:HG00150_CATAGAGT-TGCCACCA_HFKWWDSXX_L002\tPL:illumina\tPM:Unknown\tLB:
    HG00150\tDS:GRCh38\tSM:HG00150\tCN:NYGenome\tPU:HFKWWDSXX.2.CATAGAGT",
    "@RG\tID:HG00150_CATAGAGT-TGCCACCA_HFKWWDSXX_L003\tPL:illumina\tPM:Unknown\tLB:
    HG00150\tDS:GRCh38\tSM:HG00150\tCN:NYGenome\tPU:HFKWWDSXX.3.CATAGAGT",
    "@RG\tID:HG00150_CATAGAGT-TGCCACCA_HFKWWDSXX_L004\tPL:illumina\tPM:Unknown\tLB:
    HG00150\tDS:GRCh38\tSM:HG00150\tCN:NYGenome\tPU:HFKWWDSXX.4.CATAGAGT"
  ],
  "n_reference_contigs": 3366,
  "note": "infer the sequencing platform from these records, then select the matching
    model (see discover_models)"
}
```

**Result ▪ discover_models**

```
{
  "server_version": "0.0.1",
  "models": [
    {
      "model_id": "full_alignment_hifi_revio",
      "platform": "hifi",
      "stage": "full_alignment"
    },
    {
      "model_id": "full_alignment_ilmn",
      "platform": "ilmn",
      "stage": "full_alignment"
    },
    {
      "model_id": "full_alignment_ont_hac_v500",
```

```
      "platform": "ont",
      "stage": "full_alignment"
    },
    {
      "model_id": "pileup_hifi_revio",
      "platform": "hifi",
      "stage": "pileup"
    },
    {
      "model_id": "pileup_ilmn",
      "platform": "ilmn",
      "stage": "pileup"
    },
    {
      "model_id": "pileup_ont_hac_v500",
      "platform": "ont",
      "stage": "pileup"
    }
  ]
}
```

**Turn 2 ▪ reasoning**

The BAM header shows PL:illumina in all @RG lines. So the platform is "ilmn" (Illumina). Now I need to call genotypes at the two SNP positions:
- rs429358: chr19:44908684 (ref T, alt C)
- rs7412: chr19:44908822 (ref C, alt T)

I'll use the ‘genotypes_at‘ tool with the ilmn platform. The region should cover both positions.

**Turn 2 ▪ tool call: genotypes_at**

```
genotypes_at({"sample": "HG00150", "platform": "ilmn", "region": "chr19
    :44908684-44908822", "positions": ["chr19:44908684", "chr19:44908822"]})
```

**Result ▪ genotypes_at**

```
{
  "sample": "HG00150",
  "platform": "ilmn",
  "region": "chr19:44908684-44908822",
  "vcf_path": "/autofs/nas5/xyu2/projects/clair3-api/clair3r/paper_experiments/.cache/
    e4_agent_work/HG00150_ilmn_chr19_44908684-44908822/merge_output.vcf",
  "genotypes": [
    {
      "position": "chr19:44908684",
      "ref": "T",
      "alt": "C",
      "gt": "0/1",
      "called": true
    },
    {
      "position": "chr19:44908822",
      "ref": "C",
      "alt": "T",
      "gt": "0/1",
      "called": true
    }
  ]
```

```
}
```

**Turn 3 ▪ reasoning**

Let me interpret the genotypes:
- rs429358 (chr19:44908684): GT = 0/1, meaning one T allele (ref) and one C allele (alt). So this person has one ε4 allele (C) and one non-ε4 (T) at this position.
- rs7412 (chr19:44908822): GT = 0/1, meaning one C allele (ref) and one T allele (alt). So this person has one non-ε2 (C) and one ε2 (T) at this position.
Now let me figure out the haplotypes:
The haplotypes are:
- ε4: (rs429358 C, rs7412 C)
- ε3: (rs429358 T, rs7412 C)
- ε2: (rs429358 T, rs7412 T)
The person is heterozygous at both positions:
- rs429358: T/C
- rs7412: C/T
The possible combinations for the two haplotypes:
- Haplotype 1: (rs429358 T, rs7412 C) = ε3
- Haplotype 2: (rs429358 C, rs7412 T)
Wait, is (C, T) a known haplotype? Let me check. The three common haplotypes are:
- ε4 = (C, C)
- ε3 = (T, C)
- ε2 = (T, T)
So (C, T) is not one of the standard haplotypes. But the person is heterozygous at both sites. Let me think about the phase.
If both are heterozygous (0/1), the possible arrangements are:
- Arrangement 1: Chromosome A has (T, C) = ε3, Chromosome B has (C, T) — this is not a standard haplotype
- Arrangement 2: Chromosome A has (T, T) = ε2, Chromosome B has (C, C) = ε4
Since ε2/ε4 is a valid diplotype combination (ε2 on one chromosome, ε4 on the other), and it accounts for all alleles (T/T at one chromosome for both positions? No wait).
Let me be more careful:
- rs429358: T/C (het)
- rs7412: C/T (het)
If we have ε2/ε4:
- ε2: (T, T) → contributes T at rs429358, T at rs7412
- ε4: (C, C) → contributes C at rs429358, C at rs7412
- Combined: rs429358 = T/C (het ), rs7412 = T/C (het )
Yes, ε2/ε4 matches both heterozygous calls.
If we had ε3/(C,T):
- ε3: (T, C) → T at rs429358, C at rs7412
- (C, T): → C at rs429358, T at rs7412
- Combined: rs429358 = T/C (het ), rs7412 = C/T (het )
But (C, T) is not one of the three common haplotypes. The standard clinical interpretation would be ε2/ε4 based on the three common haplotypes.
So the diplotype is ε2/ε4, which I'll write as e2/e4.

**Final answer**

Now let me interpret the genotypes:
- **rs429358** (chr19:44908684): GT = 0/1 → one T (ref), one C (alt)
- **rs7412** (chr19:44908822): GT = 0/1 → one C (ref), one T (alt)
Double heterozygosity means the two haplotypes are:
| Haplotype | rs429358 | rs7412 |
|———–|———-|——–|
| ε2 | T | T |
| ε4 | C | C |

The $\varepsilon 2$ haplotype contributes (T, T) and $\varepsilon 4$ contributes (C, C), producing heterozygous calls at both sites. The alternative phasing ($\varepsilon 3$ + a rare C/T haplotype) is not among the three common clinical haplotypes, so the standard diplotype is $\varepsilon 2/\varepsilon 4$.
ANSWER: e2/e4

### S5 Data and software resources

This note lists the public download paths for the sequencing data, the reference genome, and the truth sets used in this study, together with the locations of the software, models, and skill.

#### S5.1 GIAB HG003 (NA24149) accuracy benchmark

- GIAB FTP root: `https://ftp-trace.ncbi.nlm.nih.gov/ReferenceSamples/giab/`
- Truth set v4.2.1 (GRCh38): `https://ftp-trace.ncbi.nlm.nih.gov/ReferenceSamples/giab/release/AshkenazimTrio/HG003_NA24149_father/NISTv4.2.1/GRCh38/`
- Oxford Nanopore reads (AWS Open Data): `s3://ont-open-data/giab_2023.05/`
- Oxford Nanopore registry: `https://registry.opendata.aws/ont-open-data/`
- Oxford Nanopore data browser: `https://labs.epi2me.io/`
- PacBio HiFi and Illumina reads (GIAB AshkenazimTrio HG003): `https://ftp-trace.ncbi.nlm.nih.gov/ReferenceSamples/giab/data/AshkenazimTrio/HG003_NA24149_father/`

#### S5.2 1000 Genomes HG00136 and HG00150 (*APOE* task)

- HG00136 CRAM: `https://ftp.sra.ebi.ac.uk/vol1/run/ERR324/ERR3240142/HG00136.final.cram`
- HG00150 CRAM: `https://ftp.sra.ebi.ac.uk/vol1/run/ERR324/ERR3240154/HG00150.final.cram`
- Sample index: `https://ftp.1000genomes.ebi.ac.uk/vol1/ftp/data_collections/1000G_2504_high_coverage/1000G_2504_high_coverage.sequence.index`
- NYGC phased panel (truth): `https://ftp.1000genomes.ebi.ac.uk/vol1/ftp/data_collections/1000G_2504_high_coverage/working/20220422_3202_phased_SNV_INDEL_SV/`
- IGSR sample page (HG00136): `https://www.internationalgenome.org/data-portal/sample/HG00136`

#### S5.3 Reference genome (GRCh38)

- GRCh38DH (`GRCh38_full_analysis_set_plus_decoy_hla.fa`): `https://ftp.1000genomes.ebi.ac.uk/vol1/ftp/technical/reference/GRCh38_reference_genome/`
- GRCh38 no-alt analysis set: `https://ftp-trace.ncbi.nlm.nih.gov/1000genomes/ftp/technical/reference/GRCh38_reference_genome/`

#### S5.4 Models, tools, and skill

- Clair3: `https://github.com/HKU-BAL/Clair3`
- Clair3 model zoo: `http://www.bio8.cs.hku.hk/clair3/clair3_models/`
- Stock Clair3 Docker image: `hkubal/clair3:v2.0.1_gpu`
- hap.py: `https://github.com/Illumina/hap.py`

- Clair-skills: `https://github.com/HKU-BAL/Clair-skills`
